# A single-cell cytokine dictionary of human peripheral blood

**DOI:** 10.64898/2025.12.12.693897

**Authors:** Lukas Oesinghaus, Sören Becker, Larsen Vornholz, Efthymia Papalexi, Joey Pangallo, Amir Ali Moinfar, Jenni Liu, Alyssa La Fleur, Maiia Shulman, Simone Marrujo, Bryan Hariadi, Crina Curca, Alexa Suyama, Maria Nigos, Oliver Sanderson, Hoai Nguyen, Vuong K. Tran, Ajay A. Sapre, Olivia Kaplan, Sarah Schroeder, Alec Salvino, Guillermo Gallareta-Olivares, Ryan Koehler, Gary Geiss, Alexander B. Rosenberg, Charles M. Roco, Georg Seelig, Fabian J. Theis

**Affiliations:** Department of Electrical and Computer Engineering, University of Washington, Seattle, USA; Botnar Institute for Immune Engineering, Basel, Switzerland; Institute of Computational Biology, Helmholtz Center Munich, Munich, Germany; School of Computing, Information and Technology, Technical University of Munich, Munich, Germany; Parse Biosciences, Seattle, USA; Paul G. Allen School of Computer Science and Engineering, University of Washington, Seattle, USA; TUM School of Life Sciences Weihenstephan, Technical University of Munich, Munich, Germany

**Author notes:** These authors contributed equally to this work (Lukas Oesinghaus, Sören Becker, Larsen Vornholz, Efthymia Papalexi).

## Abstract

Cytokines orchestrate immune responses, yet we still lack a comprehensive understanding of their specific effects across human immune cells due to their pleiotropy, context dependence and extensive functional redundancy. Here, we present a Human Cytokine Dictionary, created from high-resolution single-cell transcriptomes of 9,697,974 human peripheral blood mononuclear cells (PBMC) from 12 donors stimulated *in vitro* with 90 different cytokines. We describe donor-specific response variation and uncover robust consensus cytokine signatures across individuals. We then delineate similarities between cytokine response profiles, and derive cytokine-induced immune programs that organize responsive genes into data-driven, biologically interpretable functional modules. By integrating cell type-specific responses with expression of cytokines, we infer higher-order cell-to-cell and cytokine-to-cytokine communication networks exemplified by an IL-32-β-initiated signaling cascade, which rewires myeloid programs by inducing neutrophil-recruiting factors while suppressing Th1-responses and promoting IL-10-family cytokines. Finally, we show how the Human Cytokine Dictionary enables the interpretation of cytokine-driven immune responses in other studies and disease contexts, including systemic lupus erythematosus, multiple sclerosis, and non-small cell lung carcinoma. Together, the Human Cytokine Dictionary constitutes the first comprehensive cell type-resolved transcriptional screen of human cytokine responses and provides an essential open-access, easy-to-use community resource with accompanying software package to advance our understanding of cytokine biology in human disease and guide therapeutic discovery.

## Main

Cytokines are small signaling proteins that regulate immune cell differentiation and activity, coordinating systemic responses from inflammation to tissue homeostasis^1^. Clinically, cytokine activity is modulated using blocking antibodies, small molecule inhibitors, engineered cytokines, or cytokine-based immunotherapies to suppress or enhance immune responses in inflammatory disorders and cancer2–4. As central players in immune biology, cytokines and their interactions with diverse cell types have been studied for decades. However, substantial gaps remain in our understanding of these interactions for many cell type-cytokine pairs. Moreover, the heterogeneity of experimental systems as well as batch effects make it difficult to directly compare cytokine effects across studies and thus reliably decipher a universal cytokine code^5^. With single-cell technologies, it is now possible to record detailed responses to many perturbations at once, enabling direct comparison across many cytokine-cell type pairs^6−^8. Building on these developments, we here use a combinatorial barcoding single-cell RNA sequencing approach^9^ (Parse Biosciences) to generate a dataset encompassing millions of cells and hundreds of cell type-cytokine interactions, thereby taking a step towards a systematic catalog of human cytokine responses.

### Creating the Human Cytokine Dictionary

To construct the Human Cytokine Dictionary, we obtained PBMCs from 12 healthy donors (6 males, 6 females, ages ranging from 34 to 75, **Supplementary Table 1**) from Seattle Bloodworks and treated them *in vitro* with 90 individual cytokines or PBS for 24h, then sequenced using split-pool barcoding (Parse Biosciences GigaLab)^9^ (**Fig. 1a**). We selected cytokines representing the biologically and clinically most relevant signaling pathways and major cytokine families, including IL-1, common γ chain, IL-4/IL-13, common β chain, IL-6, IL-12, IL-10, IL-17, interferon (IFN), tumor necrosis factor (TNF), complement, growth factors, transforming growth factor (TGF)-β, and others^8^, with concentrations chosen towards the upper range typically used in *in vitro* studies (**Supplementary Table 2**). We retained 9,697,974 cells after applying QC thresholds and annotated clusters based on expression of marker genes^10,11^ (Methods, **Fig. S1, Fig. S2**). This resulted in 16 distinct major cell types, namely B cells (naive + intermediate), CD4 T cells (naive + memory), CD8 T cells (naive + memory), regulatory T cells (Treg), mucosal-associated invariant T (MAIT) cells, natural killer T (NKT) cells, CD56high natural killer (NK) cells (NK CD56hi), CD56low NK cells (NK CD56low), conventional dendritic cells (cDC), CD14 monocytes (CD14 Mono), CD16 monocytes (CD16 Mono), hematopoietic stem and progenitor cells (HSPC), plasmacytoid dendritic cells (pDC), innate lymphoid cells (ILC), granulocytes, and plasmablasts (**Fig. 1b**). The last four (pDC, ILC, granulocytes, plasmablasts) were discarded from downstream gene expression analysis due to a low median abundance of <10 cells per condition and donor (**Fig. S3a-b**).

**Fig. 1.**
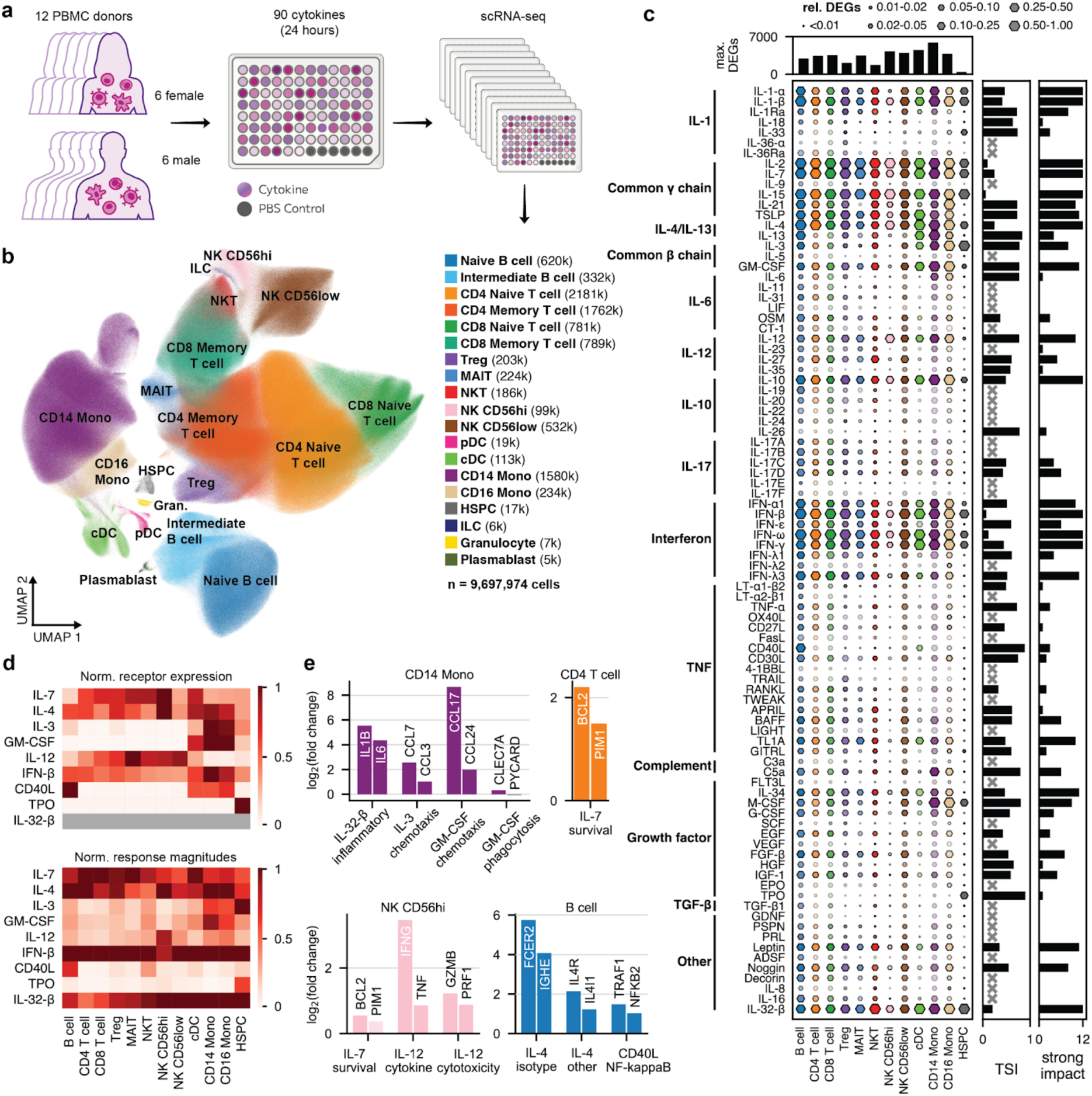
The human cytokine dictionary. **a,** Overview of the experimental procedure for generating the dictionary. **b,** UMAP of cell type populations across all conditions. The legend indicates total cell counts for each population. **c,** Overview of the response to different cytokines. The box size is proportional fraction of DEGs relative to the maximum number of DEGs for that cell type (cf. bar plot on top). The color saturation denotes the transcriptomic response magnitude as defined in the main text. The first bar plot on the right shows the tissue specificity index (TSI) of the response magnitude, a measure of whether a cytokine response is highly specific to a single cell type (TSI=1) or has the same strength across tissues (TSI=0). Gray crosses indicate that no TSI was calculated due to an absence of a strong response in any cell type. The second bar plot shows the count of cell types per cytokine in which the response magnitude is larger than a threshold value (“strong impact”). **d,** Normalized receptor expression (top) and response magnitudes (bottom) for a subset of cytokines. **e,** Fold changes for individual cytokines, cell types, and genes. Genes are grouped by function.

Few perturbations result in major shifts in cell type abundance or composition: In differential abundance tests (per donor Wilcoxon signed rank), the most abundant cell types exhibit no statistically significant (adjusted p-value (padj)<0.05) fold changes (|log2FC|>0.5) (**Fig. S3c**). We do observe significant changes in less abundant cell types such as a >2-fold decrease in NKT cells upon interferon perturbation or a >2-fold decrease in CD16 Mono upon IL-4 stimulation. Compositional modeling (scCODA)^12^, using CD4 T cells as reference, similarly identifies no large compositional changes among major cell types and largely recapitulates the perturbation-specific changes in rarer subsets (**Fig. S3d**).

We aggregated expression profiles by donor, cell type, and cytokine perturbation to robustly^13^ identify donor-consensus differentially expressed genes (DEGs) and log2FCs (cytokine treated vs PBS) (**Fig. 1c**, **Fig. S4**, methods)^14^. Subpopulations of B cells, CD4 T cells, and CD8 T cells were merged for downstream analysis due to a high similarity in their log2FCs (**Fig. S5**). The number of DEGs (padj<0.05, |log2FC|>0.25) varied strongly by cytokine and cell type (**Fig. 1c, Fig. S6a-c**). IL-1, common γ chain interleukins, IL-4, IL-10, interferons, and IL-32-β have the largest number of DEGs across cell types. Most other cytokines elicit fewer DEGs and responses that are limited to a single or few cell types (**Fig. 1c**).

To quantify the global transcriptional response to a cytokine by a more continuous measure independent of specific adjusted p-value and log2FC thresholds, we defined a response magnitude based on Euclidean distance of log2 expression vectors between each perturbation and the PBS controls and log2FCs weighted by p-values (Methods, **Fig. S6d**, color saturation in **Fig. 1c**, **Supplementary Table 3**). Response magnitude variation across cell types is captured by a tissue specificity index (TSI)^15^, which is highest for TPO (0.93, only affects HSPCs) and lowest for IL-15 (0.07, affects all cell types) (bar plot in **Fig. 1c**, **Fig. S6e**). We further define a per cell type threshold value for the response magnitude that classifies whether a cytokine has a strong impact on a particular cell type (bar plot in **Fig. 1c**, **Fig. S6f, Fig. S7**, **Supplementary Table 4**). For example, IL-4 has a strong impact on all cell types while CD40L has a strong impact only on B cells and cDCs.

### Cytokine-induced responses include expected marker genes and match receptor expression

We compared the cell type-specific expression of cytokine receptors in the baseline (PBS) state to the response magnitude of their associated cytokine responses (**Fig. 1d**, **Fig. S8a**, **Supplementary Table 5**). As expected, receptor expression and responses for IL-4, IL-7, and IL-12 are distributed widely across cell types, albeit with a much stronger response in NK CD56hi for the latter. In contrast, receptor expression and responses for CD40L, TPO, and GM-CSF are mostly constrained to B cells, HSPCs, and monocytes, respectively (**Fig. 1d**)^16–23^. Target gene expression downstream of cytokine signals also matches responses expected from the literature (**Fig. 1e**): In B cells, CD40L induces NF-κB signaling^22^ while IL-4 upregulates *IGHE* (isotype switch)^20^, its own receptor (*IL4R*), and *IL4I1*. IL-7 upregulates survival genes in CD4 T cells and NK CD56hi^21^, while IL-12 upregulates the production of inflammatory cytokines and cytotoxicity genes also in NK CD56hi^23^.

Across all cytokines and cell types, receptor expression and response magnitude are correlated (r≈0.4) and responses generally require a minimum baseline receptor expression (≈8 counts per million (cpm)) (**Fig. S8b**). However, high receptor transcript abundance does not always translate into strong responses. This is most evident for atypical ligands such as Decorin, Resistin (ADSF) or LT-α2β1, whose activities rely on non-classical or low-affinity receptor interactions^24–26^, death ligands (TRAIL, FasL) to which resting immune cells are resistant until activated^27,28^, and TGF-β1 (**Fig. S8c**). A few other cytokines, e.g., IL-2 and IL-15 in CD14 monocytes, induce strong responses despite lacking receptor expression. This is likely due to secondary cytokine release, as discussed later. More generally, mRNA expression levels do not always predict receptor function, because protein abundance, receptor stability, and post-translational modifications can modulate signaling activity.

### Cytokine response profiles are consistent across donors despite inherent heterogeneity

We next investigated donor heterogeneity (**Fig. 2a**). Cell type composition varies by donor but aligns with expectations for PBMCs^10^ (**Fig. 2b**). To understand gene expression heterogeneity before cytokine perturbation, we calculated a baseline log2FC as the ratio of cpm for a given donor in the PBS control to the median cpm across all donors in the PBS controls (**Fig. S9a**). A subset of donors (D1, D3, D4, D10) is highly correlated in its baseline log2FC across cell types (r=0.61±0.10, **Fig. 2c-d**). These donors have high baseline expression of interferon-stimulated genes (ISG)^29^ (e.g., *IFIT1-3)* in CD4 T cells (**Fig. 2e**, **Fig. S9b**). In the UMAP visualization of CD4 T cells, PBS-treated cells from these donors cluster with IFN-β-treated CD4 T cells of all other donors (**Fig. 2f**). Together these observations indicate a higher baseline interferon signaling in this group (“interferon group”). Interestingly, three out of four members of this group are the oldest donors in our set.

**Fig. 2.**
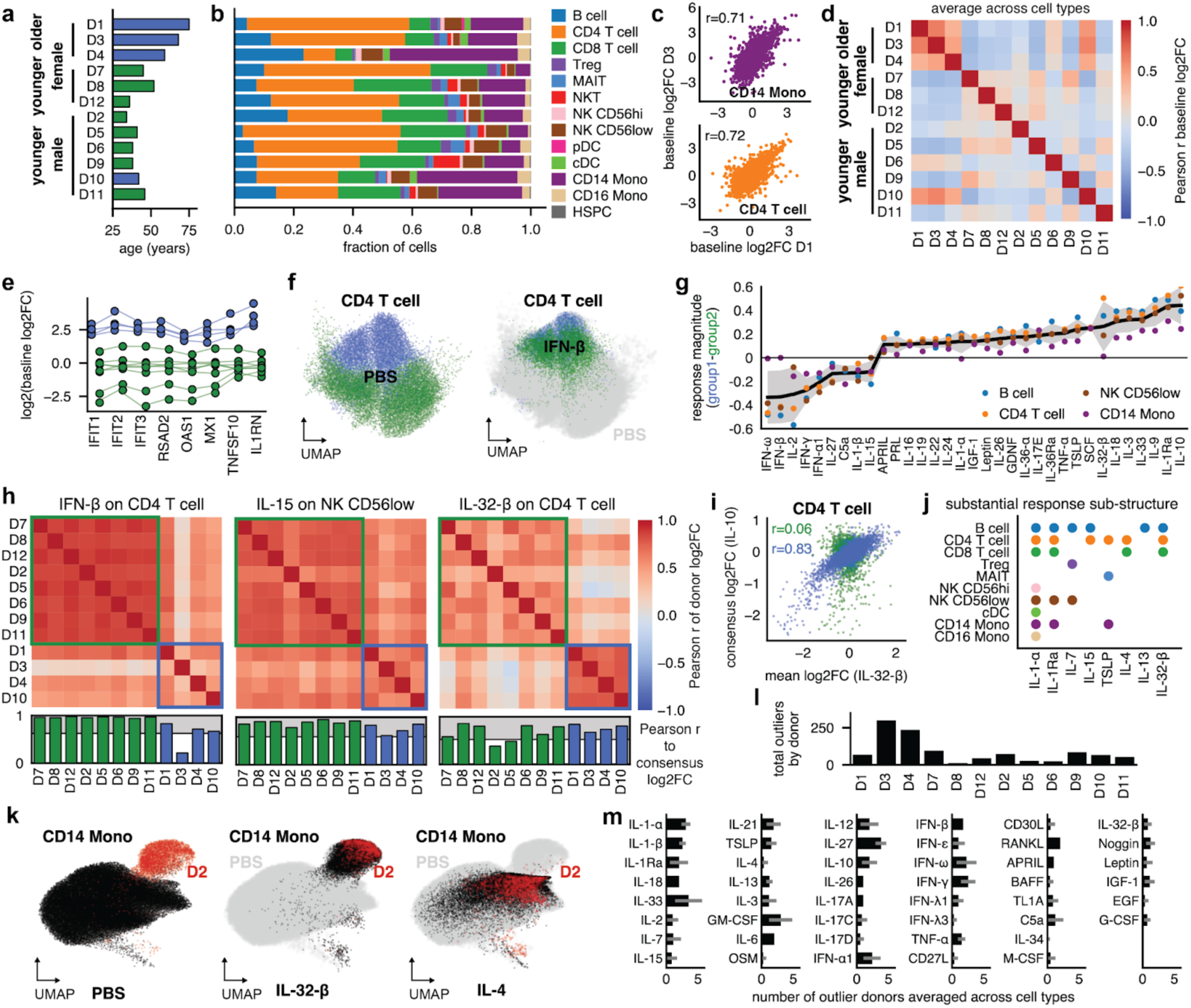
Donor variability analysis reveals subpopulations with distinct cytokine responses. **a,** Age and sex distribution of donors. **b,** Cell type fractions across cytokine perturbations by donor **c.** Comparison of baseline log2FC for donors D1 and D3 in CD14 Mono (top) and CD4 T cells (bottom). **d,** Correlations of the baseline log2FC between donors averaged across all cell types. **e,** Baseline log2FC of interferon response genes for two groups of donors identified in (d). **f,** UMAP of CD4 T cells of the two groups for PBS and IFN-β. **g,** Difference in the mean response magnitude by cytokine averaged across four cell types for the two groups. The gray area shows the standard deviation. **h,** Correlation of donor-specific log2FCs for different cell types and cytokines. The bottom barplot shows the correlation to the donor-consensus log2FCs. Responses below the shaded grey region (see Methods) are considered outliers. **i,** Donor-consensus log2FC for IL-10 versus group mean per-donor log2FC for IL-32-β in CD4 T cells. **j,** Overview of cytokines and cell types with substantial substructure in donor responses. **k,** UMAP of CD14 Mono for PBS, IL-32-β, and IL-4. Donor D2 cells are highlighted in red. **l,** Total outlier response count by donor. **m,** Average number of outlier responses across all cell types by cytokine.

Next, we asked whether this baseline heterogeneity imperils the determination of a meaningful consensus response to cytokine perturbations. Generally, donor response magnitudes correlated well for most cell types (r=0.55±0.23), suggesting that consistent responses are observed even for donors with different baselines. Still, pairwise correlations between donors were higher in the non-interferon group (r=0.69±0.19) than in the interferon group (r=0.45±0.23) or across groups (r=0.45±0.20) (**Fig. S9c-e**). For most cytokines, mean response magnitudes are similar for the two groups (**Fig. 2g**). Notable exceptions include, for example, stronger responses to IL-10, IL-1Ra, and IL-32-β and weaker responses to interferons and IL-2 in CD4 T cells for donors in the interferon group.

An individual donor’s response to a specific cytokine sometimes differed substantially from the consensus response calculated across all donors. For example, donor 3 log2FCs and donor-consensus log2FCs for the response to IFN-β are only weakly correlated, making donor 3 an outlier. Donor 3 stands out even among members of the interferon group which collectively are less correlated with the consensus (**Fig. 2h**, left, **Fig. S9f-g**). For other cytokines and cell types, e.g., IL-15 in NK CD56low, the baseline state, while still visible in the response, has a relatively weak impact and all individual donor responses are well correlated with the overall response (**Fig. 2h**, middle, **Fig. S9f-g**).

Notably, the response to IL-32-β in CD4 T cells, although not in CD14 Mono, depended heavily on the donor interferon state (**Fig. 2h**, **Fig. S9f**). There, the two donor groups exhibited responses that were internally consistent within groups (r=0.60±0.08, r=0.76±0.03) but divergent between groups (r=0.29±0.19). To understand how these responses differ biologically, we calculated correlations to the consensus log2FCs of all other cytokines. The interferon group response showed a strong correlation with the consensus IL-10 log2FCs that is absent in the non-interferon group (**Fig. 2i**), indicating an anti-inflammatory response to IL-32-β for pre-inflamed cells only. Although there are other examples of such response substructure (IL-1-α or IL-1Ra in CD4 T cells, **Fig. S10a-b**), which we define as the presence of at least two groups of donors with internally consistent but externally divergent responses (**Fig. S10c**), they are rare (**Fig. 2j**) and overwhelmingly focused on only donors 3 and 4 of the interferon group (**Fig. S10d**).

Baseline interferon-signaling was not the only source of donor heterogeneity. For example, the donor 2 CD14 Mono population was almost completely separate from those of all other donors in the UMAP of the PBS controls. However, stimulation with IL-32-β shifted all other donors’ monocytes to the same spot in the UMAP as the donor 2 baseline (**Fig. 2k**), while donor 2 monocytes showed only weak responses to IL-32-β (**Fig. S11a**), suggesting strong pre-existing IL-32-β-like signaling for donor 2. Indeed, the donor-consensus log2FCs for IL-32-β were highly correlated (r>0.91 for CD14 Mono) with the donor 2 baseline log2FCs across cell types but uncorrelated for other donors (**Fig. S11b-c**). Conversely, IL-4 treatment shifted donor 2 monocytes back to the space occupied by the other donors’ baseline (**Fig. 2k, Fig. Slid**). IL-32-β treatment of CD14 Mono in our screen upregulated *CD14* and downregulated monocyte-derived dendritic cell (moDC) markers^30^ (**Fig. Slle-f**). The overall state of macrophage differentiation markers^31^ was more consistent with a unique monocyte activation state rather than macrophage differentiation^32^ (**Fig. Slle-f**). Despite this difference in the baseline state, donor 2 was usually not an outlier in responses to cytokines other than IL-32-β (**Fig. 2l**).

Overall, outlier responses were well distributed across cell types (**Fig. S10e**) and most cytokine-cell type combinations had relatively few outliers (**Fig. 2m**, mean=1.2±0.8 in cell types with strong impact). While a few cases exhibited a clear donor-specific substructure, the overall patterns were thus robust enough that a single set of log2FCs per cell type captures a meaningful consensus response.

### Comparison across mouse and human datasets identifies shared and divergent cytokine responses

We next examined how cytokine-induced responses in human immune cells relate to those previously reported in mice. Of the 90 cytokines profiled in our study, 81 mouse homologues were also used in the cytokine response screen generated by Cui et al.^8^. To ensure that the comparison reflects genuine perturbation effects, we focused on strong cytokine responses. These were defined for our dataset as in **Fig. 1c**, and the same thresholding criterion was applied to the response magnitudes reported by Cui et al. for the mouse. We find an intersection of 8–20 cytokines per cell type that elicit strong responses in both datasets (**Fig. 3a**).

**Fig. 3.**
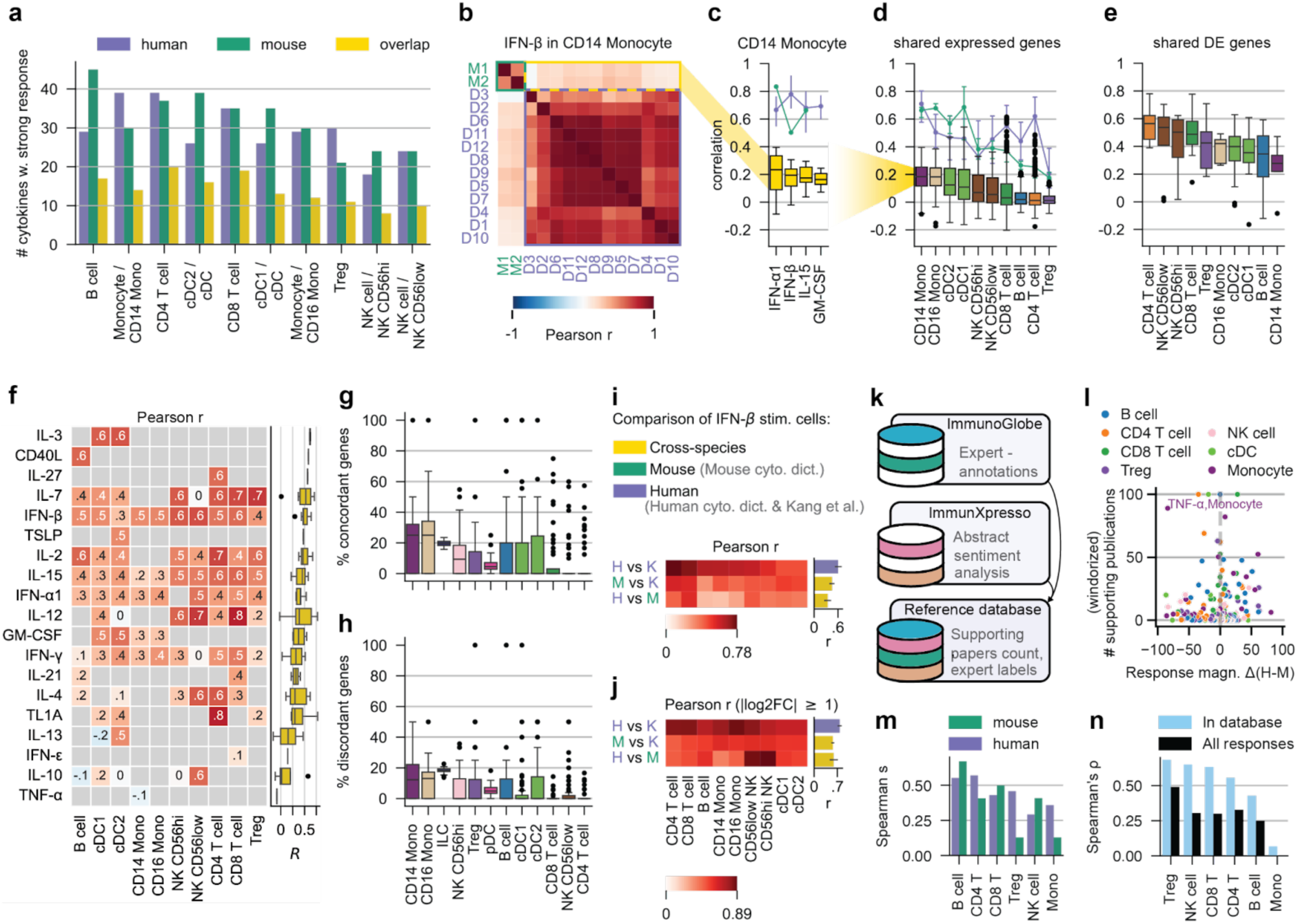
Comparison between human and mouse cytokine responses highlights overlapping yet distinct gene signatures. **a,** Per cell type, number of cytokines that induce a strong response in humans, mice or in both species (overlap). **b,** Pearson correlation of log2FC (IFN-β vs PBS) between all human donors (D1-12) and mouse replicates (M1-2) in CD14 Mono. **c,** Distribution of Pearson correlations across all cross-species sample pairs, per cytokine in CD14 Mono. The purple line shows correlations between all human donor pairs; green line shows correlations between mouse replicates. The point corresponds to the median, the error bars indicate the quartile range. **d,** Aggregation of Pearson correlations including all strong-response cytokines in a given cell type. As in c, boxplots represent cross-species correlations, purple and green curves within-species correlations. **e,** Same as d, but restricted genes that are differentially expressed in both human and mouse. **f,** Pearson correlations of average human and average mouse log2FCs for strong-response cytokines per cell type; restricted to cell type-cytokine combinations with at least 5 genes. CD14 and CD16 monocyte labels apply only to the human dataset, as the mouse dataset does not distinguish monocyte subtypes; all comparisons are therefore made against the single monocyte population in mouse. Conversely, the mouse dataset separates cDCs into cDC1 and cDC2, whereas the human dataset contains a single cDC population. **g,** Percentage of concordant genes, defined as the number of genes with log2FC>1 in both datasets plus those with log2FC<–1 in both datasets, divided by the number of genes with \log2FC\>1 in the human dataset. **h,** Percentage of discordant genes, defined as number of genes with log2FC>1 in human but log2FC<-1 in mouse or vice versa, divided by the number of genes with \log2FC\>1 in the human dataset. **i,** Pearson correlations of log2FC for IFN-β-stimulated cells across studies. Cell type labels are shared with **j**. **j,** Same as **i** but only considering genes with \log2FC>1\ in both datasets of the comparison. **k,** Integration of the ImmunoGlobe and ImmuneXpresso databases into a unified reference database of cytokine-immune cell effect annotations, with indications of number of supporting publications. **l,** Difference between human and mouse response magnitudes for the indicated cell types vs the number of supporting publications from the reference database. **m,** Spearman’s p of the correlation between response magnitude and number of supporting reference publications, for cell types with >10 cytokine effect annotations. **n,** Spearman’s p of human vs. mouse response magnitudes per cell type, for all shared cell type-cytokine relationships and subsetted to those which have supporting publications in the reference database.

To assess response similarity, we calculated Pearson correlations of gene-wise log2FCs for each cell type-cytokine condition. To ensure robustness, we only included samples (human donors or mouse replicates) containing at least 20 cells and genes with sufficiently high expression levels (≥ 20 cpm) in either PBS- or cytokine-treated cells in both species. Consistent with our previous analysis of donor variability, correlations of IFN-β-stimulated CD14 Mono across human donors were high (median r=0.78), as were correlations among the two mouse replicates (r=0.5) (**Fig. 3b**). In contrast, cross-species correlations were substantially lower (median r=0.19), a pattern that held across cytokines and cell types (**Fig. 3c,d**). Restricting the analysis to genes that are differentially expressed in both species increased correlations markedly across most sample pairs and cell types (**Fig. 3e**).

To finally compare average human and mouse responses, we created pseudobulks per condition within each species. Log2FC correlations across differentially expressed genes were generally moderate to high, but notable exceptions included IL-13 in cDCs, TNF-α in CD14 Mono, and IL-10 in B cells, all of which showed mildly negative correlations (**Fig. 3f**).

Restricting correlation analyses to shared differentially expressed genes imposes a stringent constraint and relies on DE results that may be difficult to compare across datasets, e.g., due to differences in statistical power arising from unequal cell numbers per condition. We therefore asked how many genes meaningfully contribute to these correlations. To address this, we defined *strongly regulated concordant genes* as those with log2FC > 1 in both datasets or log2FC < –1 in both datasets, and *discordant genes* as those exhibiting strong but opposing fold changes (log2FC >1 in human but <–1 in mouse, or vice versa). This analysis revealed that only a relatively small fraction of genes (mean=11.3%) display strongly concordant regulation (**Fig. 3g**) and that a non-negligible amount (mean=6.9%) of genes showed strong discordant responses (**Fig. 3h**).

Although these results indicate that there are substantial differences between the mouse and human cytokine datasets, these cannot be readily ascribed to fundamental interspecies divergence, given the differences in experimental design - most notably *in vitro* versus *in vivo* conditions, stimulation duration (4h versus 24h), and cytokine dosage. We therefore compared cytokine responses to an independent dataset of *in vitro* IFN-β-stimulated human PBMCs with a readout time similar to the mouse dataset (6h)^33^. Although correlations vary by cell type, they are consistently higher in the within-human comparison (mean r=0.61) than in comparisons across species (mean r=0.45 (mouse cyto. dict. vs Kang et al.) and r=0.34 (human vs mouse cyto. dict.)) (**Fig. 3i**) both when considering all shared expressed genes (cpm ≥ 20 in stimulated or control cells) and when restricting to genes that show large fold changes (|log2FC| > 1) in both datasets (within-human mean r=0.77; across species mean r=0.57 (mouse cyto. dict. vs Kang et al.) and r=0.59 (human vs mouse cyto. dict); **Fig. 3j**).

To extend our comparison beyond the study by Kang et al., we collected a reference database of cytokine-immune cell interactions by integrating data from the ImmunoGlobe^34^ and immuneXpresso^35^ databases. ImmunoGlobe consists of expert-curated relationships summarizing Janeway’s Immunobiology^36^; immuneXpresso was constructed by text-mining PubMed abstracts. After merging entries based on Cell Ontology identifiers^37^ (**Fig. S12a**), our reference database lists well-studied cytokine-cell interactions, including the number of supporting publications and an indication of expert review status (**Fig. 3k, Fig. S12b**). Based on the observation that stronger responses are supported by a larger number of publications (**Fig. S12c,** Spearman’s ρ=0.42 for human, ρ=0.33 for mouse), we can use this database to compare and verify differences in response magnitudes between the human and mouse datasets. Interestingly, large differences in response magnitudes occur in cell-cytokine pairs with fewer supporting publications whereas conditions that are supported by a large number of publications tend to show relatively small differences in response magnitudes, with the only clear outlier being TNF-α in monocytes (cf. **Fig. 3l**). Experimentally determined response magnitudes and database-derived numbers of publications remain fairly well correlated when the analysis is performed separately for each major cell type (**Fig. 3m, Fig. S12d**). However, we do see a difference in the Treg and monocyte response Spearman ρ, with the humans showing higher correlations between response magnitude and number of reported publications. Finally, we can directly compare response magnitudes for shared cytokine-cell pairs between the mouse and human cytokine dictionaries. We find that most cell types are decently correlated (Spearman’s ρ ∼0.25-0.5) with the exception of monocytes (**Fig. 3n**). These correlations increase if we only consider cell-cytokine pairs that also occur in the reference database, again indicating that strong changes tend to be more correlated.

### Cell type-specific production and response patterns define a detailed network of cellular communication via cytokines

To unravel how cytokines orchestrate multicellular responses, we assessed how cytokine signaling connects the cell types in our dataset. For each of 184 cytokines from a manually curated list (**Supplementary Table 6**), we first averaged the expression values of its encoding genes across all stimulation conditions, providing a stimulation-agnostic, cell type-specific view of cytokine production (**Fig. 4a**, **Fig. S13a**). This analysis revealed that production for most cytokines is strongly cell type-specific. For example, IL-10 is strongly produced by CD14 Mono, CD40L by CD4 T cells and MAITs, and IL-32 by all T cell subtypes but in particular by Tregs. When aggregating over all cytokines that belong to a specific family (e.g., all chemokines) and counting the number of produced cytokines (**Fig. 4a**, average cpm>4), we observe that certain families are preferentially produced by specific cell types. For example, monocytes and cDCs dominate chemokine production, albeit far more strongly for CXCL than CCL chemokines (**Fig. S13b**). Results from this stimulation agnostic analysis are also supported by our literature-derived reference database (**Fig. S14**).

**Fig. 4.**
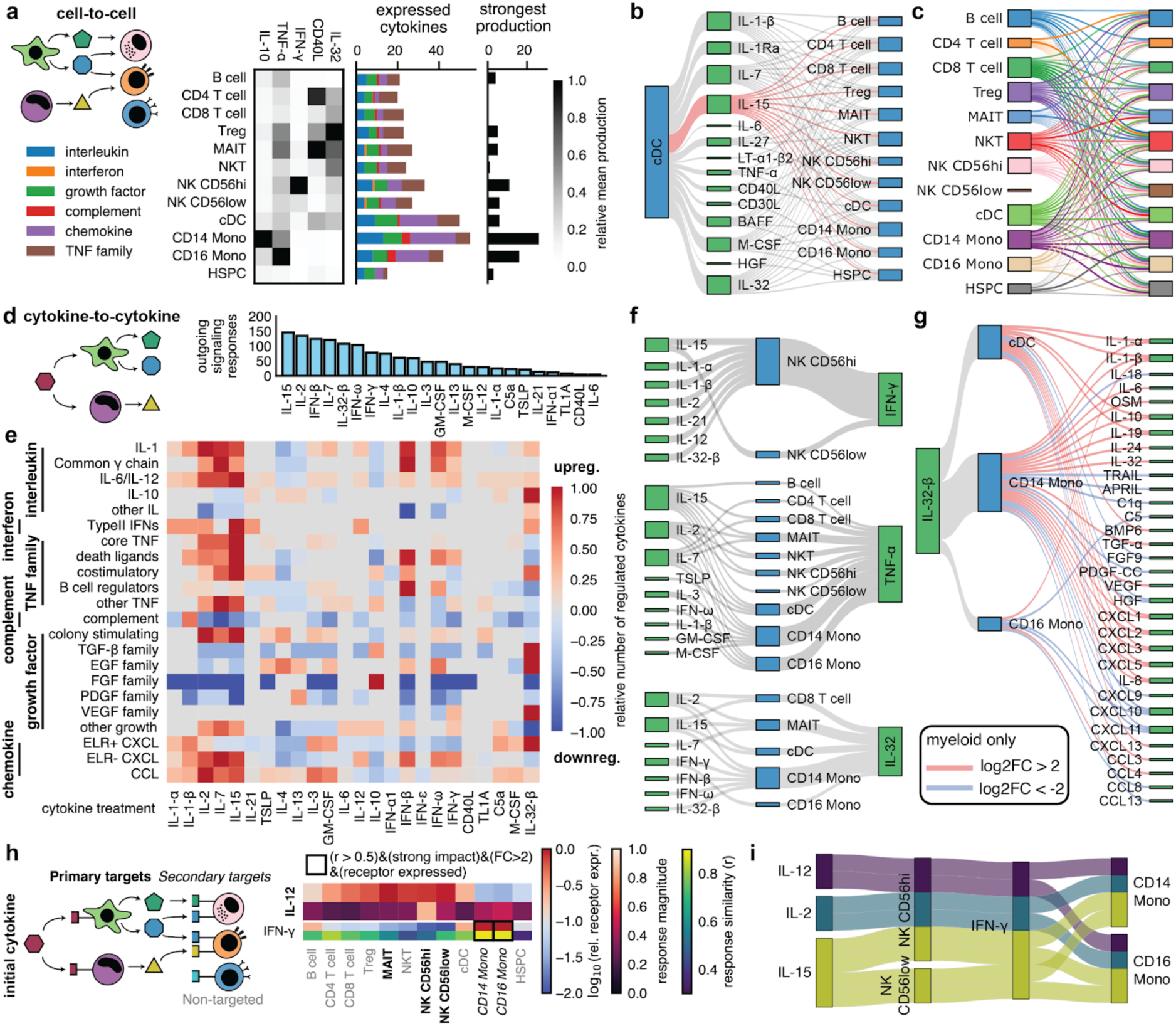
The Human Cytokine Dictionary unveils cell type-specific cytokine signaling networks. **a,** Cytokine production by different cell types and how this connects them to other cell types (sketch). The left heatmap shows the relative levels of production for five different example cytokines. The middle bargraph shows the number of expressed cytokines per cytokine family per cell type. The right bar graph shows the number of times a cytokine has its maximum expression level in a given cell type. **b,** Cytokines secreted by cDCs cause strong responses in other cell types. The height of each box is proportional to the number of connections through that box. IL-15 connections are highlighted. **c,** Strength of cell-to-cell signaling given the cytokine secretion profile. The transparency of connections shows the overall connection strength, normalized to the maximum within that cell type and subset to connections with an interaction score>0.25. **d,** The number of times a given cytokine significantly regulates another cytokine summed across all cell types. **e,** Up- and downregulation of different cytokine families by cytokine treatments. The figure sums regulation in the different cell types and then divides by the maximum value for that family to get a relative number. When different cytokines of a given family are both up- and downregulated, values are subtracted. **f,** Upregulation (paαj<0.05, logFC>1) of a specific cytokine by other cytokines. **g,** Cytokines up- or downregulated by IL-32-β (paαj<0.05, |log2FC|>2) in myeloid cells. **h,** Secondary responses due to cytokines secreted in response to the initial cytokine change the response profile. Low receptor expression and high response magnitude for IL-12 along with a high response similarity implies IFN-γ as likely responsible for the response in monocytes. **i.** IL-2, IL-12, and IL-15 all signal to secondary cells via IFN-γ released by NK cells.

Next, we studied cell-cell communication via specific cytokines. A cell-cell signaling connection was considered present when a cytokine was expressed in a given cell type and had a strong impact on a target cell type in our screen. All cell types signaled to other cell types by a variety of cytokines (**Fig. S15**, **Supplementary Table 7**), such as cDCs signaling to all other cell types via IL-15 (**Fig. 4b**)^38^.

For a more quantitative view, we defined sender strength as the relative expression for a cytokine compared to other cell types, receiver sensitivity as the response magnitude for that cytokine, and their product as the interaction score. We used interaction scores to identify the central signaling cytokines for all cell type pairs in our screen (**Fig. S16**), yielding IL-32 as a potential dominant signaling molecule connecting all T cell subtypes to all other cell types^39^. Adding the interaction scores between two cell types yields an overall cell-cell connection strength (**Fig. 4c**). Notably, while the signaling from NK CD56low is too weak to generate even a single connection, NK CD56hi signals to nearly all other cell types^40^.

### Cytokine responses enable mapping of cytokine signaling cascades via immune cells

Complementing the cell type-centric communication analysis, we next performed a cytokine-focused analysis by initially summing interactions across cell types. We find a wide range of signaling activities with the most prolific cytokine, IL-15, activating or repressing the expression of other cytokines 146 times across our cell types (**Fig. 4d-e**, **Fig. S17**, padj<0.05 and |log2FC|>1). We found some expected patterns: For example interferons and common γ chain cytokines upregulate ELR-CXCL chemokines^41^, while IL-10, IL-4, and IL-13 downregulate them (**Fig. 4e**).

Next, we looked at individual up- or downregulated cytokines for a specific stimulation and cell type (**Fig. S18-S21**, **Supplementary Table 8**). For example, IFN-γ was upregulated by various inflammatory cytokines in NK cells and TNF-α was broadly upregulated by inflammatory cytokines across cell types (**Fig. 4f**). IL-32 was upregulated by common γ chain cytokines and interferons in T cells and myeloid cells (**Fig. 4f**). In turn, IL-32-β shifted the chemokine profile in myeloid cells (monocytes, cDCs) effecting a strong shift from Th1/interferon-like recruitment (downregulation of CXCL9, CXCL10, CXCL11, and IL-18; median log2FC∼-3.8) to neutrophil recruitment^42^ (upregulation of CXCL1, CXCL2, CXCL3, CXCL5, and CXCL8 as well as IL-1-α and IL-1-β; median log2FC∼5) (**Fig. 4g**). IL-32-β is also the only cytokine that strongly upregulated the IL-10 family in myeloid cells (IL-10, IL-19, IL-24)^43^, a response that is surprisingly weaker in the interferon donor group (mean log2FC 1.2±0.4 vs mean log2FC 2.5±1.1 for IL-10 in CD14 Mono, cf. **Fig. 2h**), while Th1-attractant chemokine downregulation is stronger in the interferon group (**Fig. S22**). Taken together, IL-32-β is a highly donor context-specific cytokine that converts monocytes from an initial antiviral/T cell-based response to a potent, neutrophil-driven, and highly inflammatory (but IL-10-family-based self-regulating) strategy consistent with aggressive, localized containment.

### Response similarity allows for the detection of potential secondary cytokine responses

The 24-hour stimulation allows PBMCs not only to respond directly to the primary cytokine but also to produce and secrete secondary (and higher-order) cytokines (**Fig. 4h**). Thus, the transcriptomic profiles measured at this time point capture the cumulative response to both the initial cytokine stimulus and the downstream cytokine signaling cascade. This signaling cascade explains cases in which cell types show strong transcriptional responses to a cytokine despite low expression of the corresponding primary cytokine receptor (<5 cpm, ‘secondary target’) (**Fig. S8c**, **Fig. S23a**). To identify which secondary cytokines might be causing these indirect responses, we applied two criteria: First, the secondary cytokine must be upregulated (log2FC>1, padj<0.05) after stimulation with the primary cytokine in cell types that clearly express the primary cytokine receptor (>16 cpm, ‘primary target’). This ensures that the cell is capable of responding directly to the primary cytokine and producing the secondary one. Second, the cell type that shows the secondary response should respond to the primary cytokine in a way that resembles its response when the secondary cytokine is directly applied to the cells. We measured this similarity by correlating log2FC values requiring a Pearson correlation above 0.5. For the effect of IL-12, IL-2, and IL-15 on monocytes only IFN-γ released by NK CD56hi fulfills these criteria (**Fig. 4h-i**, **Fig. S23b**)^44^.

### Classifying cytokine responses identifies consensus groups and unique effectors

The scale and design of our dataset allows unbiased discovery of functional similarity between cytokine responses without confounding batch effects, which previously allowed us to identify the IL-10-like character of IL-32-β responses in CD4 T cells for donors of the interferon group (cf. **Fig. 2i**). We quantified the similarity of responses between cytokines in the same cell type in terms of Pearson correlations of log2FC (**Fig. S24**, **Supplementary Table 9**). For example, in CD14 Mono, responses to common y chain interleukins IL-2 and IL-15 are highly similar (r=0.92), as are responses to IL-4 and IL-13 (r=0.94), which partially share receptors and signaling pathways^45^. Conversely, pro-inflammatory IL-15 and anti-inflammatory IL-10 have a strong negative correlation (r=-0.64) (**Fig. 5a**).

**Fig. 5.**
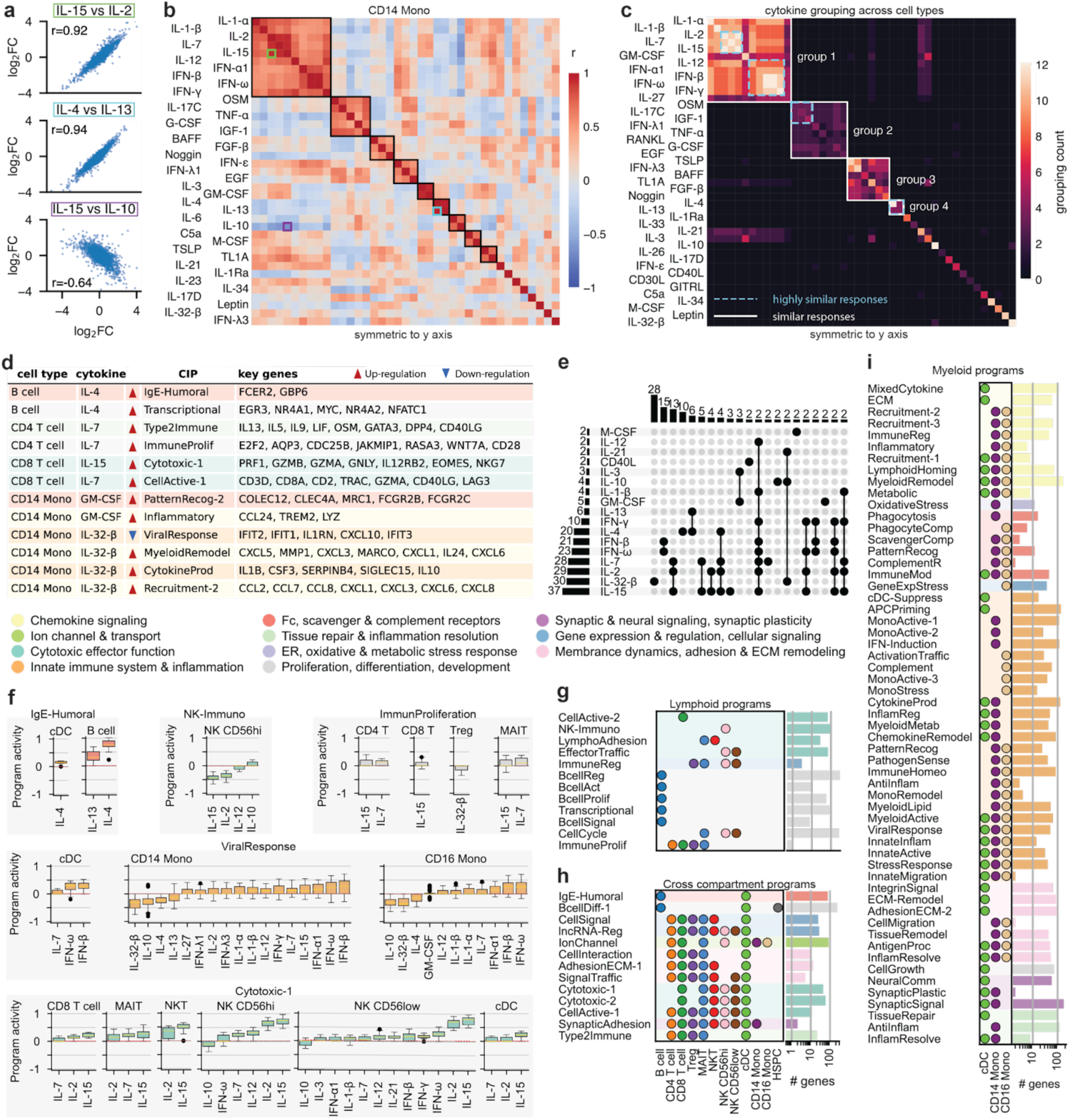
Data-driven deconvolution of cytokine responses defines cytokine-induced immune programs. **a,** Log2FCs between different cytokines in CD14 Mono. **b,** Pearson correlation between log2FCs for different cytokines in CD14 Mono. Cytokines were clustered based on their correlation patterns (black boxes). **c,** Clustering in individual cell types was used to derive consensus groups (white box). Highly similar cytokines (mean r>0.7) are marked with a dashed blue box. **d,** Table with CIPs and key genes that are associated with selected cytokines and cell types. **e,** Summary of how many programs one cytokine (left) or cytokine combinations (top) are part of (cytokines and combinations with one occurrence are omitteQ). **f**, CIP activity in relevant cell types as a function of activating cytokines. **g-i,** Significantly up- or downregulated cytokine-induced immune programs (CIPs) across cell types and the corresponding biological categories.

To systematically identify groups of cytokines eliciting similar responses in a given cell type, we clustered log2FC correlations using the Leiden algorithm (**Fig. 5b**, **Fig. S25-S27**). The largest group in all cell types consists of type 1/antiviral cytokines, most notably IL-1-α/β, common γ chain interleukins, and interferons. Other groupings are more cell type specific: For example, in monocytes GM-CSF forms a group with IL-3^18^ that is negatively correlated with interferons but in B and T cells it groups with type 1 cytokines (**Fig. 5b**, **Fig. S25**). Interestingly, IL-32-β, despite having strong effects in all cell types, does not cluster with other cytokines except for IL-10 in CD4 T cells and Tregs (**Fig. S25**).

To integrate information from all cell types, we counted the number of cell types in which a particular pair of cytokines was found in the same cluster, then used Leiden clustering on the count patterns to derive consensus similarity groups (**Fig. 5c**, **Supplementary Table 10-11**). We also defined subgroups of highly similar cytokines by a mean r>0.7. The largest group is type 1 cytokines, in which common γ chain interleukins and interferons form highly similar subgroups (group 1). IL-12 is also part of that latter subgroup, though in fewer cell types, and its high similarity to interferons in monocytes is a consequence of secondary responses to IFN-γ (cf. **Fig. 4i**). Two other large groups contain cytokines associated with inflammatory barrier defense and tissue protection (group 2) or cell growth, survival, and tissue development (group 3). IL-4 and IL-13 form a fourth r consensus group (group 4). However, while IL-4 has a strong impact on all cell types, IL-13 only significantly affects B cells, cDCs, and monocytes. Other cytokines such as IL-10 or IL-21 do not belong to any group because the responses they induce are similar to other response patterns in at best a small subset of the cell types they affect.

### Universal cytokine-induced immune programs (CIPs) reveal functional states of immune cells

To more comprehensively dissect the biology underlying cytokine-induced responses, we applied Disentangled Representation Variational Inference (DRVI)^46^ to identify cytokine induced gene programs (CIPs) - groups of genes that are jointly up- or downregulated upon cytokine stimulation. These programs represent the functional modules underlying cytokine-driven responses and may be unique to specific cell types or shared across multiple cell types. DRVI builds on the classical variational autoencoder framework but introduces key modifications to the decoder, enabling each latent dimension (CIP) to be linked to a distinct set of genes that is, to a large extent, exclusively modulated by that CIP (**Fig. S28**).

Applying DRVI to our dataset revealed 82 CIPs whose associated gene sets were manually inspected, annotated based on expert knowledge, and further summarized into 11 broad categories (**Methods, Supplementary Table 12-13**). For example, IL-4 stimulation in B cells induces a humoral effector program (IgE-Humoral) linked to IL-4-mediated class switching, as well as a transcriptional activation program (Transcriptional) that represents early signaling events, supporting differentiation toward antibody production^20^ (**Fig. 5d**). IL-32-β elicits a robust proinflammatory reprogramming in CD14 Mono, driving CIPs centered on myeloid activation (MyeloidRemodel), cytokine production (CytokineProd), and chemokine-driven neutrophil recruitment (Recruitment-2). Concomitantly, IL-32-β downregulates antiviral programs (ViralResponse), suggesting a shift from tissue-resident or antiviral states toward an acutely activated, recruitment-focused phenotype.

Since CIPs can be modulated by multiple cytokines, we next examined the overlap of programs across cytokines (**Fig. 5e**). IL-2, IL-7, IL-15, and IL-32-β modulate the largest number of programs (>½ of all CIPs), whereas M-CSF, IL-21, and CD40L each appear in only two programs. The broad influence of IL-2, IL-7, and IL-15 is consistent with their well-characterized roles in lymphoid activation, proliferation, and survival^47^. Consistent with the highly similar responses described earlier (**Fig. 5c**), IL-2, IL-7, and IL-15 co-regulate 13 programs. In contrast, IL-32-β acts predominantly independently, suggesting more unique regulatory activities.

Next, we inspected the cytokines that modulate a specific CIP (**Fig. 5f**). For example, the myeloid-associated CIP ‘Immune response to viral infection and inflammation’ (ViralResponse) is active in CD14 and CD16 Mono as well as cDCs. In cDCs, ViralResponse is primarily upregulated by the interferons IFN-β and IFN-ω, whereas in monocytes, additional cytokines like IFN-α, IL-1, and IL-12 also induce the program, consistent with their role in promoting antiviral and inflammatory gene expression^48–50^. Conversely, IL-32-β and IL-10 inhibit ViralResponse in monocytes, reflecting interferon counter-regulation. Among lymphoid programs, the CIP ‘immunomodulatory and inhibitory reprogramming of NK cells’ (NK-Immuno) is specific to NK CD56hi, where it is upregulated by IL-10 and downregulated by IL-15 and IL-2, reflecting enhanced inhibitory receptor signaling and dampened activation. In contrast, ‘Cytotoxic effector cell activation and target recognition’ (Cytotoxic-1), is upregulated by IL-15 and IL-2 in NK cells and inhibited by IL-10. Together, these results confirm that the cytokine-program associations observed in our analysis reflect established patterns of cell type-specific regulation in both myeloid and lymphoid populations.

The full set of CIPs reflects the diversity of cytokine responses in our dataset. The majority of CIPs were found in myeloid populations (57 programs, **Fig. 5i**), with fewer linked to lymphoid cells (12 programs, **Fig. 5g**) and shared across compartments (10 programs, **Fig. 5h**). This distribution likely reflects the biology of PBMCs: monocytes and dendritic cells act as innate first responders, expressing broad cytokine and pattern-recognition receptor repertoires, and serve as central coordinators that link innate and adaptive immunity. As a result, myeloid cells engage diverse programs linked to chemotaxis, phagocytosis, antigen presentation, inflammatory activation, and pattern-recognition signaling. In contrast, lymphoid cells, as adaptive effectors, predominantly deploy cytokine-driven transcriptional programs supporting effector function, proliferation, and immune trafficking, resulting in a smaller number of detectable cytokine-induced signatures in this experimental context. Together, our newly defined CIPs link cytokines to cell type-specific functional states, offering a biologically meaningful framework for interpreting cytokine-induced immune responses.

### HuCIRA decodes cytokine responses in disease

We assembled a unified reference compendium (**Fig. 6a**) of cell type-specific gene sets comprising (1) differentially up- and downregulated genes induced by individual cytokine stimulations as inferred in our DE analysis (**Fig. 1c**) as well as (2) gene sets associated with particular immune functions as defined in our CIP analysis (**Fig. 5**). These gene sets enable researchers to decode cytokine activity in independent datasets, providing a means to identify which cytokines may underlie molecular differences between experimental conditions. To facilitate such analyses, we developed huCIRA (short for human Cytokine Immune Response Analysis), an open-source, easy-to-use Python tool that interfaces gseapy^51^ and supports the use of these gene sets in enrichment analyses and differential cell-cell communication inference (**methods**). The input to huCIRA consists of our provided gene sets as well as a user-supplied transcriptomics dataset along with a specification of the conditions across which differential cytokine or program activity should be compared. To illustrate its potential, we applied huCIRA to published transcriptomic datasets from autoimmune diseases, specifically systemic lupus erythematosus (SLE) and multiple sclerosis (MS), as well as spatial transcriptomics data from a non-small cell lung cancer (NSCLC) study.

**Fig. 6.**
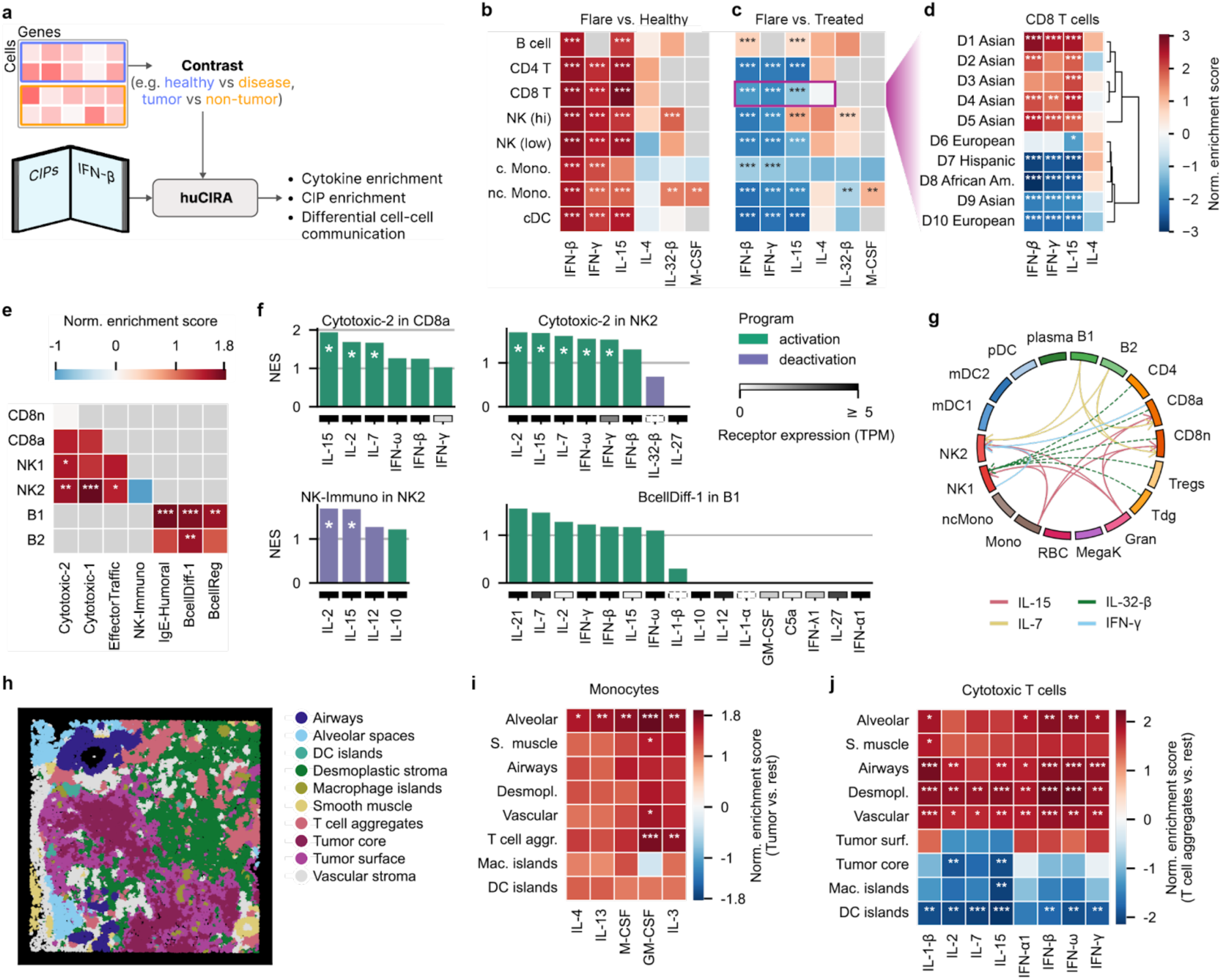
HuCIRA characterizes cytokine and immune program enrichment across conditions, donors, and diseases. **a,** Schematic representation of the cytokine response enrichment analysis. **b-c,** Transcriptomics dataset from PBMCs with 162 SLE and 99 control cases (red: up, blue: down, (*: p<0.1, **: p<0.05, ***: p < 0.01) if significant). **b,** Normalised enrichment scores (NES) for the cytokine activity of flare vs healthy controls. **c-d,** NES for the cytokine activity of flare vs treated across cell types (c) and in CD8 T cells across ethnicities (d). **e-g,** Transcriptomics dataset from PBMCs with 5 MS and 5 control cases (red: up, blue: down, (* if significant). **e,** NES for CIP activity of disease vs control across cell types. **f,** NES for cytokines that regulate the CIPs in d. **g,** Cytokine mediated cell-cell communication (direction indicated by arrow). **h-j,** Spatial transcriptomics dataset from one NSCLC patient (red: up, blue: down, (*) if significant). h, Tissue section with niche annotations. **i,** NES for the cytokine activity of tumor vs other niches for monocytes. **j,** NES for the cytokine activity of T cell aggregates vs other niches for cytotoxic T cells.

### Systemic lupus erythematosus cases exhibit strong IFN signaling activity

SLE is a complex and clinically heterogeneous disease characterized by chronic inflammation affecting multiple organs^52^. There is a well-established association between SLE disease and type I IFNs, with approximately 50% of SLE patients exhibiting elevated type I IFN blood levels, accompanied by increased expression of interferon stimulated genes (ISGs) in their peripheral blood cells^53,54^. Here, we used huCIRA to infer cytokine responses in a published single-cell transcriptomics dataset containing 1.2Mio PBMCs from 162 SLE cases and 99 healthy controls, with associated clinical assessments (healthy, flare, treated) and self-reported ethnicity^55^.

We found that responses to type I IFNs such as IFN-β are strongly enriched in all cell types in flare compared to healthy controls (**Fig. 6b, Fig. S29a**), which is in line with the elevated levels of ISGs reported in the literature^55^. Beyond type I IFNs, we observed prominent IL-15 signaling activity in CD4 and CD8 T cells as well as NK cells, reflecting enhanced survival, homeostatic expansion, cytotoxic function and memory activation^56–58^. Collectively, these effects may amplify tissue-damaging immune responses, consistent with the immune dysregulation characteristic of SLE. Interestingly, we found selective upregulation of IL-32-β signaling in NK cells and non-classical monocytes in flare cases. While the biology of IL-32 in SLE remains largely unexplored^59,60^, it highlights a potential role for IL-32-β in both understanding lupus pathogenesis and monitoring disease activity.

Next, we examined cytokine response changes when comparing flare vs treated cases. Surprisingly, the observed enrichment of IFN-β, IFN-γ, and IL-15 during flares was not reduced in the treatment condition. In fact, it was further elevated (**Fig. 6c, Fig. S29b**). Paired donor-specific analyses (n=10) revealed that this effect occurs specifically in the non-Asian donor ancestry whereas we find the expected enrichment pattern in Asian donors (**Fig. 6d, Fig. S29c-d**). While the limited availability of detailed treatment and genetic information prevents definite conclusions about ethnic contributions, these donor-specific differences highlight the need for caution when interpreting treatment-response biomarker data and underscore the value of personalized medicine approaches in complex diseases.

### Multiple sclerosis cases are marked by cytotoxic immune responses

MS is an autoimmune disease of the central nervous system (CNS) characterized by autoreactive T and B cells that contribute to demyelination and neurodegeneration^61^. Here, we used huCIRA to analyse CIPs and cytokine communication networks in a single-cell transcriptomics dataset containing blood samples from MS patients and control cases^62^. Comparing disease vs healthy samples, we observe an enrichment of CIPs including Cytotoxic-1, Cytotoxic-2, and EffectorTraffic in NK cells in disease samples (**Fig. 6e, Fig. S29e**), reflecting both enhanced killing potential and tissue-trafficking capacity, consistent with previous reports^63^. In B cells from MS cases, enriched CIPs included IgE-Humoral, BcellDiff-1, and BcellReg, indicative of dysregulated differentiation, enhanced humoral activation, and impaired immune regulation, in line with with B cell states capable of producing pathogenic antibodies^64^. Together, these programs point towards a state of systemic immune dysbalance that contributes to the CNS-directed autoimmunity.

Since the activity of each CIP is modulated by one or several cytokines, we next examined the enrichment of these cytokines in the disease context (**Fig. 6f, Fig. S29f**). Cytotoxic-2 is regulated by IFN-γ, IFN-β, IFN-ω, IL-7, IL-2, and IL-15. This finding aligns well with the known biology of NK cell activation and cytotoxic effectorness^65^. BcellDiff-1 is modulated by multiple cytokines of which IL-21, IL-7, and IL-2 are most prominently enriched. These cytokines are key regulators of B cell activation and differentiation, with IL-21 particularly driving proliferation and plasma cell maturation^66–68^. Finally, we examined the cytokines relevant to the inhibitory NK-Immuno CIP, which was downregulated in the disease state (**Fig. 6e**). IL-10 promotes its activity, whereas IL-12, IL-15, and IL-2 suppress it (**Fig. 6f**). This pattern is consistent with the enhanced enrichment of the cytotoxic CIPs observed earlier.

Next, we investigated differential cell-cell communication networks in the MS dataset (**Fig. 6g**). A cytokine communication (sender-receiver interaction) was considered valid when the cytokine was found to be differentially expressed in the sender cell and the receiver both expressed the receptor and showed a significant huCIRA enrichment (methods). In line with the identified biology from our CIPs and cytokine activity enrichment analysis, we found enhanced IFN-γ (from NK cells and CD8a T cells), IL-15 (from monocytes and granulocytes), and IL-7 (from B cells) signaling in NK cells in disease vs healthy samples.

### Tumor-localised GM-CSF activity shapes monocyte reprogramming

We next applied huCIRA to investigate cytokine activities in a non-small cell lung cancer (NSCLC) spatial transcriptomics dataset (**Fig. 6h**). We compared signaling activity between immune cells in tumor and non-tumor regions. Notably, monocytes in the tumor exhibited significantly enriched GM-CSF and IL-3 signaling, particularly relative to T cell aggregates and the alveolar niche (**Fig. 6i**). GM-CSF is often overexpressed by tumor cells in NSCLC and other solid tumors^69–71^, and its presence promotes the recruitment and differentiation of monocytes into immunosuppressive tumor-associated macrophages (TAMs)72. These findings suggest that GM-CSF-driven monocyte activation is a spatially regulated process within the TME, shaping monocyte reprogramming to support immune suppression and highlighting the spatial as well as molecular complexity of the tumor-supportive myeloid compartment described in the molecular atlas^73^.

Beyond the tumor core, tumor microenvironments (TMEs) contain distinct immunological niches such as tertiary lymphoid structure-like regions^74^ that shape local immune responses. Given the critical role of T cells in tumor cell killing, we compared tissue regions identified as ‘T cell aggregates’ with other niches (**Fig. 6j**). Responses to cytokines promoting T cell function (IL-1-β, IL-2, IL-7, IL-15, IFN-α1, IFN-β, IFN-ω, IFN-γ) were downregulated in T cell aggregates relative to the tumor core, macrophage and DC islands but upregulated compared to other niches. These data suggest a spatial separation between lymphoid-like cytokine environments and myeloid-instructive niches within the NSCLC TME, reflecting a balance between immune surveillance and immune evasion. From a therapeutic perspective, these findings support the rationale for spatially targeted immunomodulation, for example, enhancing IL-15/IL-2 or Type I IFN signaling in T cell aggregates to potentiate local effector responses, or blocking GM-CSF/IL-3 signaling in tumor cores to prevent monocyte-to-TAM polarization and restore immunocompetence.

## Discussion

Here, we present the Human Cytokine Dictionary, the largest single-cell perturbation dataset of primary human immune cells to date, which facilitates the systematic analysis of differential gene expression responses across 12 major cell types and 90 cytokines. The Human Cytokine Dictionary enables the systematic characterization of donor-specific cytokine-driven immune activities, uncovering intriguing variability among donors with a pre-existing interferon-signaling state but also describing meaningful consensus responses. The comparison to an earlier mouse dataset revealed both similarities but also substantial differences in cytokine stimulation responses. The large number of cytokines and cell types allowed inference of a detailed network of cellular communication via cytokines, finding, for example, that IL-32-β, which notably does not have a mouse homologue, converts a type I antiviral response toward a strong neutrophil-driven inflammatory program.

To complement gene sets from our DEG analysis, we used DRVI to identify cytokine-induced immune programs (CIPs) that link cytokines to cell type specific functional states, and thus offering a biologically meaningful framework for interpreting cytokine induced responses. Finally, we developed the huCIRA python package, an easy-to-use, open-access resource which consolidates our results and enables the community to study cytokine response and CIP enrichment, as well as differential cell-to-cell communication, in their own datasets. We showed the utility of huCIRA for studying cytokine signaling in autoimmune disease and cancer, revealing context-specific cytokine profiles in single-cell and spatial transcriptomic datasets.

As the first large-scale single cell cytokine perturbation dataset in human tissue, the Human Cytokine Dictionary sets the stage for broader perturbation atlasing efforts. While extensive, our study was anchored at a 24-hour time point. Capturing additional earlier and later responses would provide valuable complementary information on signaling dynamics. Expanding cytokine perturbation screens across multiple organ systems^75^ would establish the foundation of a comprehensive human perturbation atlas. Moreover, although we found intriguing donor response variability, our sample size currently provides limited resolution in clearly linking this variability to genetic or demographic traits. A follow-up screen with more donors could characterize such differences.

In line with observed scaling laws, the size of our dataset enables AI models of cytokine perturbations^76,77^ paving the way towards virtual cell models of cytokine biology^78^. Future expansions of perturbation atlases across tissues and conditions will therefore not only deepen our biological understanding but also accelerate AI-driven inference, guiding experimental design and prioritizing clinically relevant cytokine perturbations. In summary, the Human Cytokine Dictionary provides an essential resource for mapping cytokine-driven immune responses in humans. It bridges basic immunology, clinical translation, and artificial intelligence by offering both a foundation for mechanistic discovery and a training ground for predictive modeling.

## Methods

### PBMC culturing and cytokine stimulation

#### Source and thawing protocol

Viably frozen healthy human PBMC samples (6 females, 6 males) were purchased from Bloodworks Northwest (**Supplementary Table 1**), who confirms that donors provide informed consent and that sample collection was approved by their institutional review board or equivalent ethics committee. PBMC samples that were purchased for this work were received de-identified, and no additional ethical approval was identified to be required for this work. Prior to cytokine treatment PBMCs were thawed in such a way as to maximize integrity and viability and to minimize shock that could potentially alter gene expression and/or cell viability. PBMC vials were placed in a 37C water bath for 1 minute to thaw. For the dropwise media additions during PBMC thawing, RPMI + 10% FBS media warmed in a 37C water bath was used. The entire contents of the PBMC vial was transferred into a 50mL conical and a series of dropwise additions of media were pipetted into the tube. A cell aliquot was set aside for viability checks and counting while the majority of PBMCs were centrifuged at 300g for 10 minutes and the media was removed at the conclusion of centrifugation.

#### Cell resuspension and cell culture media

PBMCs were cultured in 96-well plates (Nunc™ Edge™ 96-Well, Non-Treated, Flat-Bottom Microplate, Thermo Fisher cat. #267578). For the cytokine stimulations, RPMI media (ATCC SKU: 30-2001) supplemented with 10% FBS (Gibco cat. #A4766801), 2-Mercaptoethanol (50uM final, Thermo Fisher cat. #31350010), MEM Non-Essential Amino Acids Solution (1X, Thermo Fisher cat. #11140050) and Sodium Pyruvate (1mM final, Thermo Fisher cat. #11360070) was used.

#### PBMC washes and harvest

To wash and harvest cells post cytokine stimulation, we resuspended PBMCs in their incubation media pipetting 5x with a P-200 multi-channel pipette set to 180 µL. After resuspension PBMCs were transferred to a Protein LoBind® Plate 96-well 1mL plate (cat. #951033308) and placed on ice. Stimulation 96-well plates were washed with 100ul of D-PBS to lift off any residual PBMCs and transferred into the Protein LoBind® Plate 96-well plate. After collection, cells were centrifuged at 200xg for 10 minutes at 4C.

#### Cytokines and cytokine stimulation

Cytokines were purchased from Biotechne, PBL assay science and Acro Biosystems as indicated in **Supplementary Table 2**. Cytokines were reconstituted at the recommended stock concentrations using the recommended reconstitution buffer. Cytokines were diluted to working concentrations on the day of stimulation. For each donor, 1 million PBMCs cells were plated in each well of a 96-well plate at a final concentration of 5,000/ul in cell culture media. Cells were stimulated with cytokine or 1x PBS (control) for 24 hours.

#### PBMC harvesting after cytokine stimulation

After 24 hours of cytokine stimulation PBMCs were transferred to Protein LoBind® 96-well plates (cat #951033308). In order to recover any residual cells, wells were washed once with 100ul of 1x PBS and the entire volume was transferred to the Protein LoBind® 96-well plate. PBMCs were centrifuged at 200xg for 10 minutes at 4C and supernatant was removed prior to starting the High-Throughput Evercode Cell Fixation v3 with Integra Assist Plus workflow.

### scRNAseq data generation

#### Fixation

Plates of cytokine-treated PBMC samples were fixed using the High-Throughput Evercode Cell Fixation v3 with Integra Assist Plus workflow. Fixed cells were resuspended in 75uL of the storage buffer. Three 96-well plate aliquots were made, two of which had 35ul of fixed cells each and 1 plate containing the remaining 5-10ul of fixed cells that was used for getting cell counts prior to barcoding. All plates were stored at -80C until the day of barcoding. The counting plate was thawed and counted to determine cell concentrations for each sample prior to barcoding. To ensure consistent cell proportions during cell barcoding, fixed cells were transferred to plates of variable volumes of dilution buffer.

#### Barcoding

Each plate of fixed cells underwent three rounds of combinatorial barcoding using Section 1 of the Parse Biosciences GigaLab protocol. After barcoding, cells were distributed into individual sub-libraries of ∼100,000 cells and lysed according to Parse Biosciences GigaLab protocol.

#### Sublibrary Preparation

Lysates were processed through Sections 2 and 3 of the Parse Biosciences GigaLab protocol in batches of 16 sublibraries each. At the end of Section 3, an Illumina i7 and i5 index was added to each sublibrary to produce a sequenceable molecule. Each sublibrary was then converted into an Ultima Genomics-compatible sequencing library and sequenced on the UG-100 sequencer at approximately 31,000 mean reads per cell.

#### Split-pipe processing

Fastq files generated from Ultima Trimmer workflow were processed through a modified version of spit-pipe v1.4.0 to accommodate libraries processed through the GigaLab. For each sublibrary, split-pipe generated a cell by gene matrix (count_matrix.mtx) as well as relevant metadata for each cell (cell_metadata.csv) and a list of genes to which split-pipe maps fastq files (all_genes.csv). Downstream analysis was performed Gene and transcript definition was based on Ensembl release 109 and genome build 38 (https://ftp.ensembl.org/pub/release-109/gtf/homo_sapiens/Homo_sapiens.GRCh38.109.gtf.gz)

#### Data processing and clustering

Data processing and clustering was performed in Python 3.10 with Scanpy (v1.10.3). Cells were filtered for a minimum of 400 genes and maximum of 7000 genes detected and <15% mitochondrial content to filter out low quality cells and some doublets. We additionally filtered doublets using Scrublet^79^ with default parameters and clustered cells as described below. Cells from clusters that contained >50% predicted doublets were removed to produce a final AnnData object containing 9,697,974 cells that were then re-clustered for cell type annotation.

Transcript counts per cell were normalized to 10,000 per cell and log transformed (sc.pp.normalize_total, sc.pp.log1p). Highly variable genes were identified with sc.pp.highly_variable(min_mean**=**0.0125, max_mean**=**3, min_disp**=**0.25). The resulting matrix subset on highly variable genes was scaled to unit variance and zero mean (sc.pp.scale, max_value**=**10). Principal component analysis (PCA) was performed on the scaled highly variable genes (sc.pp.pca). The first 30 PCs were retained for downstream steps. A k-nearest neighbor graph was built in the 30 PC space (sc.pp.neighbors, n_neighbors = 15, n_pcs = 30). Next, we ran UMAP to embed the neighbor graph in two dimensions (sc.tl.umap). Finally, we performed leiden clustering to group similar cells together (sc.tl.leiden, resolution = 1.0).

#### Cell type annotation

We performed differential gene expression analysis to identify cluster-specific gene markers. Clusters lacking unique markers were merged with the nearest cluster. We then manually annotated each cell type using standard PBMC marker genes^10,11^. Marker expression was visualized with dot plots to confirm specificity (see **Fig. S1b**).

### Differential cell type abundance analysis

We analyzed both total cell type counts and cell type composition across donors and cytokine treatments. For total counts, we first adjusted for potential technical effects caused by well position by fitting a linear model between the total number of cells recovered (summed across cell types) and the distance of each well to the plate’s center row. We used this model to normalize total counts across rows. The log2FC is the mean of log2FC values of the count for a given cell type and perturbation to the PBS count across donors. To calculate p-values, we used a two-sided Wilcoxon signed-rank test comparing, within each donor, the normalized counts under perturbation versus PBS. For compositional analysis, we used scCODA^12^ (design ∼cytokine+donor) with CD4 T cells as the reference category as they showed relatively stable total cell numbers across cytokine perturbations.

### Differential gene expression analysis

We utilized pertpy’s^80^ interface to edgeR^14^ to identify differentially expressed genes (DEGs). DEG analysis was performed on pseudobulked data, where all cells belonging to the same cell type and cytokine condition were aggregated per donor. For each cell type-cytokine subset, we included only donors for which both the PBS and cytokine-treated pseudobulks contained at least 10 cells. Within each subset, we further retained only genes that were expressed in at least 5% of cells in either the PBS or cytokine-treated condition. Subsequently, DEGs were computed separately for each cell type-cytokine subset using the design formula ∼donor + cytokine, with differential expression assessed relative to the PBS control condition.

To mitigate potential technical artifacts, we conducted a secondary differential expression analysis. This analysis employed the same overall design but, instead of pooling cells from all six PBS wells, we performed six independent statistical tests per cytokine - each using a distinct PBS well as the reference. Our rationale was that biologically driven differential responses would be consistently observed across all six comparisons, whereas technical artifacts associated with well position on the 96-well plate would not.

In this secondary analysis, a gene was considered differentially expressed if it satisfied both adjusted p-value < 0.1 and |log_2_FC| > 0.25. For each gene, we then counted the number of PBS wells for which it was identified as differentially expressed. This information is supplied alongside the initial results of the DEG analysis (computed on pooled PBS cells) and can be used as a filter criterion, e.g. for the human cytokine dictionary we define a gene to be differentially expressed if its adjusted p-value < 0.05, the corresponding |log_2_FC| > 0.25 and a gene is differentially expressed with respect to at least 4 out of 6 wells.

To remove additional unstable or well-biased genes, we used the per-PBS well log2FC values to identify genes with an excessive standard deviation to mean ratio across wells. We calculated the mean-to-stddev-ratio = stddev(gene)/(|mean(gene)|+0.25) across the six reference wells for each gene in each cytokine-cell type condition. Genes with a mean-to-stddev-ratio above 1 were defined as having a high mean-to-stddev-ratio in a given condition. Genes that had a high mean-to-std-ratio for at least 10 cytokines in a given cell type were defined as having a high mean-to-std-ratio in that cell type. Genes with a high mean-to-std-ratio in at least 5 cell types were removed from the DEGs.

### Calculation of the response magnitude, strong impact classification, and tissue-specificity index

We used two different measures to calculate the global response magnitude M of the cellular transcriptome to a cytokine stimulation: First, we used the (UMI normalized) cpm (with a pseudocount of 1) pseudobulk-averaged gene expression vector v. The values are subset to genes with a total average pseudobulk count of at least 20 in at least one of the stimulation conditions for that cell type and log2-transformed before calculating the Euclidean distance between the PBS control and the cytokine:

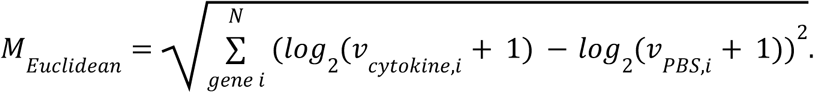

Second, we use edgeR-calculated significance (padj) and fold change (FC) values. Significance values are clipped at 10^-10^, and a distance is calculated as

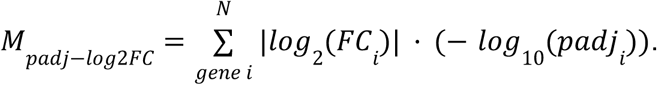

Both metrics are winsorized at the 95% percentile, then normalized to values between 0 and 1. The used overall magnitude M is the average of these two values. To determine whether a cytokine has a strong impact on a given cell type, we set a threshold value *T* at three times the mean of the response magnitude of values below the 35th percentile. This empirically captures the part of the distribution without substantial effects for each cell type without relying on a particular shape of the distribution (cf. Fig. S6f).

The tissue specificity index (tsi) of the response to a cytokine c is calculated as^15^:.

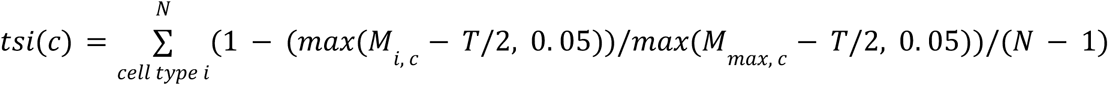

wherein the deduction of half the threshold value *T* helps to reduce noise.

### Characterization of donor-specific responses

For each donor-cell type-cytokine condition, we filtered all analyses to genes with at least 10 raw reads in the pseudobulk. To characterize the donor baseline state, we calculate a baseline FC as the ratio of per donor gene cpm (pseudocount of 1) in the PBS condition to the median gene PBS cpm across donors, subsetting to protein-coding, annotated, non-ribosomal, non-mitochondrial, and non sex-specific (located on the X or Y chromosome) genes using pybiomart for increased interpretability.

Per-donor response magnitudes were calculated by the Euclidean measure described above for all conditions with at least 1000 filter-passing genes. The per-donor response magnitude was divided by the square root of the length of the donor gene expression vector *v* for normalization. Donor-specific FC values for each cytokine and cell type were calculated by dividing the gene cpm for a condition by its PBS value, discarding any condition with fewer than 20 cells.

We also calculated a non-normalized raw response strength to quantify trends in absolute response strength differences between donors:

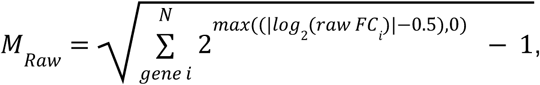

wherein the subtraction of 0.5 helps to deal with higher noise levels for individual donor pseudocount comparisons.

### Donor outlier and substantial substructure detection

Outlier detection serves to find donors whose responses differ substantially from the consensus response across donors, whether due to a much weaker response or a directionally different response. To find such outlier donors in cell type-cytokine conditions, we first calculate the Pearson correlation *r* between the edgeR-calculated donor-consensus log2FCs across donors and the donor-specific log2FCs for each donor. We then get the median of the top 8 correlation values. Any donor whose correlation *r* fulfills

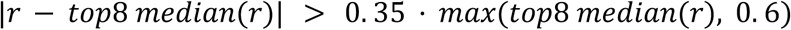

is considered an outlier for that particular cell type and cytokine.

The goal of substantial substructure detection is to find subsets of donors that have substantially higher correlations within the subsets than between subsets, indicating substantial but genuinely different types of responses to that cytokine depending on the donor state. The following algorithm was repeated for Leiden resolution parameters of 0.3, 0.5, and 0.7, taking the smallest parameter that lead to successful substructure detection:

First, donors (for a given cell type and cytokine) are clustered on their pairwise donor-specific log2FC correlations using the Leiden algorithm with a given resolution parameter. We required at least one community of size 4 and a second community of size 2 to consider this step a success. We then calculated the mean of correlation values within the donors of these groups (discarding the diagonal) and the between the donors of different groups. If two such communities have a margin of at least max(0.2, 0.3 * max(r_donor_i,j|i≠j)) to another community, substantial substructure is present.

### Receptor expression analysis

Receptor genes for each cytokine were manually curated for all cytokines except IL-32-β, which does not have a known receptor gene. Receptor expression was calculated for the PBS control samples. When a receptor consisted of multiple subunits, we used the minimum cpm expression across these genes on the assumption that all subunit genes must be expressed for the receptor to be active. When there were multiple possible receptors for a single cytokine, we used the maximum expression across all alternative receptors on the assumption that any of these receptors is sufficient for activity. We do not take into account possible differences between different receptors for a single cytokine. The relative receptor expression was calculated by dividing by the raw expression by the maximum expression value across all cell types.

In considering the relationship between receptor expression and the response magnitude, we first note that we observe an empirical threshold of around 8 cpm receptor expression necessary to observe responses. For classification of responses, we chose thresholds with some distance to this empirical threshold to avoid classifications that depend too sensitively on its exact value. Accordingly, a response magnitude 1.5 times larger than the strong impact threshold along with a receptor expression below 5 cpm is therefore considered a potential candidate for a secondary response. Conversely, we considered all cytokines with a response magnitude below the strong impact threshold and a receptor expression above 32 to have weak responses despite very clear strong receptor expression. A cell type with a receptor expression above 16 and a response magnitude 1.5 times larger than the strong impact threshold is considered a primary target for a cytokine.

### Comparison to the mouse cytokine dictionary dataset

Single cell data for the mouse cytokine dictionary was downloaded from https://www.immune-dictionary.org/app/home and converted to scanpy compatible format via zellconverter^81^. The downloaded dataset contains 110,378 cells with 31,053 profiled genes. We used pybiomart (https://jrderuiter.github.io/pybiomart; version 0.2.0) to convert genes to human homologs; this reduced the number of genes to 16,004. The number of counts per cell was normalized (scanpy.tl.normalize_total) and subsequently averaged per cell type-cytokine combination to obtain pseudobulks per mouse replicate. Due to the *in vivo* setting of the mouse dataset, there is only a single cytokine stimulation condition available per individual mouse. For the replicate-specific comparison of log2 fold changes, we hence used the weighted mean across all PBS-treated mice, where weights are proportional to the number of cells per mouse.

Our human dataset was preprocessed analogously except that donor-specific log2 fold changes were computed with respect to the donor-specific PBS-treated cells. The response magnitudes for the mouse dataset were downloaded from the figure source data (Source data Fig. 1). For the donor/replicate-unspecific comparison of log2 fold changes, we used the weighted mean across all samples within a cell type-cytokine condition where weights are proportional to the number of cells per sample.

For the comparison with the independent human PBMC dataset^33^, we obtained raw count data and cell type annotations from the Gene Expression Omnibus (GSE96583) and normalized counts on a per-cell basis (scanpy.tl.normalize_total). Because donor information is not available in the metadata, we computed average expression profiles per each cell type and stimulation condition (control, IFN-β) by weighting all cells equally.

#### Reference database construction

To construct the reference dataset of cytokine-cell relationships, we used the ImmunoGlobe^34^ and ImmuneXpresso^35^ databases. As both databases use Cell Ontology IDs (CLIDs) to identify cell types, they were merged on these keys and cytokine identities to make a dataset of unique cell-cytokine relationships. These included both cell source (that is, cell secreting a cytokine) relationship and cell target (cytokines having some kind of an effect on a cell) relationships. The unique CLIDs present in the resulting merged dataset were examined, and manually assigned to groups corresponding to our cell cluster identities when possible. More generic CLIDs, such as ‘T cell’, which could not be assigned to a particular cluster, were excluded for analysis. The overall dataset construction process can be seen outlined in **Fig. S12a.**

As neither database differentiated between NK cell subtypes, NK CD56low cells were chosen to represent NK cells given they are the most common in peripheral blood^40^. Similarly, as most monocyte annotations did not differentiate between CD14 or CD16 monocytes, CD14 monocytes were used to represent monocytes more generally^82^. Finally, for the mouse cytokine dictionary comparisons to the reference dataset, cDC1 cells were used as the representative cDC cell type.

For the cytokine stimulation comparisons, supporting paper counts were winsorized at the 95th percentile and scaled between 0 and 100 per cell type (as both the response magnitude and mouse cytokine dictionary response magnitude values were scaled in a similar way). For the high quality analysis, only annotations shared between the two databases for cell types with greater than 25% coverage (that is, with more than 20.25 cytokines with supporting paper information) were used. Cell types with less than this cutoff were cDCs and HSPCs. Mouse cytokine dictionary values for response magnitude were downloaded from figure source data.

For the cytokine secretion comparisons, supporting paper counts were scaled between 0 and 1 per cytokine (similar to how the relative mean production and mouse cytokine relative expression were scaled between 0 and 1 per cytokine). Mouse cytokine dictionary source figure data was used for the comparisons. Only high-quality annotations of secretion which had both mouse and human measurements were used to calculate Spearman’s ρ. Cytokines with higher numbers of annotations were considered to be any with six or more annotated secreting cell types.

### Grouping cytokine responses

Genes with a maximum expression of 20 cpm, an average expression of at least 4 cpm, and a total maximum pseudobulk count of at least 20 across all stimulation conditions were considered robustly expressed in a given cell type. We calculated the similarity between different cytokine stimulation conditions in a cell type by the Pearson correlation of consensus log2FC values for all expressed genes. To find highly similar cytokines across cell types, we averaged the similarity (Pearson correlation) values across all cell types for which both cytokines have a strong impact (cf. *Calculation of the response magnitude and impact classification*). If there are at least 3 cell types with strong responses and the mean r is larger than 0.7, two cytokines were considered highly similar. To sort cytokines into highly similar groups, we applied this relationship transitively.

To group cytokines more loosely within a cell type, we applied the Leiden algorithm^83^ using the python leidenalg package using the Pearson correlation similarity matrix as input. For each cell type, we first subset to cytokines with strong impact, and then constructed a weighted, undirected graph where each node represents a cytokine and each edge weight corresponds to the Pearson correlation of consensus log2FC values between two cytokines. Self-loops were eliminated by setting the diagonal of the similarity matrix to zero. We then applied the Leiden algorithm using the CPMVertexPartition method with a resolution parameter of 0.5 to generate communities. To further cluster cytokines across different cell types, we repeated the Leiden clustering procedure using grouping counts, i.e. the number of times two cytokines were clustered by the Leiden algorithm within a cell type. Cytokines were filtered to those that participated in clustering for at least two cell types. Row sums were computed and used to symmetrically normalize the matrix by multiplying with the inverse square roots of the row sums (with a small epsilon added to avoid division by zero). Pairwise cosine distances were calculated for the normalized matrix. Leiden clustering (resolution parameter = 0.5) was applied to the distance matrix to identify consensus cytokine communities.

### Cytokine production, crosstalk, and identification of potential secondary responses

Human immune-related cytokines (broadly defined), cytokine genes, and families were manually curated and classified into families using the mouse cytokine dictionary^8^, immuneXpresso^35^, Janeway’s Immunobiology^36^, and other information sources. A cytokine gene was considered expressed according to the same criteria described above for genes in general (mean cpm > 4, max cpm > 4, at least 20 total counts in at least one condition). For cytokines consisting of multiple subunits, we took the minimum values across the subunit genes on the assumption that all subunits must be expressed for the cytokine to be active. We only display cytokines that are expressed in at least one cell type. To show the production of cytokines by cell type, we show the mean for a cell type divided by the maximum mean across cell types.

To determine whether cell-cell communication is present, we test whether a cytokine is expressed in a cell type and whether it has a strong impact on a target cell type. Strong impact was calculated as described above. If both are the case, we consider the cytokine as mediating a response between the two cell types. To quantify the connection strength for a cell type-cytokine-cell type connection, we define sender strength as the relative expression of a cytokine in a cell type (compared to all other cell types), considering only cell types in which the cytokine is expressed. We additionally required a mean cpm > 10 in at least one cell type. Receiver sensitivity is defined as the response magnitude for stimulation with the cytokine for a given cell type. The product of sender strength and receiver sensitivity is the interaction score. The overall connection strength between two cell types is the sum of their interaction scores across cytokines. For the overall cell-cell connection plot, we subset to connections with an interaction score of at least 0.25 to prune less relevant connections for plotting purposes.

To quantify cytokine-cytokine signaling cascades, we classified cytokines according to the consensus log2FC as significantly upregulated (padj<0.05, log2FC>1) or downregulated (padj<0.05, log2FC<-1) in response to a cytokine stimulation in a given cell type. For cytokines with multiple subunits, we required that all genes fulfill both the significance and the fold change requirements. We used the same thresholds to count up- and downregulation by cytokine family. Upregulation of the same cytokine across different cell types is counted multiple times. To account for differing family sizes, we divide by the maximum number of up- or downregulated members for each family.

To identify secondary responses, we first identify primary and secondary target cell type (cf. *Receptor expression analysis*). We then identify all cytokines upregulated in primary target cells with a log2FC>1 as potential secondary cytokines. For each potential secondary cytokine and secondary cell type, we check whether the secondary target cell type is a primary target cell type for direct stimulation with the secondary cytokine and whether the response similarity, i.e., the Pearson correlation of log2FC values, is larger than 0.5. Any potential secondary cytokine that fulfills the criteria is considered a likely genuine secondary cytokine affecting a given cell type.

### Inferring cytokine-induced immune programs (CIPs)

To discover universal cellular programs within and across cell types, we trained DRVI^46^ for 50 epochs using all genes, employing a 256-dimensional latent space. Donor ID was included as a batch covariate. The encoder and decoder subnetworks each comprised three 512-dimensional hidden layers. To encourage positive latent factors, we incorporated a Continuously Differentiable Exponential Linear Unit (CELU) activation function^84^ with α=0.1 in the bottleneck layer. This training yielded 147 non-vanishing factors {z_i}_i=1…147, i.e., factors having absolute value more than one in at least one cell. For each factor z_i, DRVI provides a sorted list of genes along with an interpretability score that indicates how relevant each gene is to that factor. We kept the 133 factors that had at least one gene with an interpretability score ≥ 0.5. Furthermore, for each of the 133 programs identified in this manner, and for every combination of donor, cytokine, and cell type, we performed separate t-tests against their six associated PBS control cell distributions. A contrast was considered significant if all of the following criteria hold:

1. The t-test yielded a multiple-testing corrected p-value < 0.001 in at least four out of six tests (i.e. against at least four out of six PBS wells). Multiple testing correction (Bonferroni) was applied within a cell type - cytokine condition across all programs.
2. (2) The absolute program activity 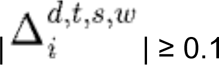, where the program activity is defined as

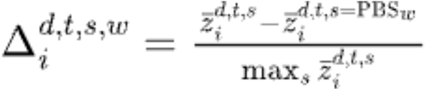

with

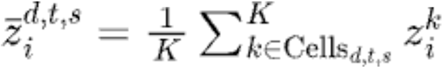

where K is the number of cells within a donor (d), cell type (t) and cytokine stimulation (s) combination, and where w is indicating the contrasting PBS well (out of the six PBS wells).

1. To ensure that identified programs reflect consistent responses rather than donor-specific outliers, we required each contrast to meet the aforementioned significance criteria (1) and (2) in at least four donors.
2. The cytokine induced program regulation direction was sufficiently robust across donors. Robustness was assessed by a simple voting scheme: for each cell type and cytokine, we counted how many donors showed upregulation versus downregulation of a program, and considered the contrast significant only when the absolute difference between these counts was at least four.

Taken together, these filtering steps resulted in 95 significantly differential programs. Out of these, 13 programs were furthermore manually excluded as they either represented cell type-specific effects (9 programs), technical artifacts (3 programs), or lacked biological relevance (1 program). The remaining 82 programs were then manually annotated and categorized into 11 general classifications. We refer to these programs as cytokine-induced immune programs (CIPs), which enable the inference of CIP activities within single-cell transcriptomics datasets.

### Using the human cytokine dictionary to detect cytokine responses, functional immune states, and differential cell-cell communication patterns

Our newly constructed human cytokine dictionary provides cell type-specific gene signatures that are up- or downregulated in response to cytokine stimulation, as well as signatures that define cytokine-induced immune programs (CIPs). These gene sets can be incorporated into gene set enrichment analyses (GSEA) to assess differential biological patterns across conditions. Our GSEA workflow uses the prerank module from gseapy^51^ (the dictionary is also compatible with alternative enrichment tools) and takes as input a gene set (e.g., genes upregulated by IL-10 or those associated with a given CIP) together with two count matrices, X_a and X_b, representing the transcriptomic profiles of cells from the two conditions to be compared. In the first step, we compute average expression vectors for each condition and combine these into a single vector of expression differences where each component is normalized by the sum of the within group standard deviations, i.e., 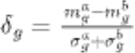. The components of this vector are then ranked, placing the largest differences at the top and the smallest at the bottom. The enrichment test proceeds by iterating through the ranked vector gene by gene, increasing a running sum if the current gene belongs to the user supplied gene set and decreasing the running sum if it does not. The rationale of this procedure is that gene sets whose members cluster near the top or bottom of the ranked difference vector - corresponding to large positive or negative expression differences across conditions - produce a pronounced peak in the running sum. This peak, taken as the maximum absolute deviation, defines the observed enrichment score. Statistical significance is estimated via a permutation test by generating a reference null distribution through repeated iterations with randomly sampled gene sets; we set the number of random permutations to 20,000. The final enrichment score is normalized by the mean of this null distribution.

The enrichment procedure involves several hyperparameter choices. To ensure robustness to these we performed the analysis across a range of parameter values and integrated the resulting outcomes to determine statistical significance. For cytokine-based gene sets, we used results from our differential expression analysis but applied five minimum thresholds (0.8, 0.9, 1.0, 1.1, 1.2) for log2 fold changes. In parallel, we also applied five expression cutoffs (6–10 cpm) to exclude lowly expressed genes. This yielded a 5 x 5 grid of enrichment results for cytokine gene sets and five results for CIPs, for which log2 fold-change thresholds are not applicable.

We only considered an enrichment to be significant if all of the following criteria were met:

1. the test could be performed for at least one-third of the hyperparameter combinations, ensuring that results were not driven by a single parameter choice; tests were skipped when the resulting gene set contained fewer than eight or more than 1000 genes;
2. at least two-thirds of the valid tests produced *P* < 0.1; and
3. the direction of enrichment was consistent across hyperparameter settings, thereby excluding cases in which significance arose for different hyperparameter combinations but with opposing signs.

The output of the cytokine enrichment analysis can be combined with gene expression information to infer differential cell-cell communication. To infer potential senders of cytokines we perform a Wilcoxon test for differential gene expression on cytokine production-associated genes to compare each cell type against all other cell types. We consider a cell type to be a sender if the adjusted (Benjamini-Hochberg) p value < 0.1, the log2 fold change is positive and the average gene expression exceeds 10 cpm. A cell type is considered a receiver if we find significant enrichment for a cytokine and if the average receptor gene expression exceeds 10 cpm. For multimeric receptors we apply the expression threshold criterion to the gene with the lowest average gene expression.

We showcase the usage of our cytokine dictionary on three independent datasets:

#### Systemic lupus erythematosus

As a first application, we used the enrichment framework to analyze a single-cell dataset of systemic lupus erythematosus^55^. The dataset comprises 1,263,676 cells with expression profiles for 30,933 genes. Cells were obtained from 261 donors classified by disease status as healthy (99 donors), managed (146), or flare (19); for 10 donors in the flare group, there is a second set of cells available in the post-treatment (treated) stage. For enrichment analysis, we used the default hyperparameters and filters described above. For the donor-specific analysis, we applied a cell type-specific donor filter and only ran the analysis on donors with at least 20 cells in that cell type. The data was normalized (scanpy.pp.normalize_total) and log (scanpy.pp.log1p) transformed prior to the enrichment analysis. The clustering of enrichment results for matched samples was done with scipy.cluster.hierarchy using the method “average” and euclidean distance as metric.

#### Multiple Sclerosis

We secondly demonstrate the use of CIPs as well as differential cell-cell communication on a multiple sclerosis dataset^62^ comprising 65,326 single cells with expression profiles for 10,266 genes. The dataset includes cells from individuals with multiple sclerosis (35,483 cells) and healthy controls (29,843 cells), sampled from either peripheral blood (21,664 cells) or cerebrospinal fluid (CSF; 21,654 cells). Before performing enrichment analysis, we normalized total counts per cell (scanpy.pp.normalize_total) and applied log-transformation (scanpy.pp.log1p). Enrichment was carried out using the default parameters described above.

#### Non-small cell lung cell cancer

Third, we apply our dictionary to a spatial transcriptomics dataset generated using CosMx Spatial Molecular Imaging (Nanostring) from a non-small cell lung cancer tumor sample obtained from a single patient^73^. The dataset contains 340,644 cells, distributed across six spatial sections and includes 960 genes. We focus on section 28 which contains 58,423 cells. Each cell has a cell type annotation as well as a niche label, which reflects its neighborhood properties (e.g. cell type composition), which we use to define contrasts for our enrichment analysis. This dataset is particularly challenging for our enrichment analysis due to the relatively small gene panel (relative to the previous non-spatial single cell datasets) and consequently the reduced gene overlap with our cytokine gene signatures. We introduce an additional filter criterion and only test enrichment of cytokines for which at least 10% of the signatures overlap with the genes available in the dataset. Furthermore we increased the threshold of the first robustness criterion for statistical significance (1) and required at least 50% of the hyperparameter grid to produce valid results (i.e., to not be filtered out due to insufficiently small gene sets). All other hyperparameters were kept at their default values. Prior to the enrichment analysis we normalized the total counts per cell and log transformed the data as described before.

#### Code usage example

Usage of huCIRA requires only a few lines of code:

The input transcriptome should include common immune cell types across different conditions. After loading in your data and the human cytokine dictionary, the run_one_enrichment_test() function computes cytokine enrichment scores of one queried immune cell based on gene expression profiles of the chosen conditions, as demonstrated in the below pseudocode:

~~~
import scanpy as sc
import hucira as hc
# 1. Load your data
adata = sc.read h5ad(“your transcriptome.h5ad”)
human cytokine dictionary = hc.load human cytokine dict()
# 2. Run cytokine enrichment analysis for B cells between healthy
and diseased patients
enrichment results = hc.run one enrichment test
 (adata=adata,
 df hcd all = human cytokine dictionary,
 contrasts combo = (“healthy”, “disease”),
 celltype combo = (“B cell”, “B”),
 contrast column = “condition”,
 celltype_column = “cell_type”,
 direction = “upregulated”,
 threshold expression = 0.0
)
# 3. Investigate enrichment scores of cytokines in your data enrichmentresults
~~~

### Statistical analysis

The statistical tests used are described for each analysis in the corresponding text. Error bars on bar plots show the standard unless otherwise mentioned. Boxplots show median and interquartile range; whiskers are drawn to the farthest datapoint within 1.5 times the interquartile range from the nearest hinge.

## Data availability

Single cell data will be available at https://www.parsebiosciences.com/datasets/10-million-human-pbmcs-in-a-single-exp eriment/. Additional analysis results are made available as Supplementary Tables. The dataset is available under the CC BY-NC 4.0 license.

## Code availability

Code used for the analysis of this study is available on GitHub at https://github.com/theislab/HumanCytokineDict. Our python package huCIRA is available at https://github.com/theislab/huCIRA.

## Acknowledgements

We thank Niki Kilbertus for the discussion of statistical analyses. LO acknowledges funding by the EMBO Postdoctoral Fellowship Program. SB is supported by the Helmholtz Association under the joint research school “Munich School for Data Science - MUDS”. LV acknowledges funding by the Helmholtz Association’s Initiative and Networking Fund through CausalCellDynamics (grant # Interlabs-0029), and support from the Deutsche Forschungsgemeinschaft (DFG, Projektnummer 452881907, CRC TRR 338). GS acknowledges support by NIH Awards 5R01GM149631-03, R33CA286947-01A1 and R01CA297834-01. FJT acknowledges support from the Deutsche Forschungsgemeinschaft (DFG, Projektnummer 452881907, CRC TRR 338) and the European Union (ERC, DeepCell – 101054957).

## Author contributions

Conceptualization: LO, SB, LV, EP, GG, CMR, ABR, GS, FJT

Conducted laboratory experiments: SM, BH, EP, CC, AS, GG, JP, MN, OS, HN, VKT, AS, OK, SS, AS, GGO

Data analysis: LO, SB, LV, EP, JP, RK, AAM, JL, AL, MS

Visualization: LO, SB, LV, AAM, AL

Supervision and funding acquisition: CMR, ABR, GS, FJT

Writing original draft: LO, SB, LV, AAM, AL, MS

Writing, review and editing: LO, SB, LV, GS, FJT

## Corresponding authors

Correspondence to, and

## Competing interests

FJT consults for Immunai, CytoReason, BioTuring, Genbio and Valinor Discovery, and has ownership interest in RN.AI Therapeutics, Dermagnostix, and Cellarity. GS, CMR and ABR are co-founders of Parse Biosciences. The remaining authors declare no other competing interests.

## Notes

https://www.parsebiosciences.com/datasets/10-million-human-pbmcs-in-a-single-experiment/

https://github.com/theislab/HumanCytokineDict

https://github.com/theislab/huCIRA

