## Supplementary Information for "A single-cell cytokine dictionary of human peripheral blood"

[†] correspondence:

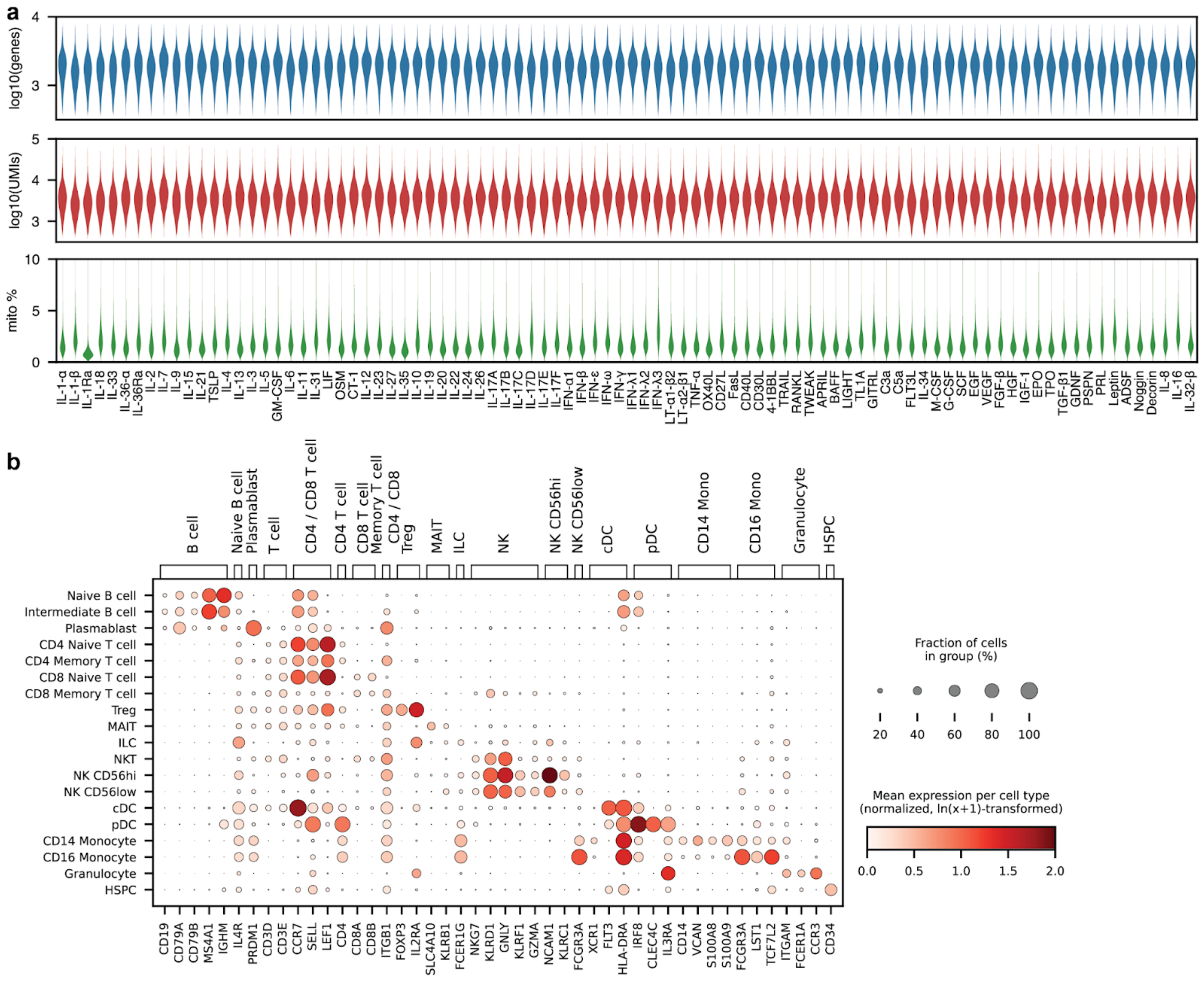

**Fig. S1. Quality control metrics and marker gene expression. a,** Distribution of the number of detected genes, total UMIs, and percent mitochondrial genes across all cells for a given cytokine perturbation after QC steps. **b,** Expression of the marker genes used to annotate cell types.

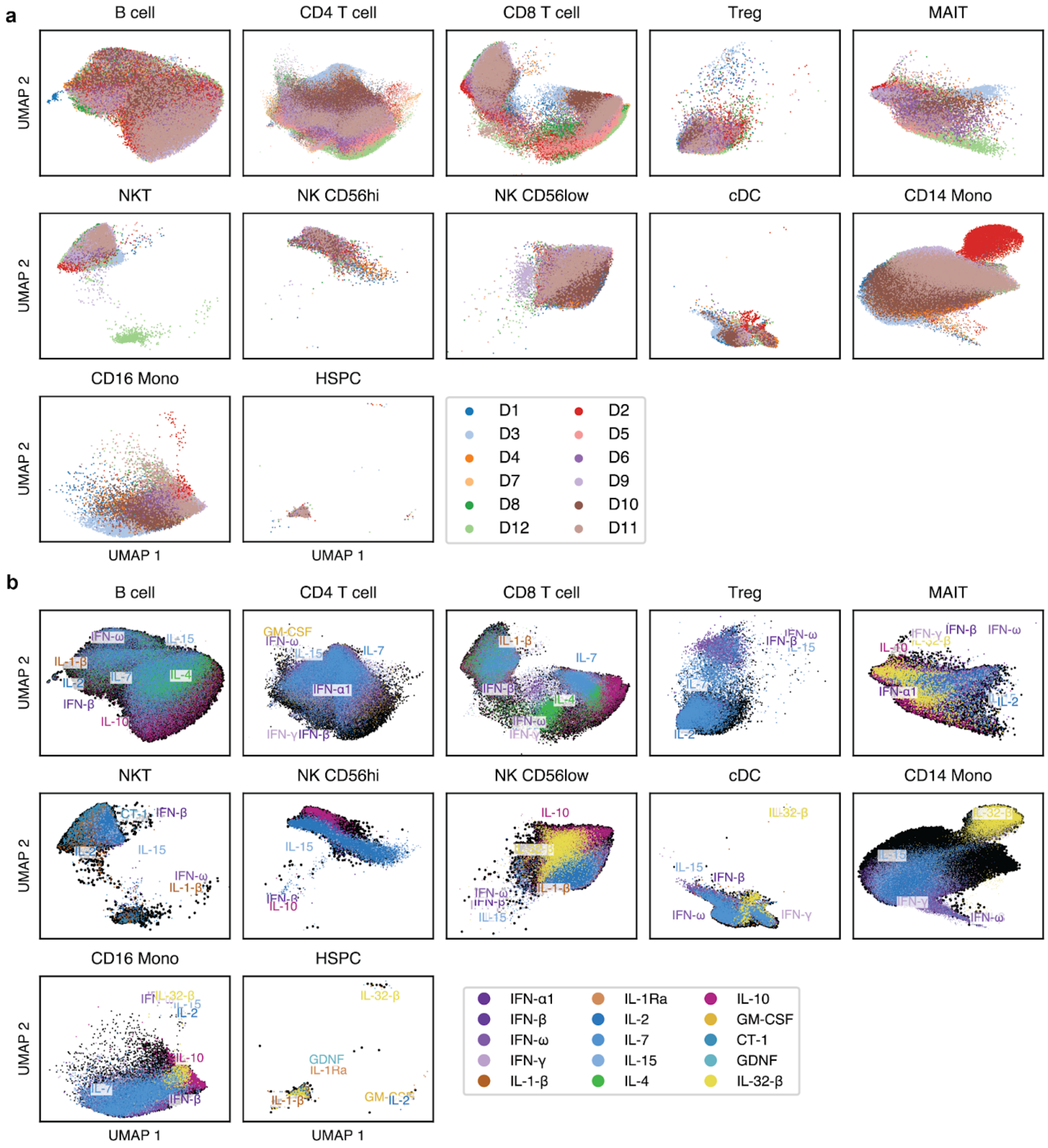


**Fig. S2. UMAP of cell type populations by donor and cytokine stimulation. a,b,** UMAP of the cell type populations displaying (**a**) the PBS condition by donor and (**b**) the cytokines causing the strongest shift in the position of the population for different cell types. Although strong cytokine perturbations shift cells in the UMAP space, cells overall cluster more strongly by cell type rather than by donor or perturbation.

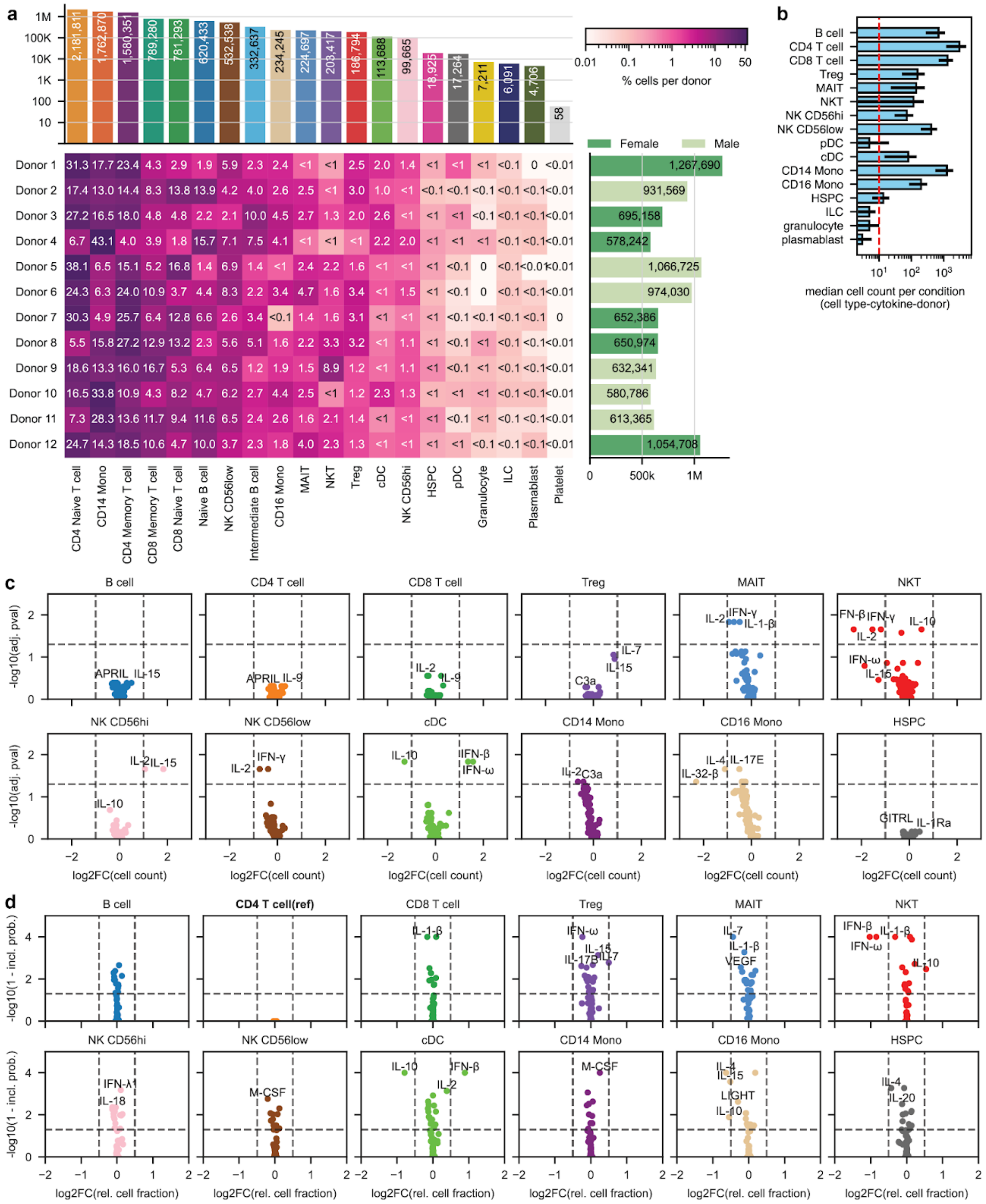


**Fig. S3. Cell type abundance analysis. a,** Fraction of counts per cell type for each donor. **b,** Median cell count per condition. The dashed red line shows the cutoff of 10 cells per condition (cell type-cytokine-donor triplet) we used to exclude cell types from the analysis. **c,** Volcano plot of differential cell count analysis (absolute numbers) between cytokine perturbations and the PBS controls. **d,** Bayesian compositional modeling (scCODA) using CD4 T cells as reference. The y axis plots the posterior probability of no effect.

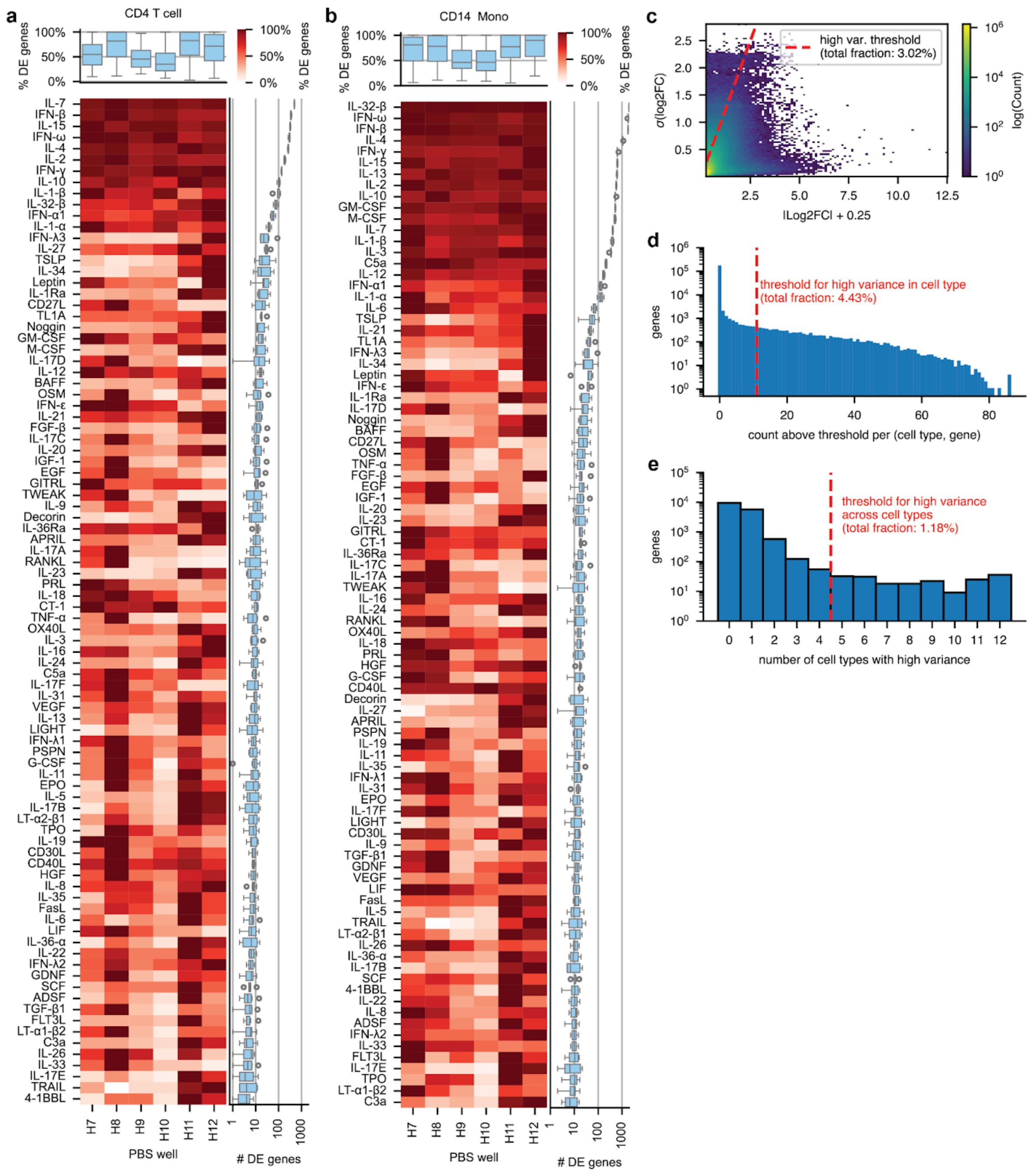


**Fig. S4. Rationale and process for additional DEG filtering based on consistency across wells. a, b,** Relative number of DEGs (|Log2FC|>0.25, padj<0.05) obtained for different PBS wells in (**a**) CD4 T cells and (**b**) CD14 monocytes. The box plot on top shows the distribution of relative values by well and the box plot on the right shows the distribution of absolute numbers by cytokine. DEGs used in the main text are filtered for consistency across wells. **c,** Standard deviation of log2FCs for a particular gene, cell type, and cytokine perturbation across wells versus their mean. Some genes are highly variable across wells (dashed red lines) **d,** Number of times a certain gene is above the high variability threshold shown in (**c**) in a particular cell type. Those genes that are highly variable for at least 11 cytokine treatments are considered highly variable in that cell type. **e,** The number of times a gene is highly variable in a given cell type. Any gene that is highly variable in at least 5 cell types is considered highly variable across cell types and filtered from the DEG analysis.

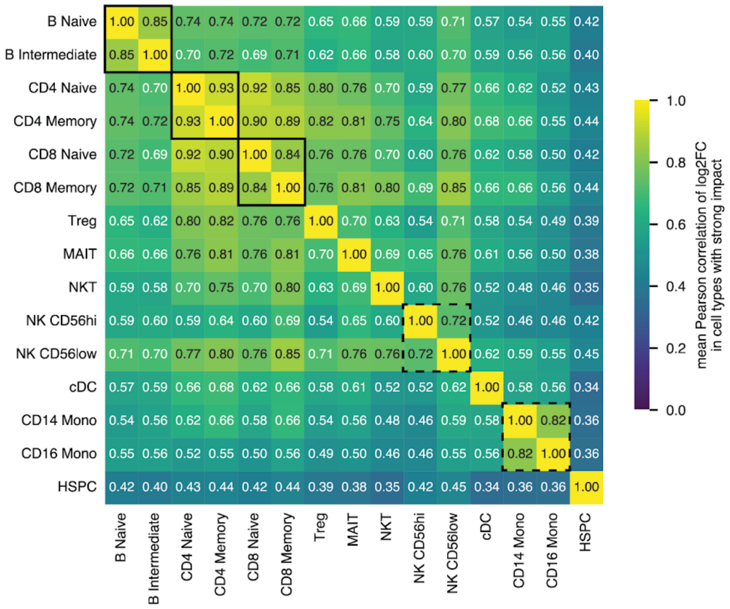


**Fig. S5. Merging of cell types.** Mean Pearson r between log2FC values in cell types with a strong impact. Solid boxes show cell types that were merged, dashed boxes cell types that could be considered for merging but were kept separate. While these cell types are merged for the analysis in the main text, we provide pseudobulk data and DEGs both for the merged cell type and individual cell types for all boxes.

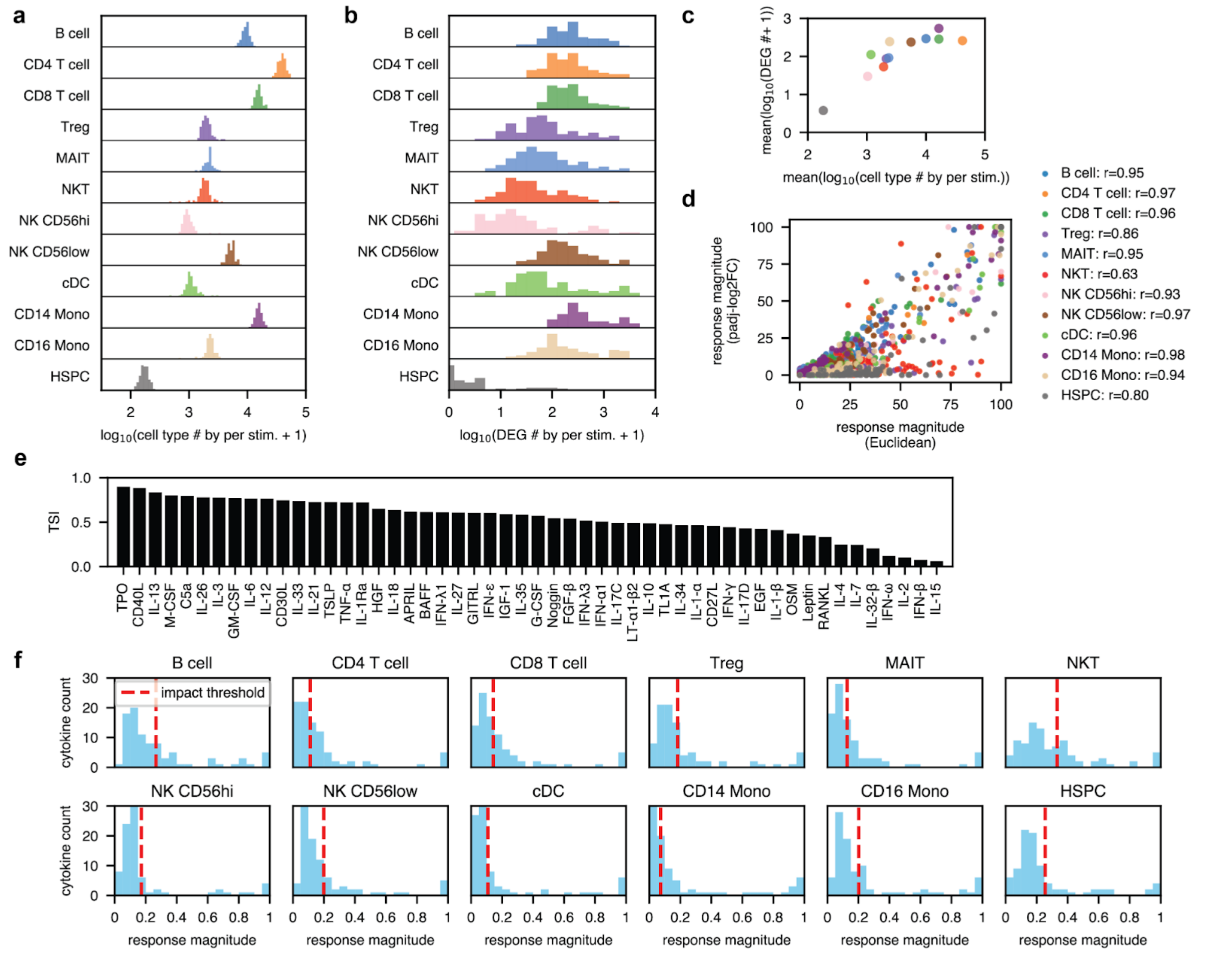


**Fig. S6. Cytokine DEG counts, response magnitudes, the tissue-specificity index, and strong impact classification. a**, **b**, Distribution of the total number of cells (**a**) and DEGs by cell type (**b**) across all cytokine perturbations before log2FC filtering. **c**, Association between the mean of the values in (**a**) and (**b**). **d,** Two measures of the response magnitude (Euclidean distance and padj-log2FC-based) are highly correlated in all cell types. **e,** Tissue-specificity index of the response magnitude for cytokines with at least one strong response. **f,** Distribution of the overall response magnitude (mean of Euclidean and padj-log2FC-based) across the cell types. The dashed red line shows the response magnitude threshold used to define strong impact.

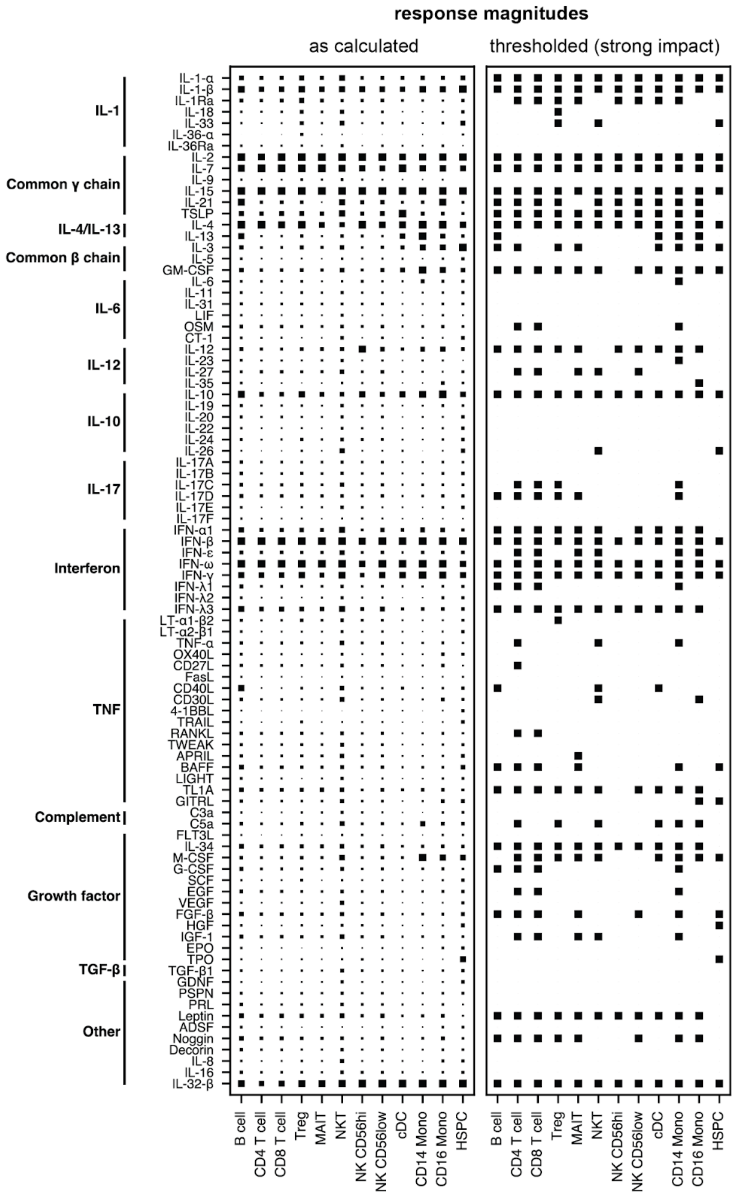


**Fig. S7. Response magnitude by individual cell types and cytokines.** Left: The calculated response magnitude is linearly proportional to the box size. Right: The presence of a box indicates a strong response.

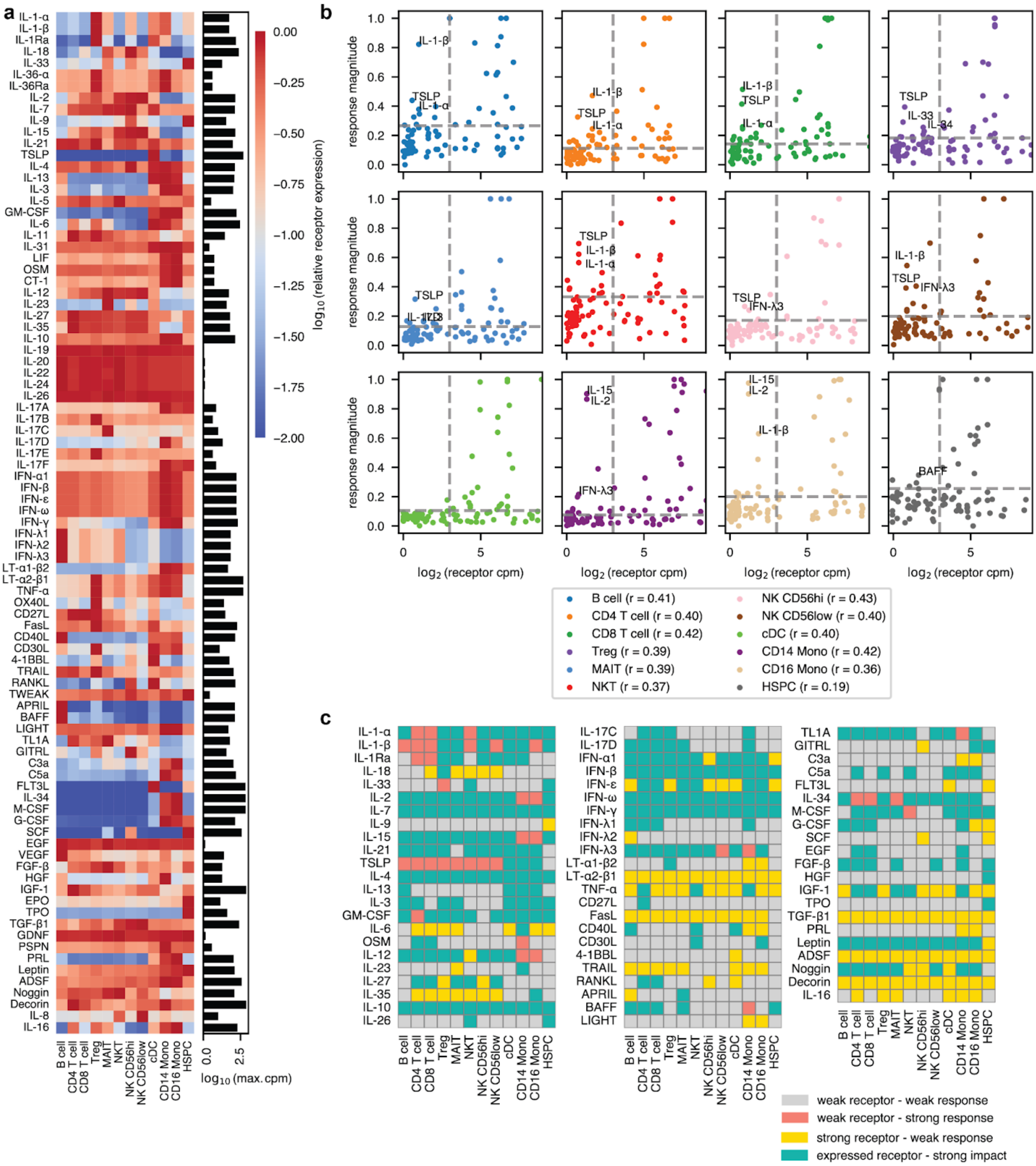


**Fig. S8. Cytokine receptor expression and its association with the response magnitude. a,** Relative receptor expression (cpm divided by the maximum across cell types) for all cytokines with known receptors (i.e., every tested cytokine except IL-32-β). The y-axis label shows the name of the cytokine for which the receptor expression is shown. The bar plot on the right shows the maximum cpm for each receptor. **b,** Response magnitude versus absolute receptor expression for all cytokines and cell types. The response magnitude threshold for strong activity is shown as a horizontal dashed line, and a receptor expression of 8 cpm is shown as a vertical dashed line and acts as the approximate threshold for functional receptor expression. For each cell type, a number of cytokines with a strong response despite low receptor expression are highlighted by name. **c,** Some cytokines show weak responses despite strong receptor expression (>32 cpm, response magnitude < threshold). Others show strong responses despite low receptor expression (<5 cpm, response magnitude > 1.5x threshold). We additionally annotate those with expressed receptors and a strong response (>8 cpm, response magnitude > threshold) and those with weak receptor expression and a weak response (<8 cpm, response magnitude < threshold).

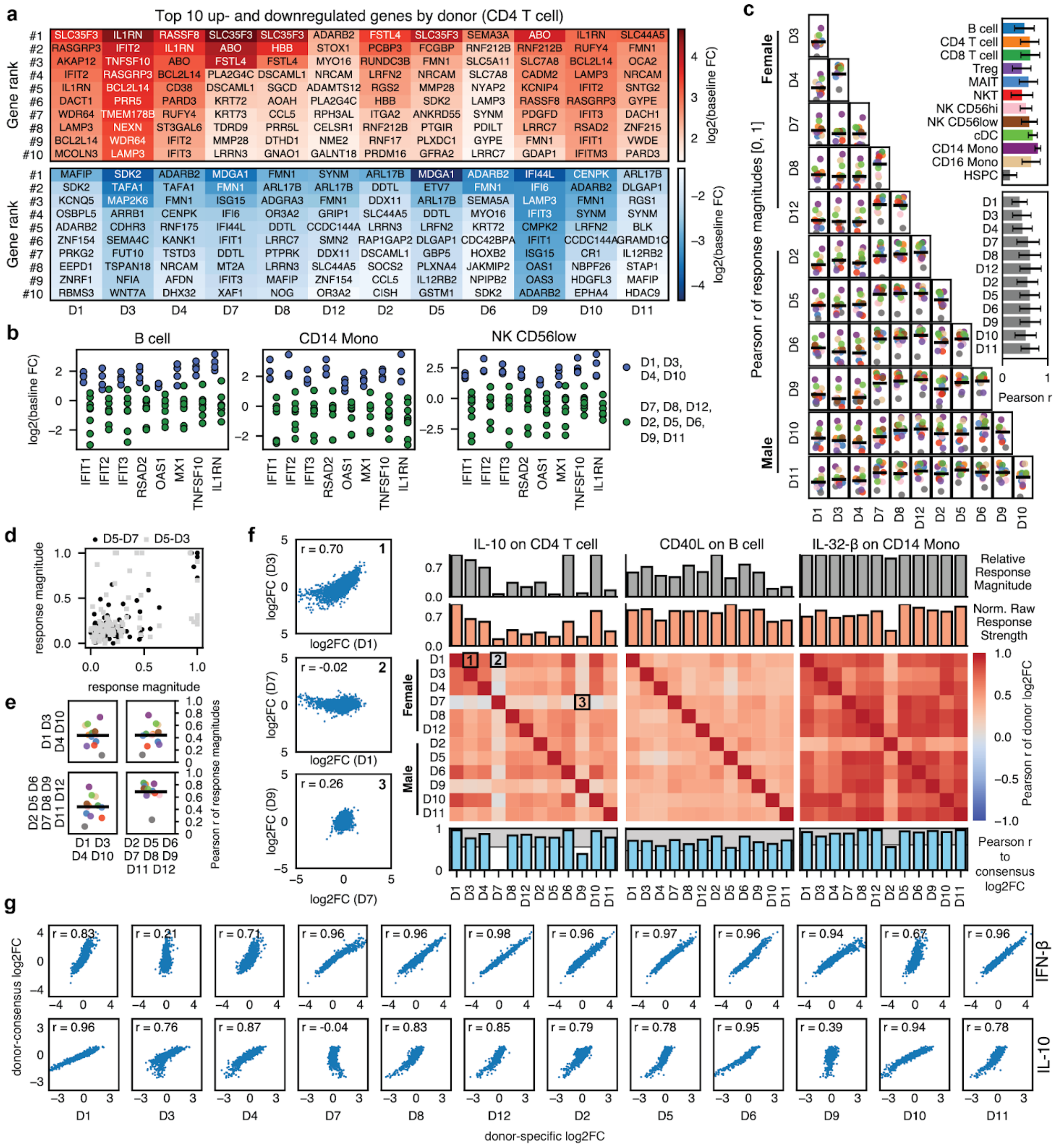


**Fig. S9. Additional analysis of donor variability in cytokine responses. a,** Top up- and downregulated genes in the baseline FC for each donor. Genes were pre-filtered to exclude ribosomal and mitochondrial genes, sex chromosome-linked genes, and non-protein coding genes. **b,** Baseline log2FC of interferon response genes for two groups of donors and three cell types. **c,** Pearson r of response magnitudes for all 90 cytokines within a given cell type between two donors. The solid line shows the mean across all cell types. Bar plots on the right show the marginal means across cell types and donors. **d,** Correlation of the response magnitudes for two sets of donors. **e,** As in (**a**) but averaged over the two indicated groups of donors. **f,** Correlation of donor-specific log2FC scores for different cell types and cytokines. The topmost barplot shows a response magnitude calculated using the Euclidean distance of vectors. The second barplot from the top shows a noise-robust measurement of the total response strength, normalized to the strongest response across donors for the shown condition. The bottom barplot shows the correlation to the donor-consensus log2FCs. The shaded gray area denotes a region around the median which we interpret to mean that the response is consistent, whereas responses outside this region are considered outliers. **g,** Donor-specific log2FC versus donor-consensus log2FC for two cytokines (IL-10 and IFN-β) and all donors. The Pearson r is shown on the top left of each plot.

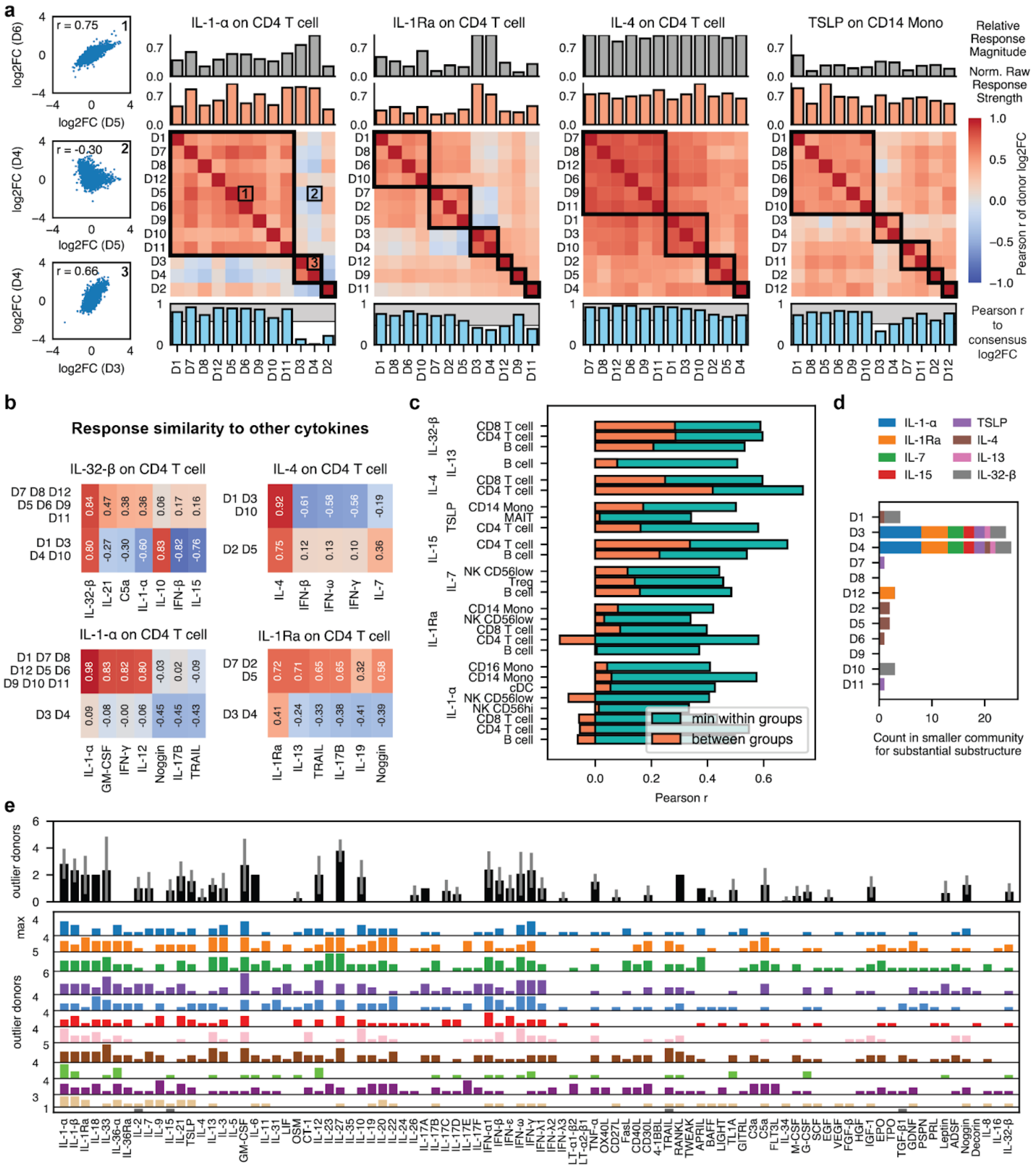

**Fig. S10. Detection of substantial sub-structure in donor responses. a,** Correlation of donor-specific log2FC scores for different cell types and cytokines. The topmost barplot shows a response magnitude calculated using the Euclidean distance of vectors. The second barplot from the top shows a noise-robust measurement of the total response strength, normalized to the strongest response across donors for the shown condition. The bottom barplot shows the correlation to the donor-consensus log2FCs. The shaded gray area denotes a region around the median which we interpret to mean that the response is consistent, whereas responses outside this region are considered outliers. Boxes indicate groupings of correlation patterns by Leiden clustering. **b,** Pearson r of the donor-group averaged log2FC for different cytokines and cell types (indicated in individual titles) compared to the most similar donor-consensus log2FC vectors across cytokines in the same cell type. **c,** Pearson r of donor group responses for cytokine-cell type pairs where substantial sub-structure was detected. We show the minimum of the two within-group comparisons and the between-group comparison. **d,** Presence of different donors in the minority group of responses with substantial substructure by cytokine. **e,** Count of donors with outlier responses for each cell type and cytokine. The maximum for each cell type is indicated on the left. The barplot on the top shows the mean across cell types.

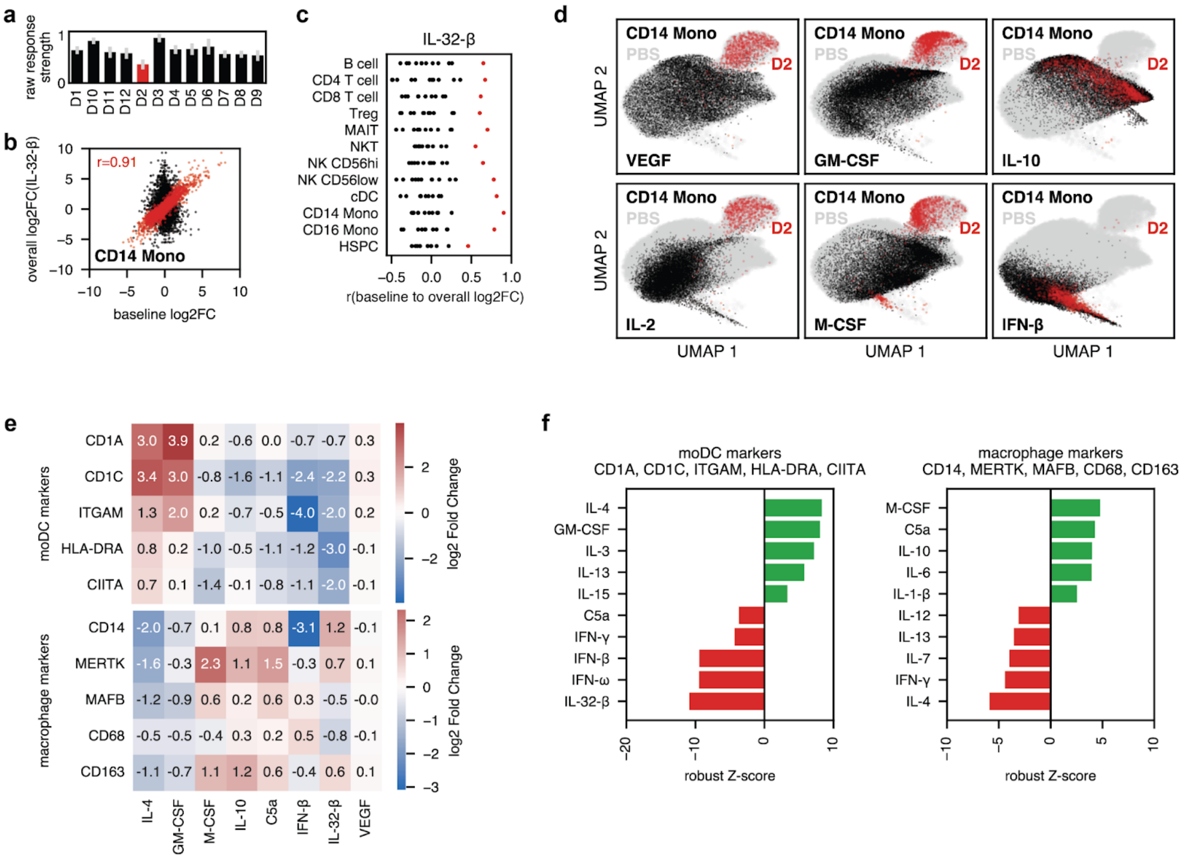


**Fig. S11. Donor 2 shows evidence of baseline IL-32-β signaling. a,** Raw response strength to IL-32-β by donor averaged over cell types. **b,** Baseline log2FC versus donor-consensus IL-32-β log2FC in CD4 T cells for either donor 2 or all other donors. **c,** Correlation between the baseline log2FC and the donor-consensus log2FC for IL-32-β by donor in all cell types. Donor 2 is highlighted in red. **d,** UMAP of CD14 monocytes in response to different cytokine perturbations. **e,** Up- and downregulation of macrophage (top) and moDC (bottom) identity markers by several relevant cytokines. Log2 Fold Changes were calculated using across-donor cpm means for the perturbation condition divided by across-donor cpm means for the PBS condition because filtering criteria for the consensus (edgeR-derived) log2FC impede the calculation of robust z scores across perturbation conditions. **f,** Sum of robust z-scores (calculated across perturbation conditions) for the markers shown in (**e**) by cytokine.

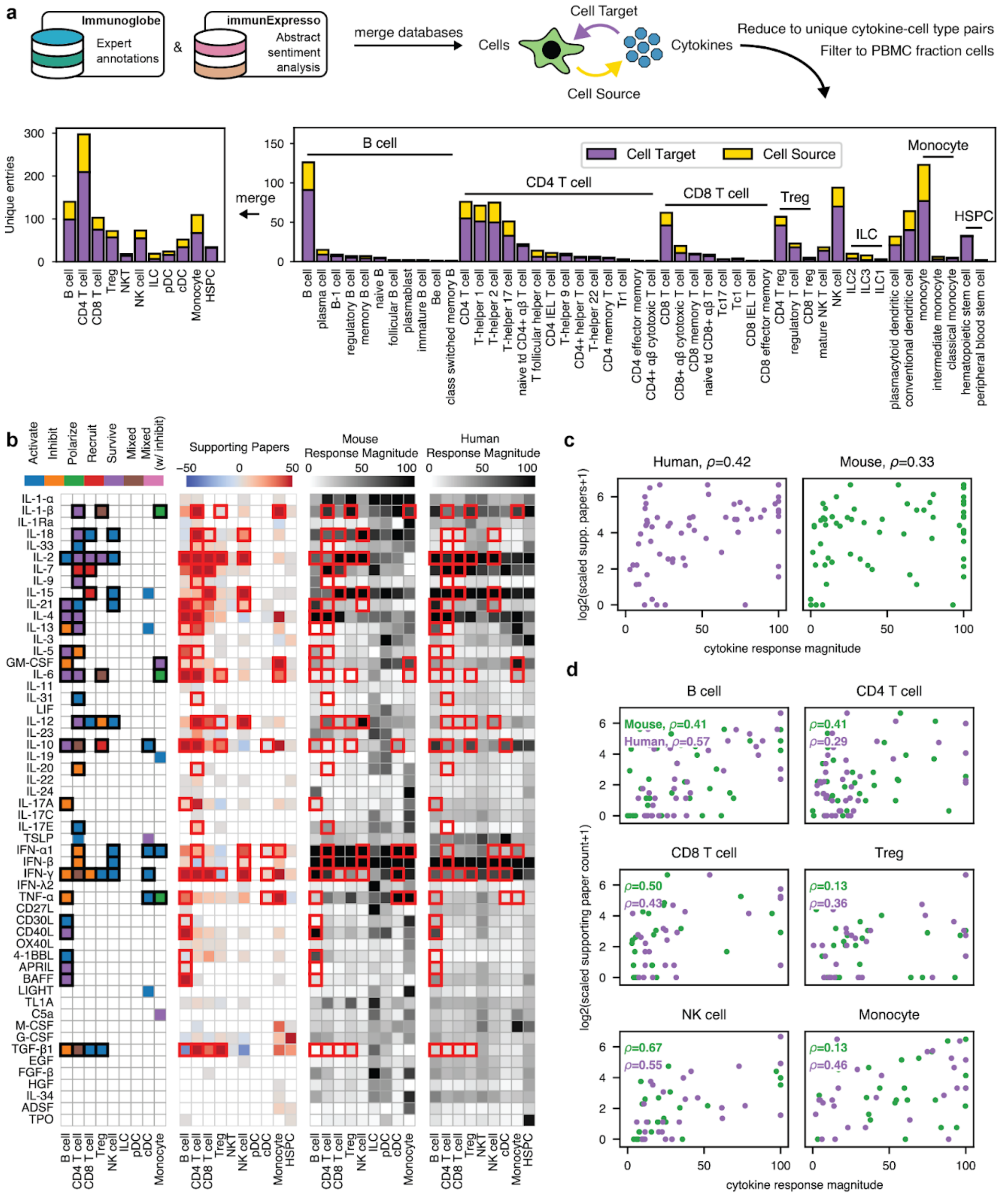


**Fig. S12. Database comparison with cytokine response magnitudes. a**, The construction of the reference dataset from the two input databases by merging on cell ontology identifiers. **b**, All relevant annotations for cytokine effects on specific cell types. From left to right, these include ImmunoGlobe (the colors in the legend represent the authors' annotation, with the two mixed categories being any combination of the other labels with or without inhibition included), ImmuneXpresso (where paper counts were considered positive if there were more positive labels per the relationship than negative, or negative otherwise), mouse cytokine response magnitude data for relevant cytokines, and our response magnitude values. Boxed cells are high-quality annotations. For the human data, we used NK CD56low to represent NK cells and CD14 Mono to represent monocytes as these are by far the majority populations. **c,** Correlation between mouse or human response magnitudes and scaled paper counts across all cell types by species. **d,** Correlation between mouse or human response magnitudes and scaled paper counts for each cell type.

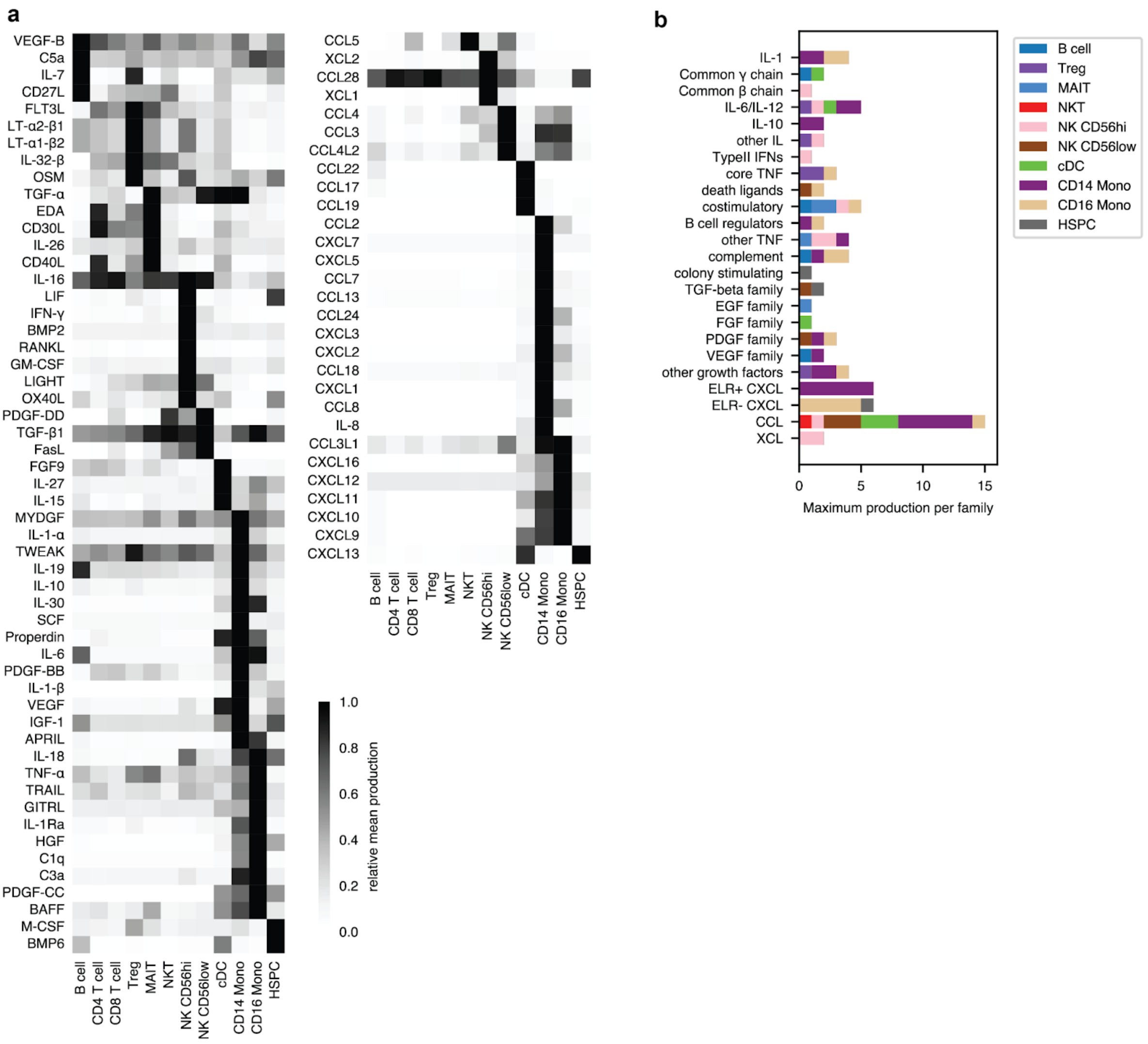


**Fig. S13. Expression of individual cytokines by cell type across stimulation conditions. a,** Mean production of cytokines across stimulation conditions for all cytokines expressed in at least one cell type. **b,** Number of times a given cell type has the strongest production of cytokines and chemokines of different families.

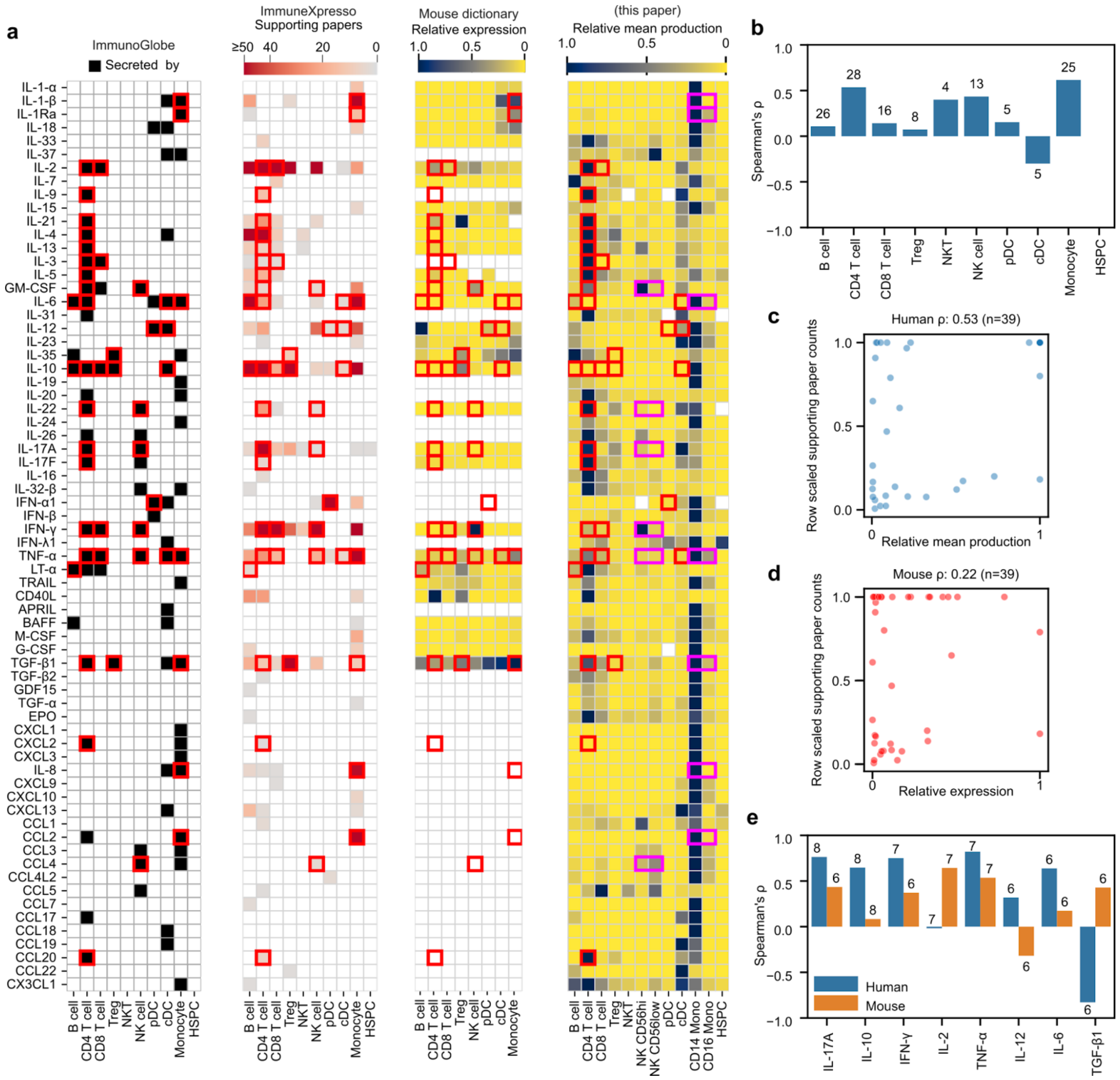


**Fig. S14. Cytokine secretion reference dataset comparisons. a,** All cytokine secretion values for immunoGlobe, ImmuneXpresso, relative expression mouse cytokine dictionary information, and relative mean expression (this publication) from left to right. Note that the boxed values are those shared between databases, with the purple two by one boxes denoting cell types which were simplified for the end analysis. **b,** Correlation of relative mean expression values with row-scaled paper counts per cell type for all annotations with paper count information. Numbers are n for cells with secretion data. **c,** Human relative mean production vs row scaled paper counts (see methods) for only high quality annotations shared between the mouse and human datasets. **d,** Mouse relative expression vs row scaled paper counts for only high quality annotations shared between the mouse and human datasets. **e,** Per-cytokine correlations of relative expression and production with row scaled supporting paper counts for well-studied cytokines.

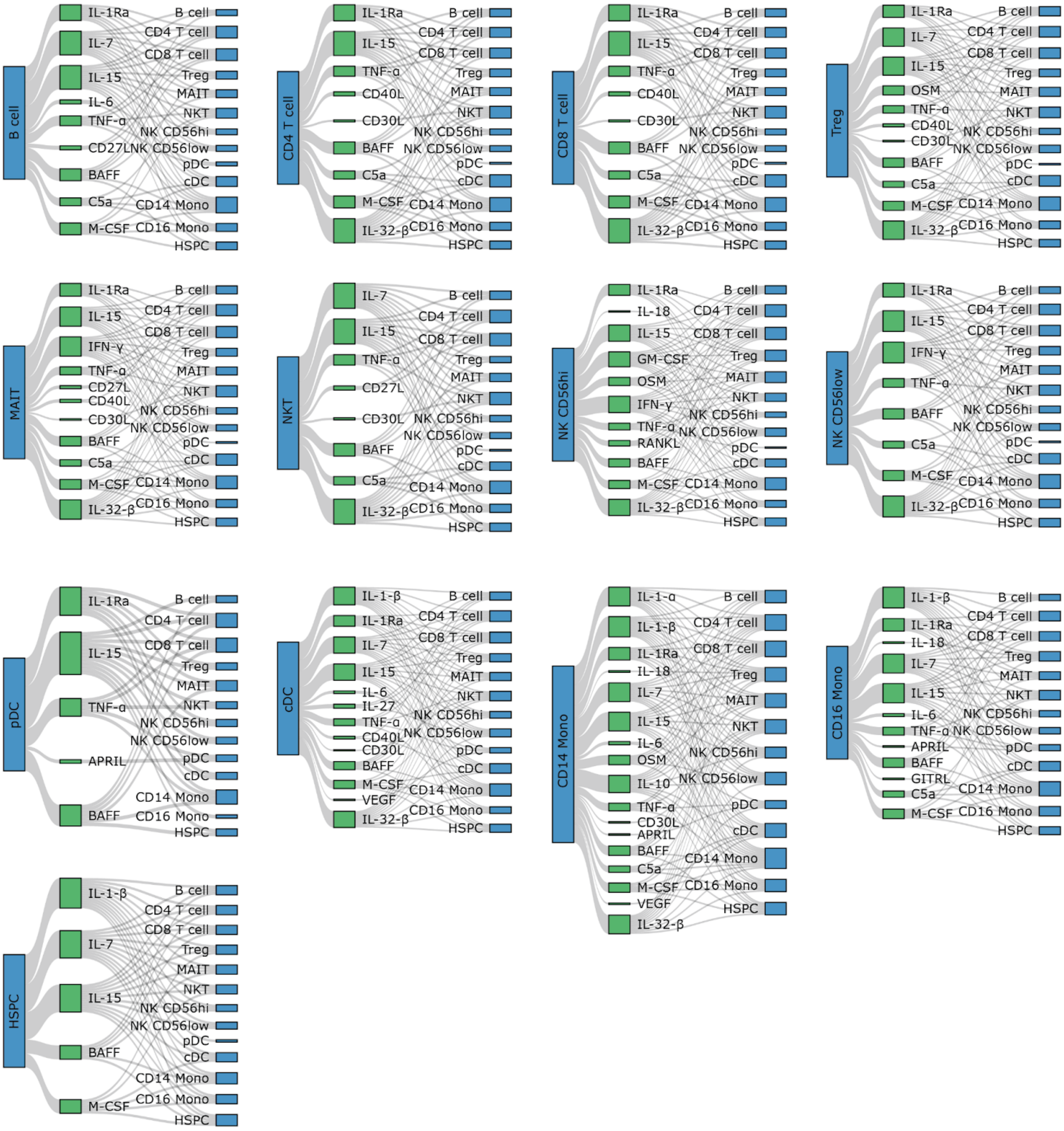


**Fig. S15. Cytokines secreted by a given cell type cause a strong impact in other cell types.** Each connection shows expression of a cytokine by a cell type that fulfills the strong impact criterion in the target cell type.

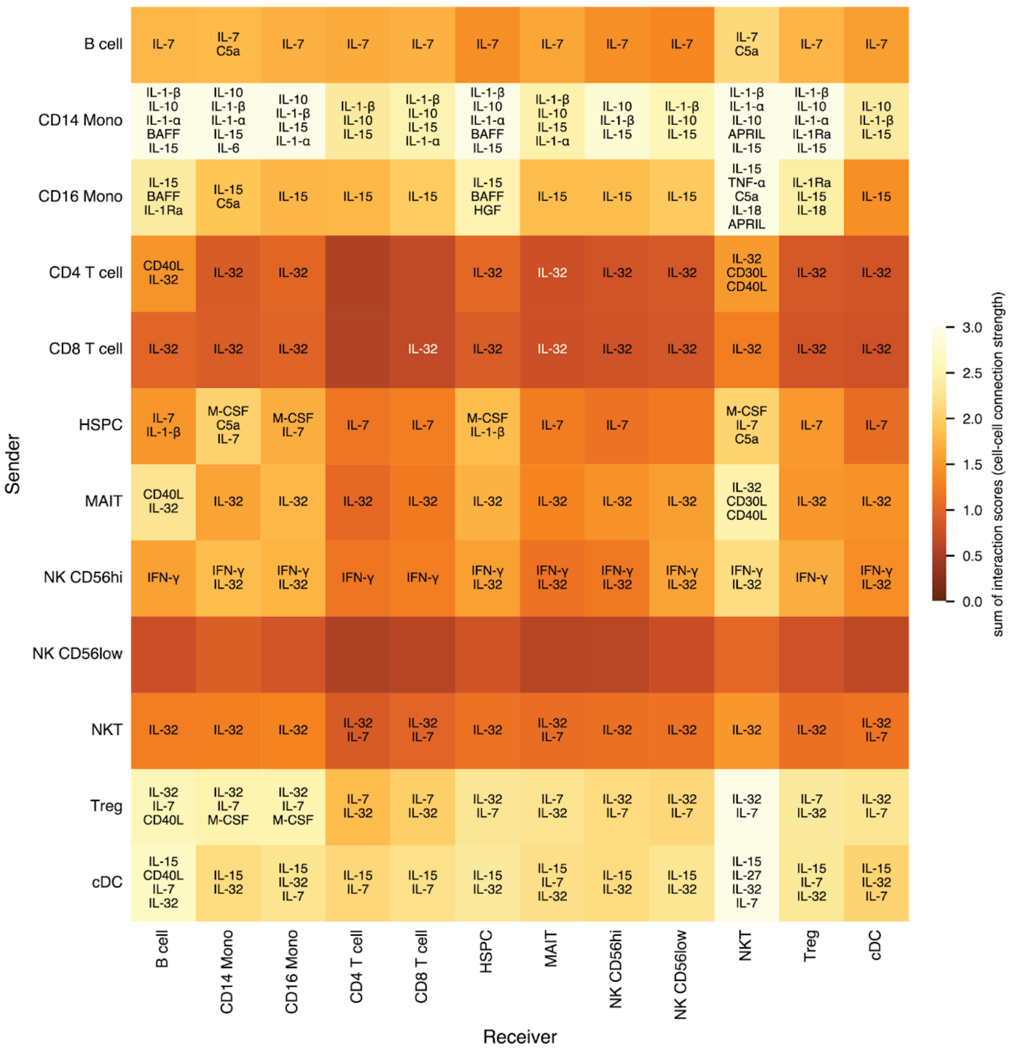


**Fig. S16. Cell-cell communication map by specific cytokines.** For each sender cell-cytokine-target cell combination, an interaction score was calculated as the product of the relative production of that cytokine in the sender cell type compared to other cell types times the response magnitude of the cytokine in the target cell type. An overall cell type-cell type connection strength is calculated as the sum of their interaction scores. Cytokines with an interaction score above 0.25 are annotated.

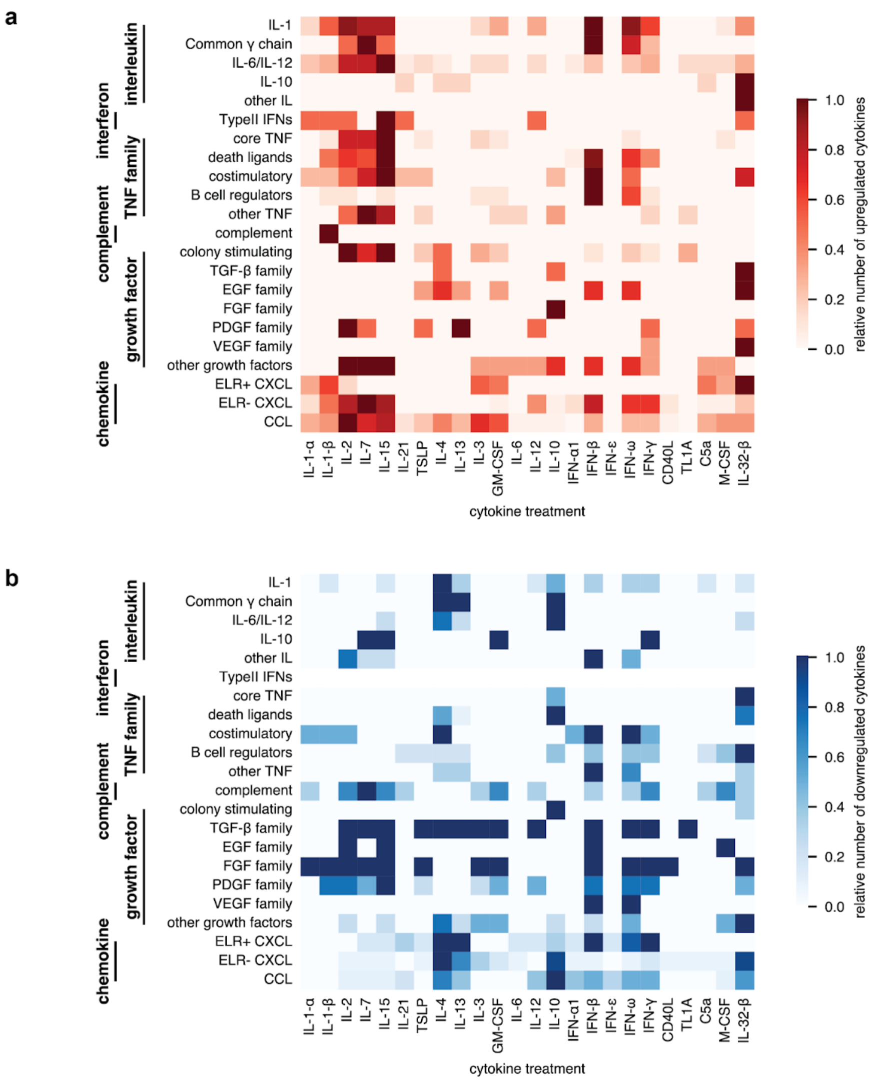


**Fig. S17. Cytokine-cytokine family crosstalk. a,** Up- and **b,** downregulation of different cytokine families by cytokine treatments. The figure sums regulation in the different cell types and then divides by the maximum value for a given family to derive a relative number.

| produced cytokine | cell type | treatment | mean cpm | FC to PBS |
|---|---|---|---|---|
| APRIL | CD16 Mono | IL-3 | 87 | 1.8 |
| BAFF | CD14 Mono | IFN-β | 718 | 3.2 |
| BMP6 | cDC | IL-32-β | 170 | 8.9 |
| C1q | CD16 Mono | IFN-γ | 105 | 1.7 |
| C3a | CD14 Mono | IL-32-β | 75 | 2.9 |
| C5a | B cell | IL-12 | 42 | 1.2 |
| CCL13 | CD14 Mono | IL-4 | 135 | 5.0 |
| CCL17 | cDC | IL-4 | 570 | 28.2 |
| CCL18 | CD14 Mono | IL-4 | 225 | 15.5 |
| CCL19 | cDC | IL-1-β | 172 | 3.8 |
| CCL2 | CD14 Mono | M-CSF | 2115 | 7.5 |
| CCL22 | cDC | TSLP | 2508 | 4.7 |
| CCL23 | CD14 Mono | IL-4 | 71 | 53.3 |
| CCL24 | CD14 Mono | IL-3 | 609 | 5.7 |
| CCL3 | NK CD56low | IL-15 | 570 | 11.8 |
| CCL3L1 | CD16 Mono | IL-32-β | 154 | 20.0 |
| CCL4 | NK CD56low | IL-15 | 317 | 5.9 |
| CCL4L2 | NK CD56low | IL-2 | 51 | 3.3 |
| CCL5 | NKT | IFN-ω | 731 | 1.3 |
| CCL7 | CD14 Mono | M-CSF | 363 | 32.4 |
| CCL8 | CD14 Mono | IL-2 | 967 | 7.4 |
| CD30L | MAIT | IL-7 | 127 | 1.6 |
| CD40L | MAIT | IL-15 | 150 | 4.8 |
| CXCL1 | CD14 Mono | C5a | 498 | 7.1 |
| CXCL10 | CD16 Mono | IL-15 | 4069 | 4.3 |
| CXCL11 | CD14 Mono | IFN-β | 1725 | 28.0 |
| CXCL13 | HSPC | IL-32-β | 978 | 30.3 |
| CXCL16 | CD16 Mono | IL-32-β | 236 | 1.7 |
| CXCL2 | CD14 Mono | IL-32-β | 134 | 5.7 |
| CXCL3 | CD14 Mono | C5a | 450 | 3.0 |
| CXCL5 | CD14 Mono | C5a | 2569 | 41.1 |
| CXCL7 | CD14 Mono | C5a | 474 | 37.2 |
| CXCL9 | CD16 Mono | IL-2 | 1972 | 10.1 |
| EDA | CD4 T cell | IL-7 | 240 | 1.9 |
| FGF9 | cDC | IL-10 | 45 | 4.0 |
| FLT3L | Treg | IL-24 | 136 | 1.4 |
| FasL | NK CD56low | IL-15 | 299 | 2.3 |
| GITRL | CD16 Mono | IL-21 | 61 | 9.4 |
| GM-CSF | NK CD56hi | IL-1-β | 53 | 9.0 |
| HGF | CD16 Mono | CD27L | 64 | 1.2 |
| IFN-γ | NK CD56hi | IL-1-β | 367 | 19.7 |
| IL-1-α | CD14 Mono | IL-32-β | 122 | 26.5 |
| IL-1-β | CD14 Mono | IL-32-β | 3287 | 46.7 |
| IL-10 | CD14 Mono | IL-32-β | 60 | 4.9 |
| IL-15 | cDC | CD40L | 1082 | 1.3 |
| IL-16 | NK CD56hi | IL-34 | 177 | 1.3 |
| IL-18 | CD16 Mono | IFN-ε | 56 | 1.3 |
| IL-1Ra | CD16 Mono | IFN-β | 1038 | 5.9 |
| IL-24 | CD14 Mono | IL-32-β | 52 | 71.2 |
| IL-27 | cDC | IFN-γ | 43 | 5.3 |
| IL-30 | CD14 Mono | IFN-β | 131 | 7.3 |
| IL-32 | MAIT | IL-15 | 351 | 2.9 |
| IL-6 | CD14 Mono | C5a | 133 | 7.8 |
| IL-7 | B cell | CD40L | 244 | 1.6 |
| IL-8 | CD14 Mono | IL-32-β | 2299 | 35.1 |
| LIF | NK CD56hi | IL-15 | 104 | 5.8 |
| LIGHT | NK CD56hi | IL-1-β | 100 | 1.4 |
| LT-α1-β2 | MAIT | IL-7 | 125 | 5.5 |
| LT-α2-β1 | MAIT | IL-7 | 125 | 12.6 |
| M-CSF | HSPC | IL-24 | 206 | 1.0 |
| MYDGF | CD14 Mono | IL-8 | 66 | 1.4 |
| OSM | MAIT | IL-7 | 41 | 21.6 |
| OX40L | NK CD56hi | IL-32-β | 114 | 3.9 |
| PDGF-CC | CD16 Mono | IL-4 | 527 | 1.8 |
| PDGF-DD | NK CD56low | IL-4 | 462 | 1.2 |
| Properdin | CD16 Mono | GITRL | 95 | 0.9 |
| RANKL | NK CD56hi | CD30L | 59 | 1.5 |
| SCF | CD14 Mono | M-CSF | 89 | 10.8 |
| TGF-α | CD14 Mono | IL-4 | 190 | 5.6 |
| TGF-β1 | CD16 Mono | Leptin | 217 | 1.2 |
| TGF-β2 | HSPC | FGF-β | 40 | 23.6 |
| TNF-α | MAIT | IL-7 | 106 | 6.6 |
| TRAIL | CD16 Mono | IFN-β | 2193 | 3.1 |
| VEGF | CD14 Mono | IL-32-β | 114 | 7.9 |
| XCL1 | NK CD56hi | IFN-λ3 | 75 | 1.5 |
| XCL2 | NK CD56hi | TSLP | 45 | 1.6 |

**Fig. S18. Table of maximum expression values per expressed cytokine by cell type and stimulation condition.** We show the condition with the largest cpm value for a given cytokine, which doesn't necessarily imply a strong upregulation for that cytokine relative to baseline, as it might be expressed constitutively (cf. the FC to PBS column).

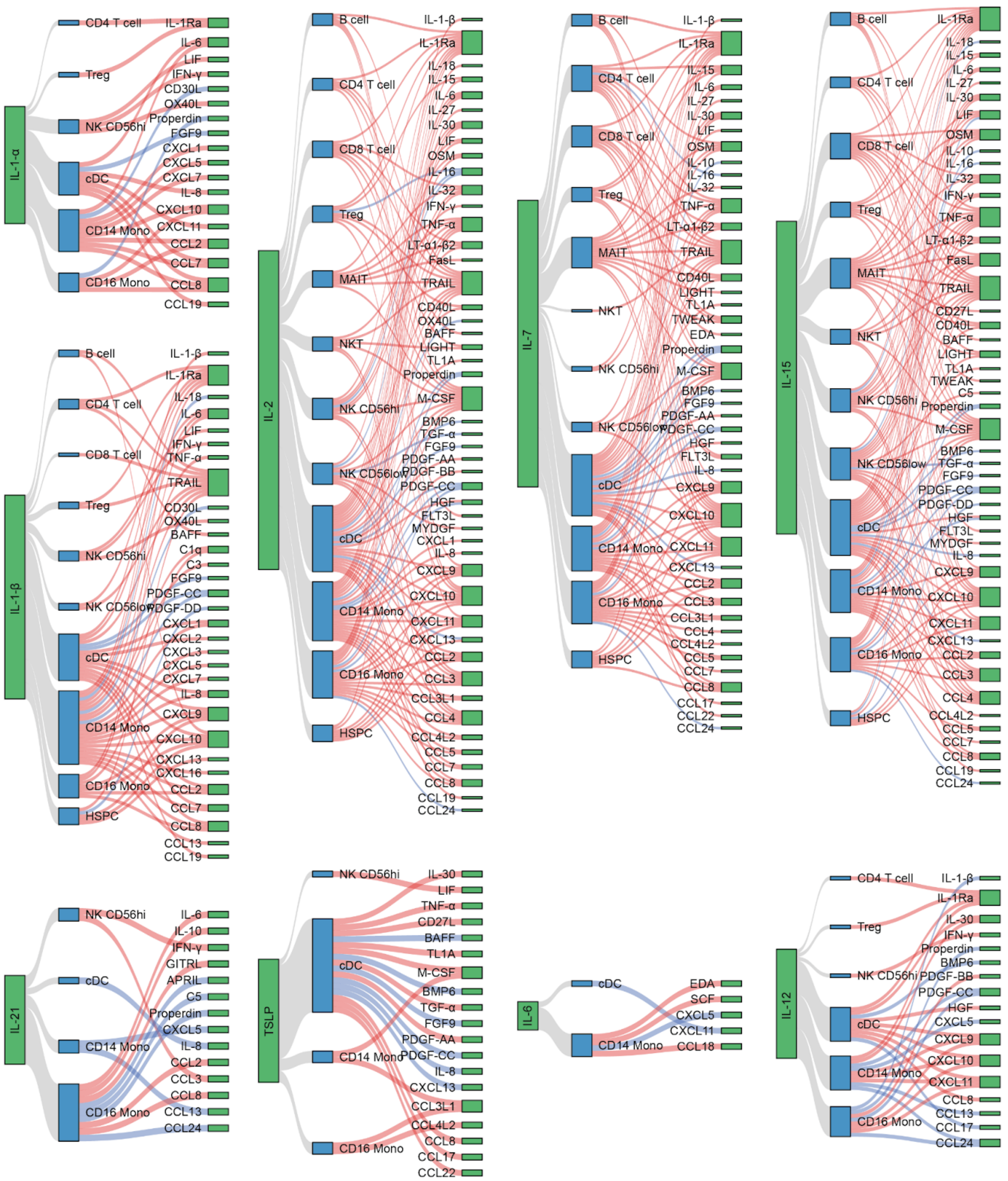


**Fig. S19. Cytokine-to-cytokine crosstalk (IL-1-β, IL-2, IL-7, IL-15, IL-21, TSLP, IL-6, IL-12).** Each connection shows regulation (FC > 2 in blue, FC <0.5 in blue) of a given cytokine by another cytokine in a given cell type.

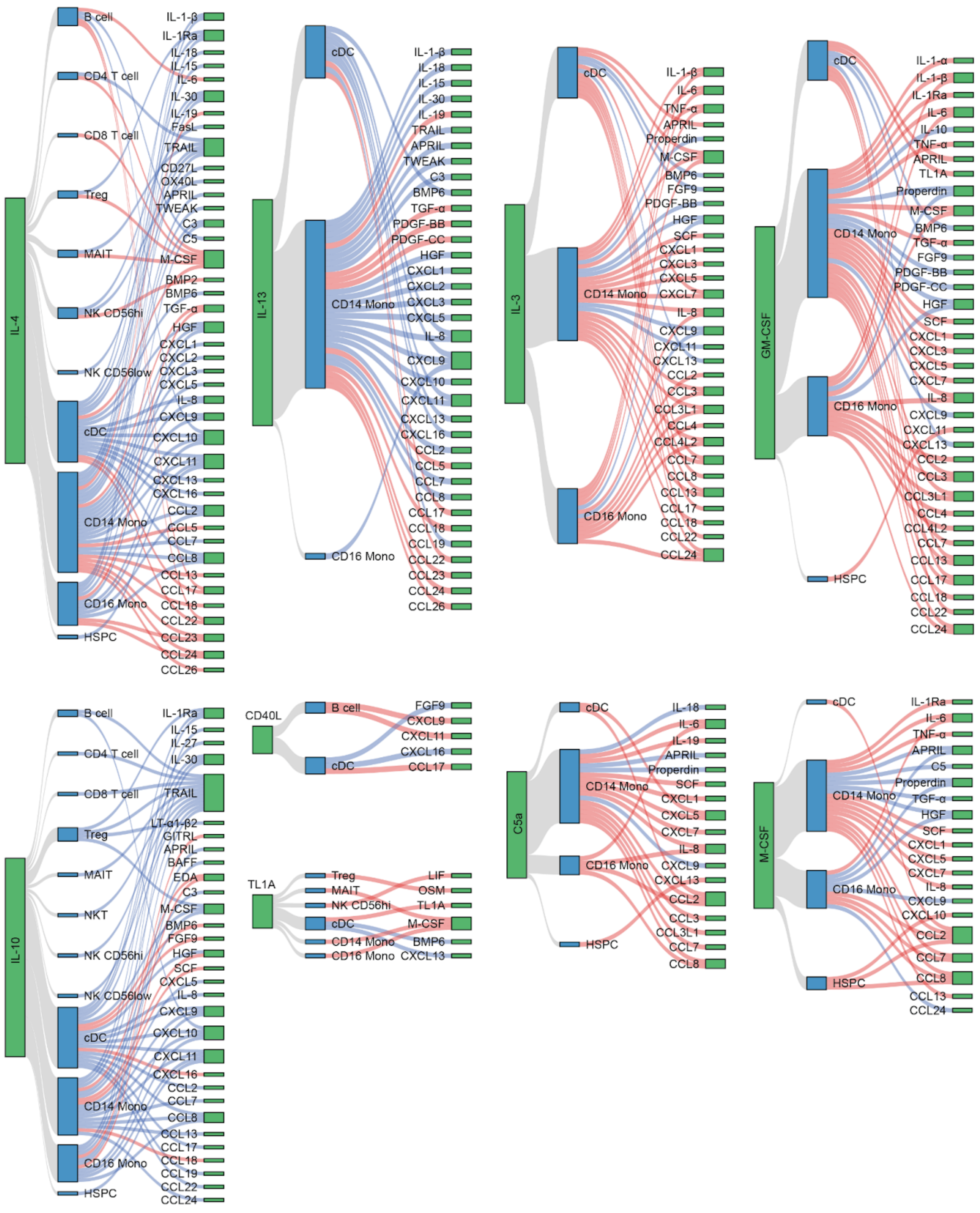


**Fig. S20. Cytokine-to-cytokine crosstalk (IL-4, IL-13, IL-3, GM-CSF, IL-10, CD40L, TL1A, C5a, M-CSF).** Each connection shows regulation (FC > 2 in blue, FC <0.5 in blue) of a given cytokine by another cytokine in a given cell type.

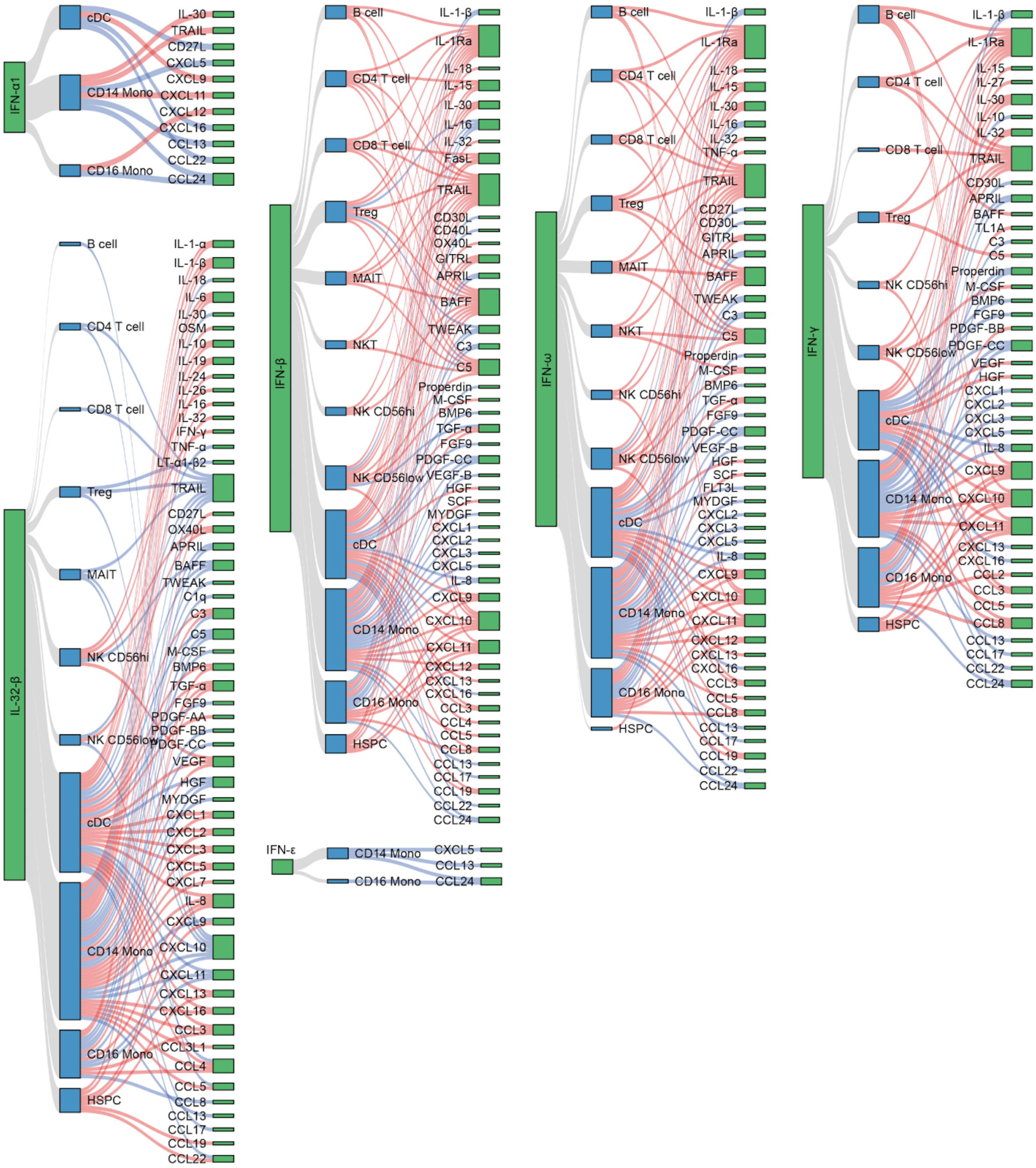


**Fig. S21. Cytokine-to-cytokine crosstalk (IFN-α1, IFN-β, IFN-ω, IFN-γ, IFN-ε, IL-32-β).** Each connection shows regulation (FC > 2 in blue, FC <0.5 in blue) of a given cytokine by another cytokine in a given cell type.

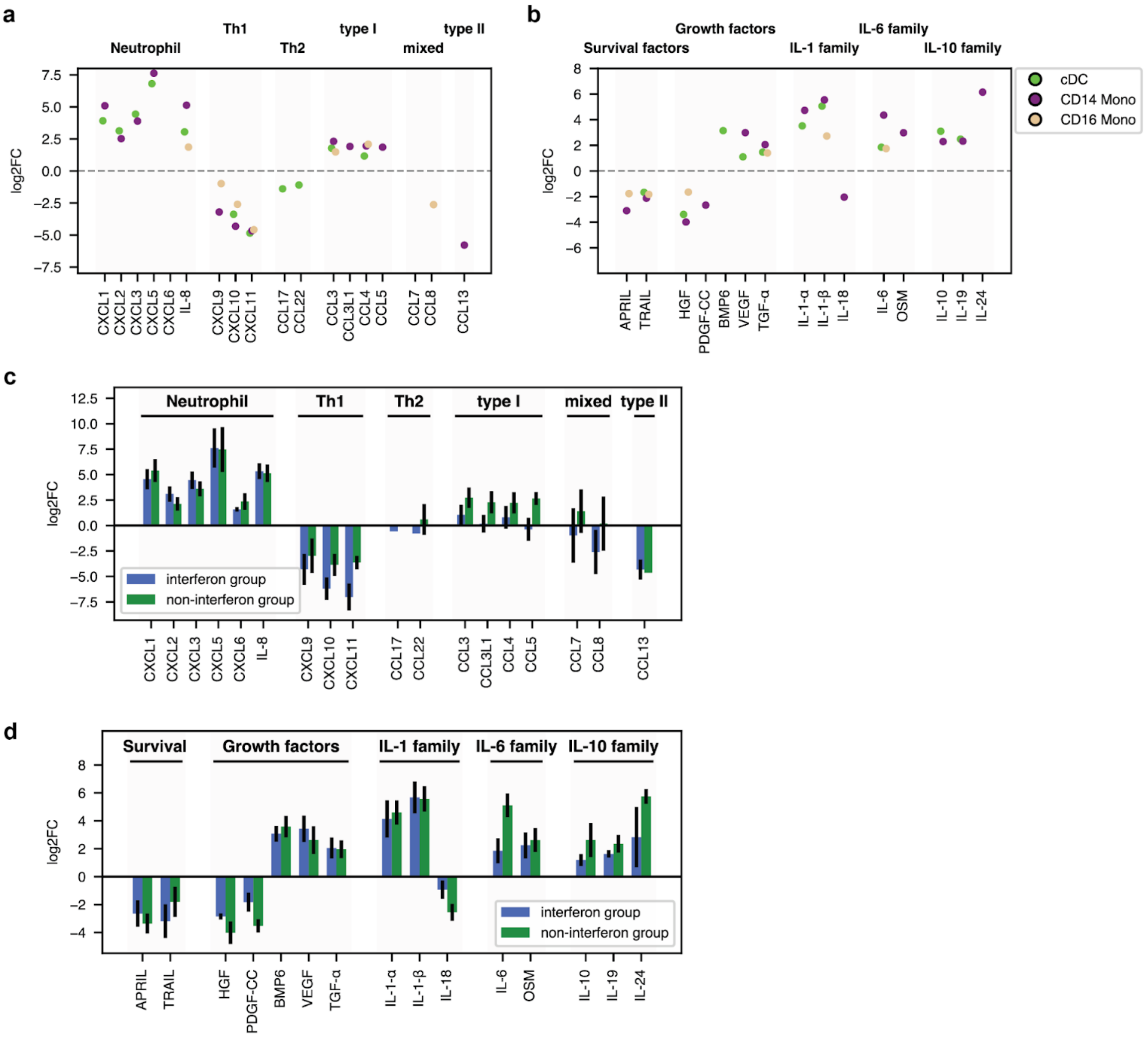


**Fig. S22. Cytokine signaling induced by IL-32-β in myeloid cells and by donor group. a,b,** Fold changes for chemokines (**a**) and other cytokines (**b**) grouped by cytokine family in myeloid cell types. Missing values indicate that the cytokine did not pass the filters used for DEG calculation. **c,d,** Fold changes for chemokines (**a**) and other cytokines (**b**) grouped by cytokine family in CD14 monocytes for two different groups of donors. See Fig. 2 for the list of donors associated with each group. Donor 2 was excluded from this calculation to the strong pre-existing IL-32-β signaling.

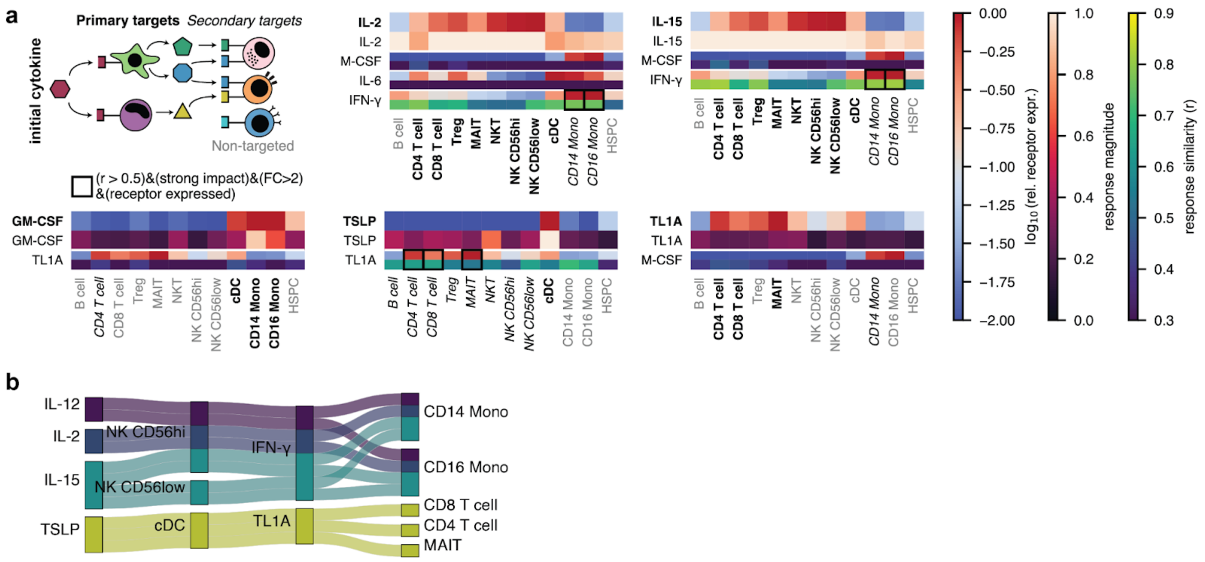

**Fig. S23. Exploration of potential secondary responses for different cytokines. a,** Potential secondary responses were detected by a combination of low receptor expression and high response magnitude in at least one cell type. We show all potential secondary cytokines, i.e., cytokines significantly upregulated (FC>2, padj<0.05) in at least one primary target cell type with receptor expression and a strong response in at least one secondary cell type, are shown. High response similarity (r>0.5) is used as an additional criterion to narrow down potential secondary signaling mechanisms (black box). **b,** IL-2, IL-12, and IL-15 all likely signal to secondary cells via IFN-γ released by (CD56-positive) natural killer cells. TSLP might use TL1A signaling via cDCs as a mechanism to affect T cells.

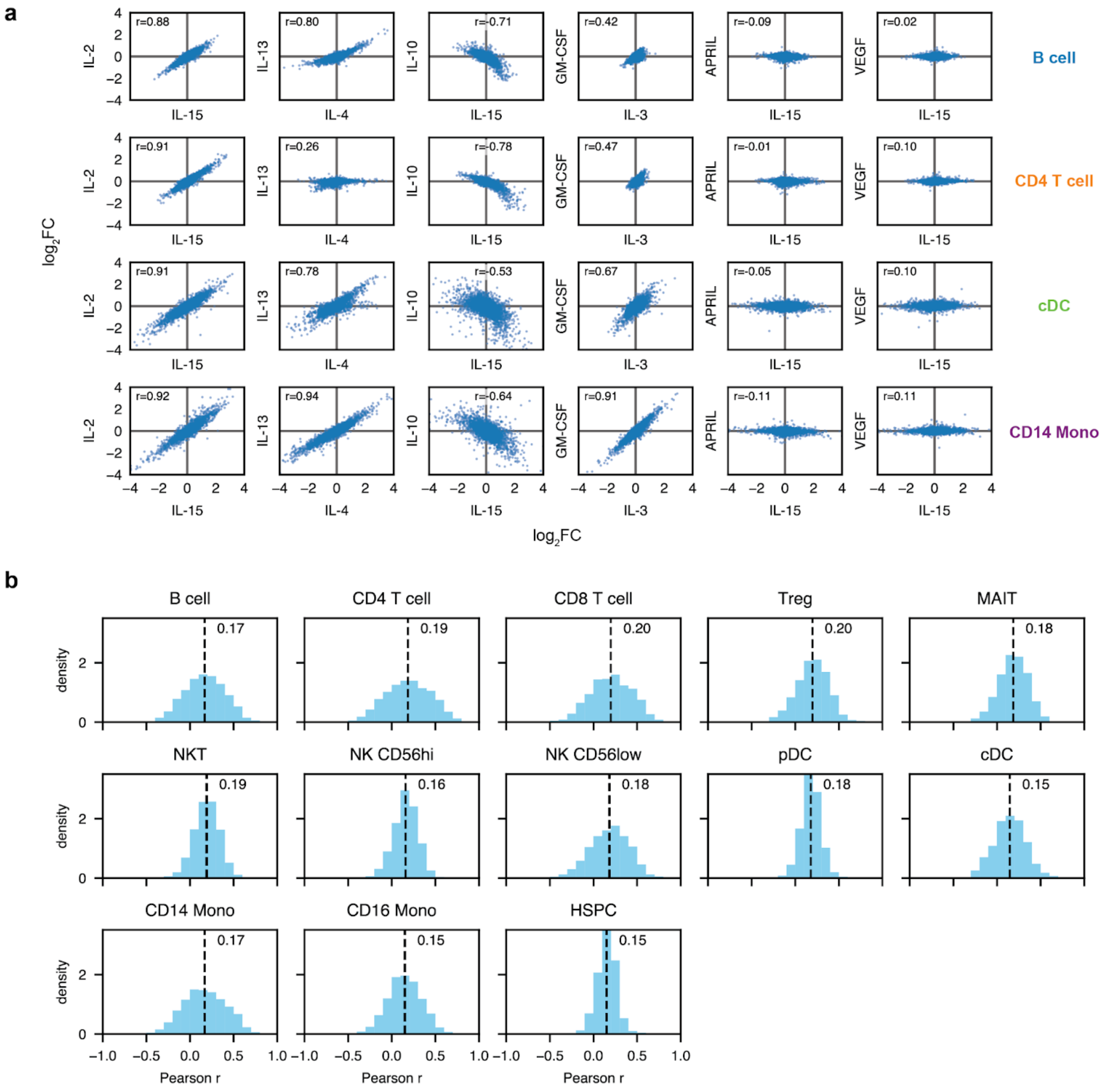


**Fig. S24. Cytokine-cytokine log2FCs correlations quantify response similarity. a,** Scatter plots of log2FC value correlations for different chosen cytokines in B cells, CD4 T cells, cDCs, and CD14 Monocytes. **b,** Distribution of Pearson r values between all cytokine pairs in different cell types. The dashed line and number show the mean value. There is a slight bias towards positive correlation.

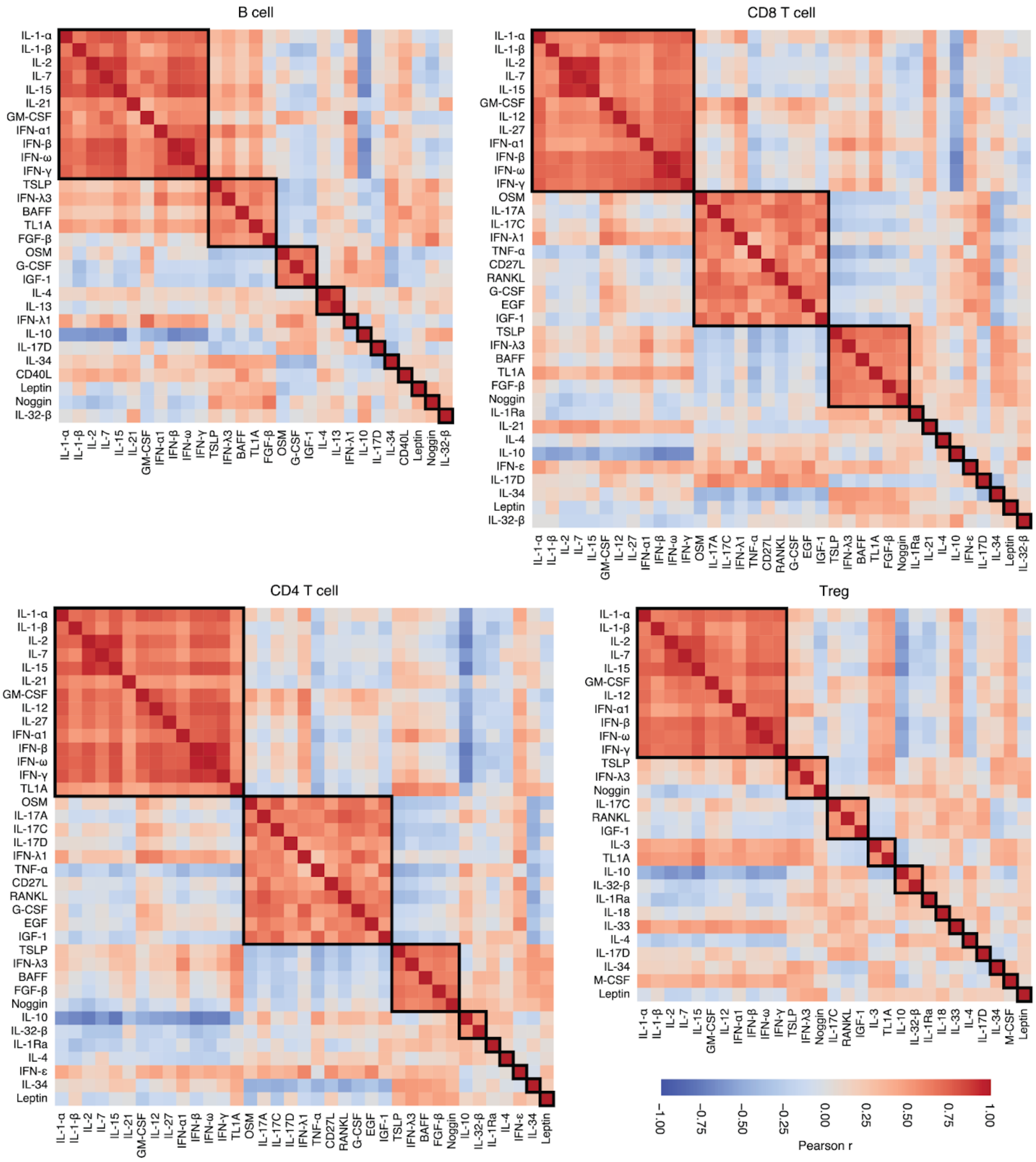


**Fig. S25. Correlation heatmaps between log2FC values for different cytokines in B cells, CD4 and CD8 T cells, and Tregs.** Cytokines were grouped based on their correlation patterns using the Leiden algorithm (black boxes).

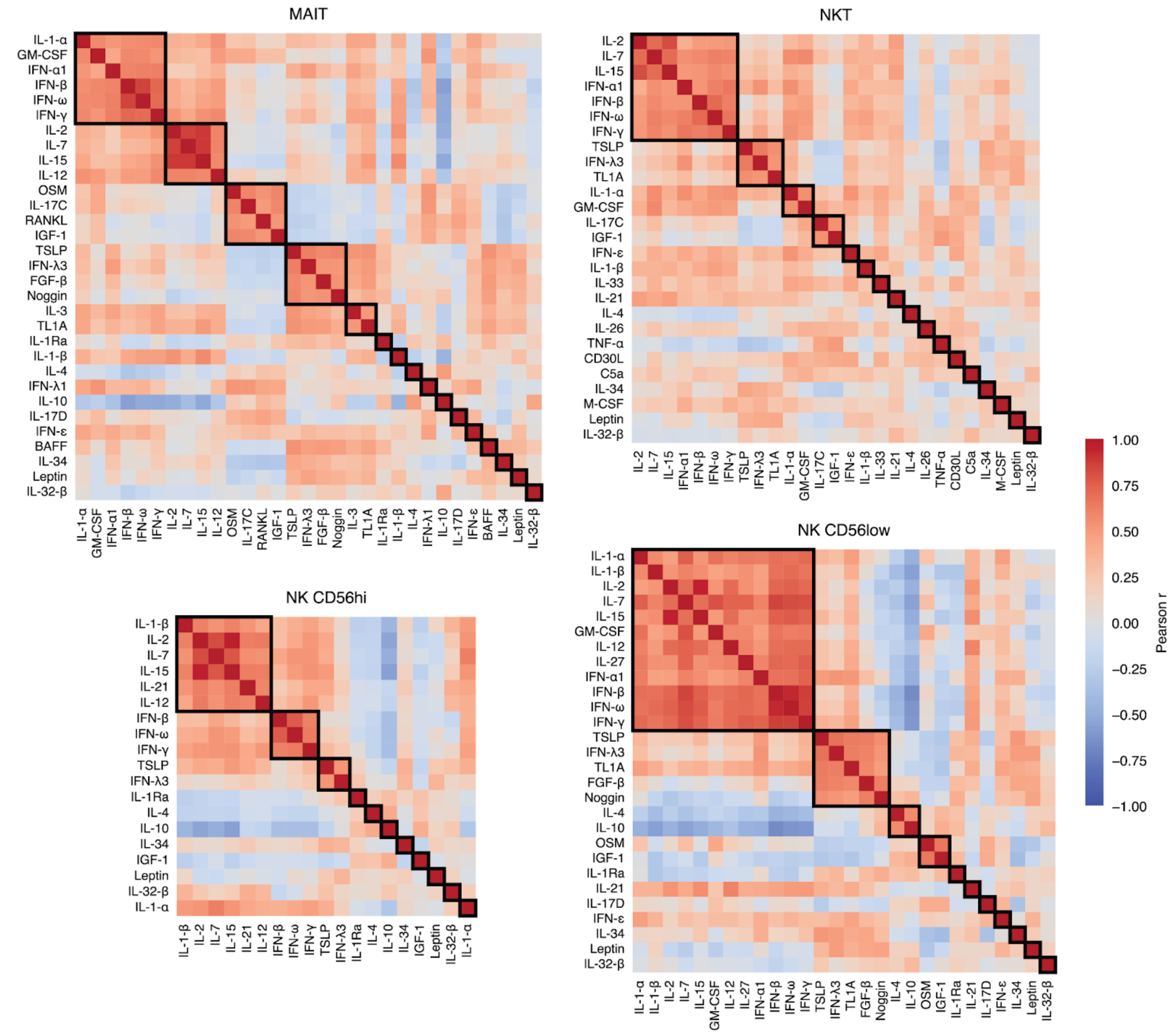


**Fig. S26. Correlation heatmaps between log2FC values for different cytokines in MAIT, NK CD56hi and CD56low, and NKT cells.** Cytokines were grouped based on their correlation patterns using the Leiden algorithm (black boxes).

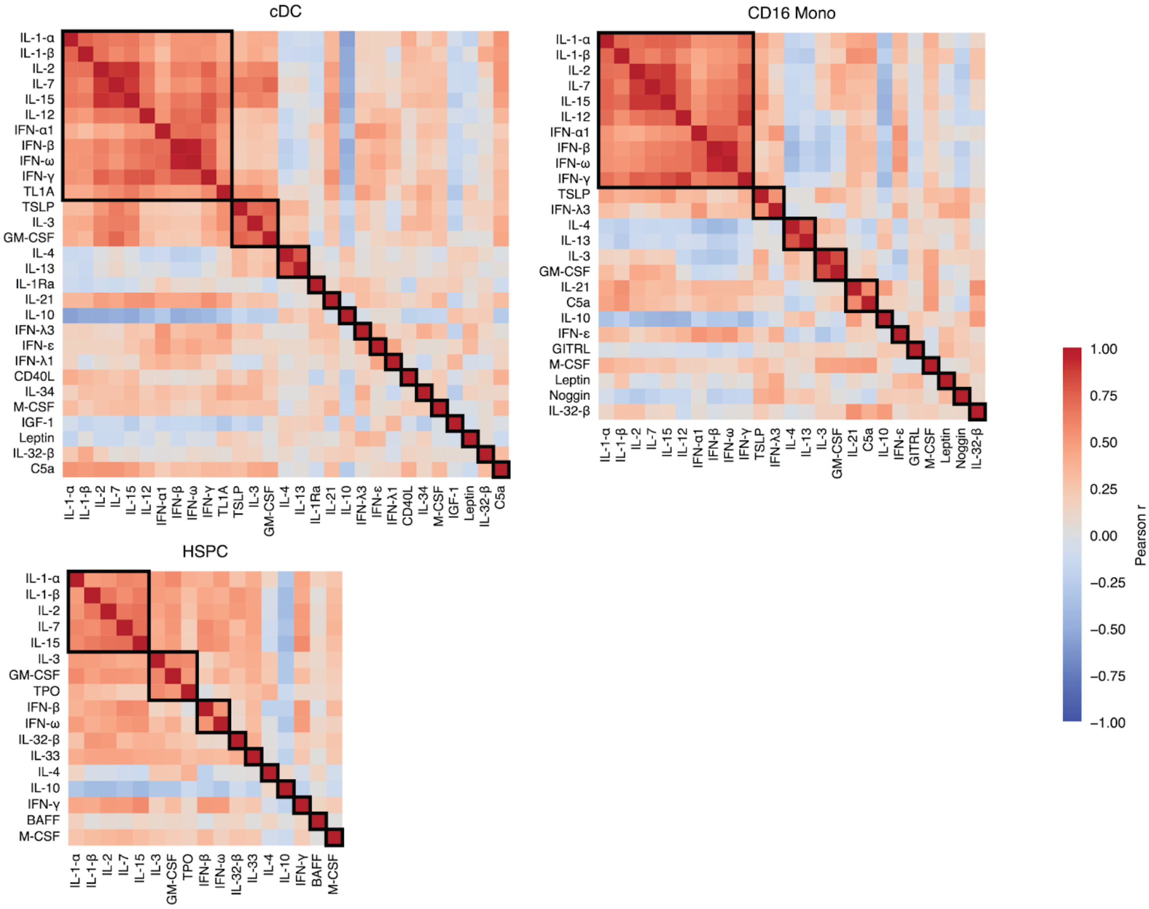


**Fig. S27. Correlation heatmaps between log2FC values for different cytokines in cDCs, CD16 Monocytes, and HSPCs.** Cytokines were grouped based on their correlation patterns using the Leiden algorithm (black boxes).

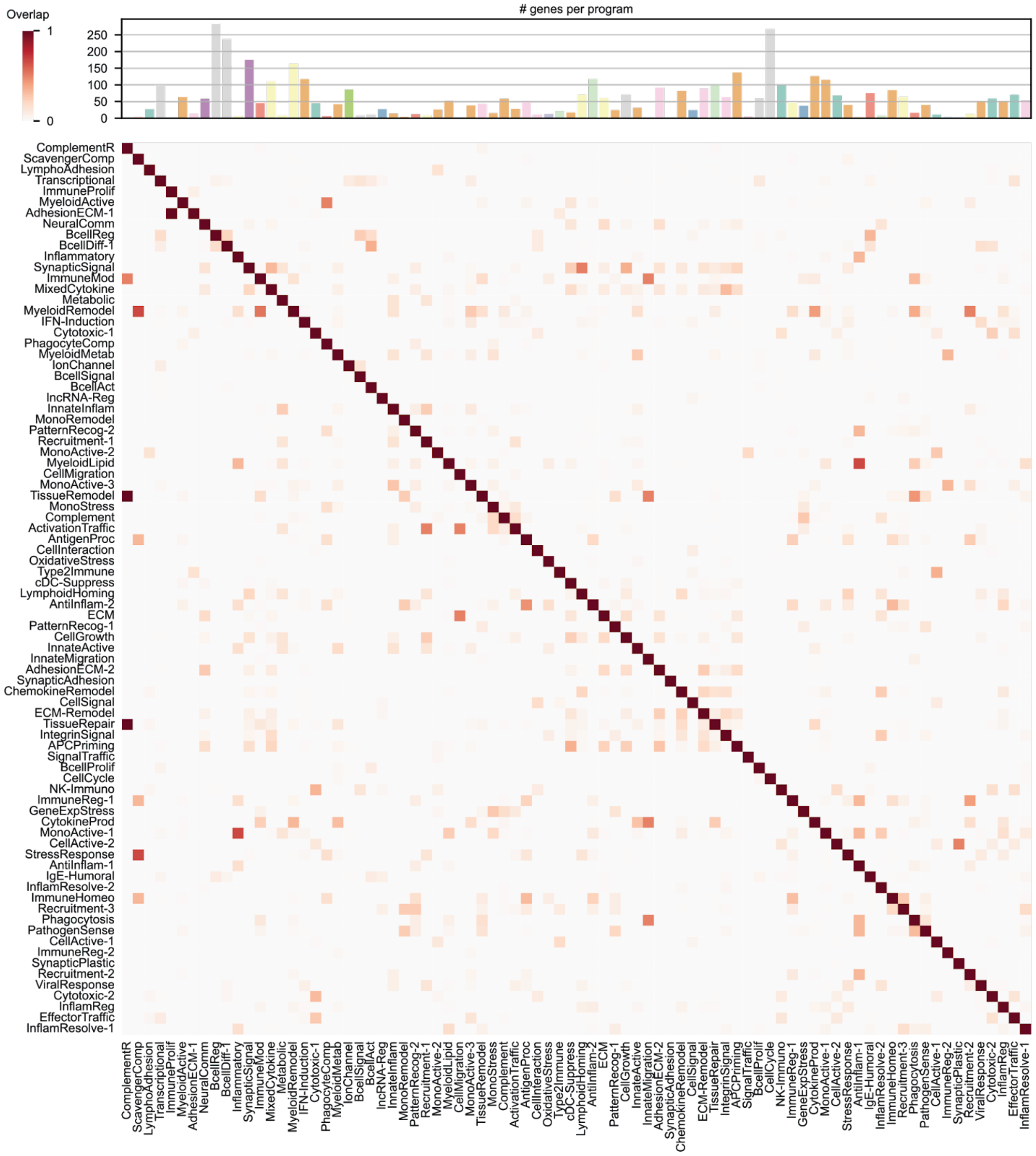


**Fig. S28. Overlap of CIP-associated genes.** The barplot above shows the total number of genes per program. The main plot shows that there is little overlap in genes between programs.

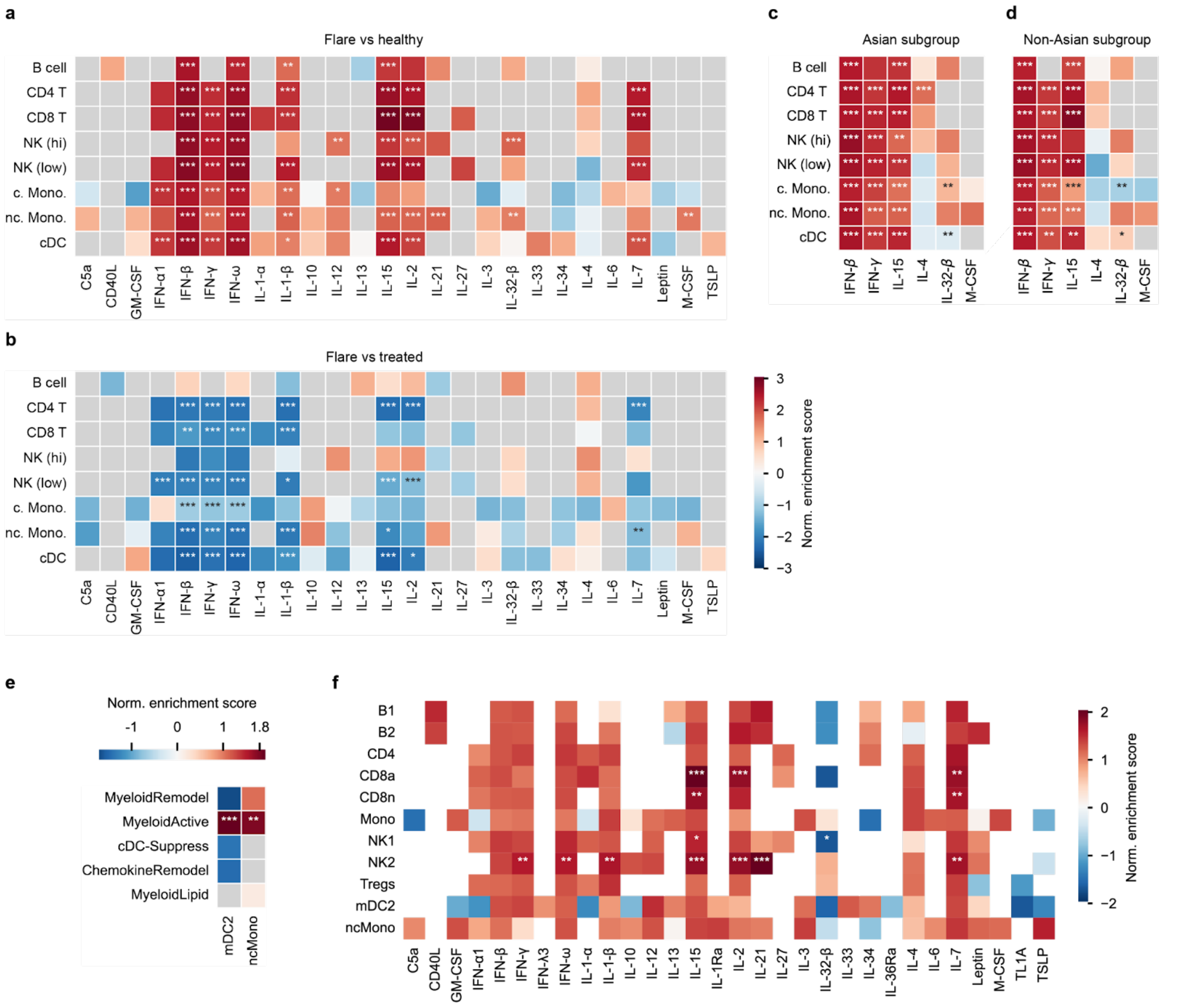


**Fig. S29. Additional results of the enrichment analysis. a,** Enrichment results for all tested cytokines when comparing *Flare* group vs. Healthy group. **b,** Enrichment results for all tested cytokines when comparing Flare group vs. treated group. **c,** Enrichment results for the Asian subgroup, when comparing Flare vs. Healthy. **d,** Enrichment results for the Asian subgroup, when comparing *Flare* vs. *Healthy*. Results in (c) and (d) do not show qualitative differences between both subgroups. **e**, CIP enrichment results for the myeloid compartment, comparing MS vs healthy. **f**, Enrichment results for all tested cytokines when comparing MS vs healthy.
